# Hippocampal CA3 Nex/Neurod6^+^ neuron-specific TNFR2 alleviates chronic neuropathic pain by sex-dependently engaging opioid and endocannabinoid pathways

**DOI:** 10.1101/2025.11.24.690195

**Authors:** Sreejita Arnab, Payam Fathi, Erin Jones, Aravind Meyyappan, Kayla L Nguyen, Valerie Bracchi-Ricard, Shruti Gupta, Roman Fischer, Veronica J. Tom, John R. Bethea

**Affiliations:** Department of Anatomy and Cell Biology, The George Washington University, Ross Hall, Washington, D.C. 20052, USA; Department of Neurobiology & Anatomy, Drexel University College of Medicine, Philadelphia, PA 19129, USA; Center for Immunity and Immunotherapies, Seattle Children’s Research Institute, Seattle, WA 98101, USA; Institute of Cell Biology and Immunology, University of Stuttgart, Stuttgart, Germany

**Keywords:** Chronic neuropathic pain, Chronic constriction injury, Tumor necrosis factor receptor 2, Nex^+^ (Neurod6) neurons, hippocampus, Proopiomelanocortin, Beta-endorphin (β-endorphin), Mu-opioid receptor, Naltrexone, Oleoyl ACP hydrolase, endocannabinoid system, CB_1_R, AM251

## Abstract

Chronic neuropathic pain (CNP) develops as a result of persistent neuroinflammation and maladaptive synaptic plasticity in the central nervous system following nerve injury. While tumor necrosis factor receptor 2 (TNFR2) signaling has been extensively studied in pain resolution, the expression of this receptor on specific neuronal populations and molecular pathways involved in spontaneous pain recovery still remains poorly defined. In this study, we investigated the role of TNFR2 signaling within hippocampal Nex/Neurod6⁺ pyramidal neurons in promoting recovery from chronic constriction injury (CCI), a well-established rodent model of neuropathic pain. To achieve neuron-specific deletion of TNFR2, we generated tamoxifen-inducible conditional knockout mice (NexCre^ERT^^2^:TNFR2^F/F^). We demonstrate that knocking out TNFR2 from Nex⁺ neurons prevents spontaneous pain recovery in both males and females. Thus, establishing that a supraspinal TNFR2 neuroimmune axis is necessary for pain recovery. Exogenous administration of a TNFR2 agonist at 7, 10, and 13 dpi (i.p.) significantly improved mechanical withdrawal thresholds in both sexes of wild-type mice but did not alleviate pain in Nex-specific TNFR2 knockouts, indicating that neuronal TNFR2 expression is required for TNFR2-mediated analgesia. Bulk RNA sequencing of hippocampal tissue collected at six weeks after CCI revealed that TNFR2 activation upregulates genes such as *Pomc,* involved in the opioid pathway, and oleoyl-ACP-hydrolase (OLAH), involved in the endocannabinoid pathway. Consistent with these findings, immunostaining and Western blot analyses showed that TNFR2 agonism restored cornu ammonis (CA3) region POMC and β-endorphin protein levels that were otherwise suppressed after CCI. Behavioral experiment demonstrated that systemic blockade of the µ-opioid receptor with naltrexone (administered daily from 7-21 dpi (s.c.)) completely prevented TNFR2-mediated pain recovery in males but only partially in females. In contrast, inhibition of cannabinoid 1 receptor (CB1R) signaling with AM251 (administered at 7, 14, and 21 dpi (i.p.)) abolished TNFR2-driven analgesia in both sexes. Together, these results reveal that hippocampal TNFR2 signaling in Nex/Neurod6⁺ neurons is critical in recovery from chronic neuropathic pain. TNFR2 activation promotes analgesia by engaging endogenous β-endorphin/µ-opioid and endocannabinoid pathways in a sex-dependent manner, establishing TNFR2 agonism as a promising non-addictive therapeutic approach for chronic pain resolution.

**Significance:** Chronic neuropathic pain (CNP) results from persistent neuroimmune signaling and is driven by maladaptive circuit plasticity. Due to the complexity of factors contributing to CNP, it often leaves patients with few treatment options, which, unfortunately, are either temporary or might be addictive. We have characterized a novel supraspinal mechanism through which tumor necrosis factor receptor 2 (TNFR2) signaling, specifically in hippocampal Neurod6/Nex^+^ expressing pyramidal neurons, is necessary for pain recovery following nerve injury. Pharmacological activation of TNFR2 in these neurons alleviates pain by engaging both endogenous opioid and endocannabinoid signaling pathways. We specifically demonstrate that TNFR2 agonism upregulates proopiomelanocortin (POMC) expression and β-endorphin levels in the hippocampus. We further identify that pharmacological inhibition of either the μ-opioid receptor or cannabinoid 1 (CB1) receptor is sufficient to impair the effectiveness of TNFR2 agonist mediated pain resolution. Our findings thus uncover a novel neuroimmune mechanism where the TNFR2 agonist, exogenously activating the pro-resolving TNFR2, mitigates CNP by releasing endogenous pain neuromodulators. Here, we highlight that TNFR2 agonism could serve as a non-addictive therapeutic strategy for the resolution of chronic neuropathic pain.

## Introduction

Somatosensory injury or disorders that promote sustained chronic neuroinflammation resulting in maladaptive synaptic plasticity ultimately drive the development of chronic neuropathic pain (CNP)^1^. In the United States, chronic pain impacts over 50 million individuals (over 20% of the adult population), and a large portion of these cases involve neuropathic pain that interferes with mobility, sleep, daily functioning, and reduced quality of life^2^. CNP is characterized by hyperalgesia, heightened sensitivity to noxious stimuli, and allodynia or pain elicited following what should be a normally innocuous stimulus^3^. The complex and multifactorial nature of CNP presents formidable challenges for therapeutic intervention, and the continuing opioid epidemic underscores the need for additional safe, durable, and effective treatment options^4–6^.

Tumor necrosis factor (TNF) is an important neuroimmune signaling cytokine involved in injury-induced neuroinflammation and tissue repair^7–9^. Elevated TNF levels have been observed in nerve lesion biopsies of humans with CNP as well as in additional peripheral nerve injury models^7^. Membrane-bound transmembrane TNF (tmTNF), a biologically active form with predominantly anti-inflammatory and neuroprotective properties, engages with TNF receptor 2 (TNFR2) to promote immune regulation, tissue regeneration, and neuronal survival^8–10^. While we have previously shown that exogenous TNFR2 activation, using a novel TNFR2-selective agonist, sex independently promotes long-term resolution of neuropathic pain, the specific cell types and downstream signaling pathways that promote pain relief remain uncharacterized^10^.

Evidence from a previous study has shown the role of the hippocampus (Hc) as a supraspinal node that integrates nociception with cognition and affect^11^. Chronic pain induces remodeling of hippocampal network organization, disrupts hippocampal neurogenesis, and impairs synaptic plasticity, leading to pronounced memory deficits and altered pain affect^12–15^. The ventral hippocampus contributes to the affective motivational dimension of pain, aligning with its broader role in emotional processing^16^. Hippocampal pyramidal neurons encode nociceptive information during neuropathic states, suggesting that hippocampal principal cells participate directly in pain representation^17^. Consistent with this, we have also previously demonstrated that Nex, also known as Neurod6, which is expressed in excitatory pyramidal projection neurons, regulates chronic pain signaling in a sex-specific, supraspinal circuit TNFR1/p38α MAPK-dependent manner^18^. Nex/Neurod6 is a helix-loop-helix transcription factor enriched in supraspinal glutamatergic pyramidal projection neurons that connect the neocortex and hippocampus^19^. While we have previously outlined the contributions of Nex/Neurod6 neuron TNFR1 expression to pain chronification, whether signaling by TNFR2 (within hippocampal Nex+ pyramidal neuron populations) can contribute to pain resolution has not been determined.

Endogenous neuro-modulatory systems that shape hippocampal plasticity might also engage with TNFR2-mediated mechanisms controlling synaptic function and plasticity in the hippocampus^20,21^. Neurons within the hippocampus have been previously shown to express the prohormone POMC, and the endogenous opioid neuropeptide β-Endorphin (β-End) is one of a number of post-translationally processed peptides derived from POMC^22^. β-End plays a demonstrated role in attenuating inflammation, specifically limits TNF-α production, and mitigates pain-related behaviors across multiple models^23,24^. Furthermore, stimulation of the hippocampal POMC/MC4R (melanocortin-4 receptor) pathway restores synaptic plasticity in disease contexts and functionally links the CA1 and CA3 regions^25^. Likewise, the endocannabinoid (EC) system exerts anti-nociceptive effects by signaling through cannabinoid receptors-1 (CB1R) and −2 (CB2R)^26^. CB1R is enriched at presynaptic terminals throughout the

CNS and PNS and is activated by N-acyl ethanol amines such as anandamide (AEA) and monoacylglycerols such as 2-arachidonoylglycerol (2-AG)^26^. Of particular relevance to lipid-peptide crosstalk, metabolites derived from oleic acid, which is one of the primary products hydrolyzed by the enzyme oleoyl-ACP hydrolase (OLAH)^27^, including N-oleoyl glycine and oleamide, can act as CB1R agonists^28,29^. Pain engages hippocampal endocannabinoid signaling, where elevation of anandamide diminishes hyperalgesia and reinstates hippocampal LTP, thus consistent with a CB1R-dependent analgesic mechanism^30^. These pathways provide a conceptual framework in which hippocampal opioid and endocannabinoid signaling converge to recalibrate nociceptive processing during recovery; however, the existence of neuro-immune signaling axes that can leverage this to modulate pain recovery responses has not been previously explored. While tmTNF/TNFR2 signaling within the nervous system has been broadly characterized as contributing to immune regulation and neuroprotection, the contributions of TNFR2 signaling on well-defined neuronal populations within the CNS have not been previously characterized^10,31^. In this study, we provide evidence that TNFR2 signaling in well-defined Nex/Neurod6^+^ expressing supraspinal pain circuits influences pain recovery following a chronic constriction injury (CCI) model of chronic neuropathic pain (CNP). Furthermore, we demonstrate that hippocampal Nex/Neurod6 neuronal TNFR2-driven CNP resolution is dependent on canonical opioid- and endocannabinoid-related nociceptive control.

## Results

### Nex neuron specific TNFR2 expression is necessary for spontaneous chronic neuropathic pain recovery

It has been demonstrated that supraspinal excitatory pyramidal neurons play a crucial role in pain resolution^16^. While we have previously shown that TNFR2 is essential for recovery from mechanical allodynia, the CNS specific TNFR2 expressing cellular populations contributing to this effect are not known^2^. We generated Nex neuron specific TNFR2 knockout mice to test whether TNFR2 signaling in supraspinal excitatory pyramidal neurons contributed to the resolution of CNP. Tamoxifen inducible Nex-Cre (NexCre^ERT2^) mice were crossed with TNFR2-floxed mice (TNFR2^F/F^) to generate NexCre^ERT2^:TNFR2^F/F^, allowing for temporally selective control of TNFR2 deletion in cortical and hippocampal pyramidal neurons. RNAscope ISH was used to confirm the presence of basal Nex neuron TNFR2 mRNA expression within the CA3 region of the hippocampus and to demonstrate tamoxifen-induced recombination efficiency of TNFR2 loss 21 days after the final tamoxifen injection, as shown in **Figure 1A-C**. Spontaneous CNP recovery was evaluated in male and female CCI induced TNFR2^F/F^ (control ) mice or tamoxifen recombined NexCre^ERT2^:TNFR2^F/F^ (Nex^TNFR2-KO^) as described in the experimental timeline shown in **Figure 1D**. Recovery following CCI was evaluated over 12 weeks following injury through von-Frey filament testing of mechanical allodynia^10^. We found that NexCre^ERT2^:TNFR2^F/F^ mice were resistant to spontaneous CNP recovery, whereas TNFR2^F/F^ control mice began to recover starting at 6 weeks post-injury, indicating that Nex neuron specific TNFR2 is necessary for recovery from CCI induced CNP (**Figure 1E**).

**Figure 1:**
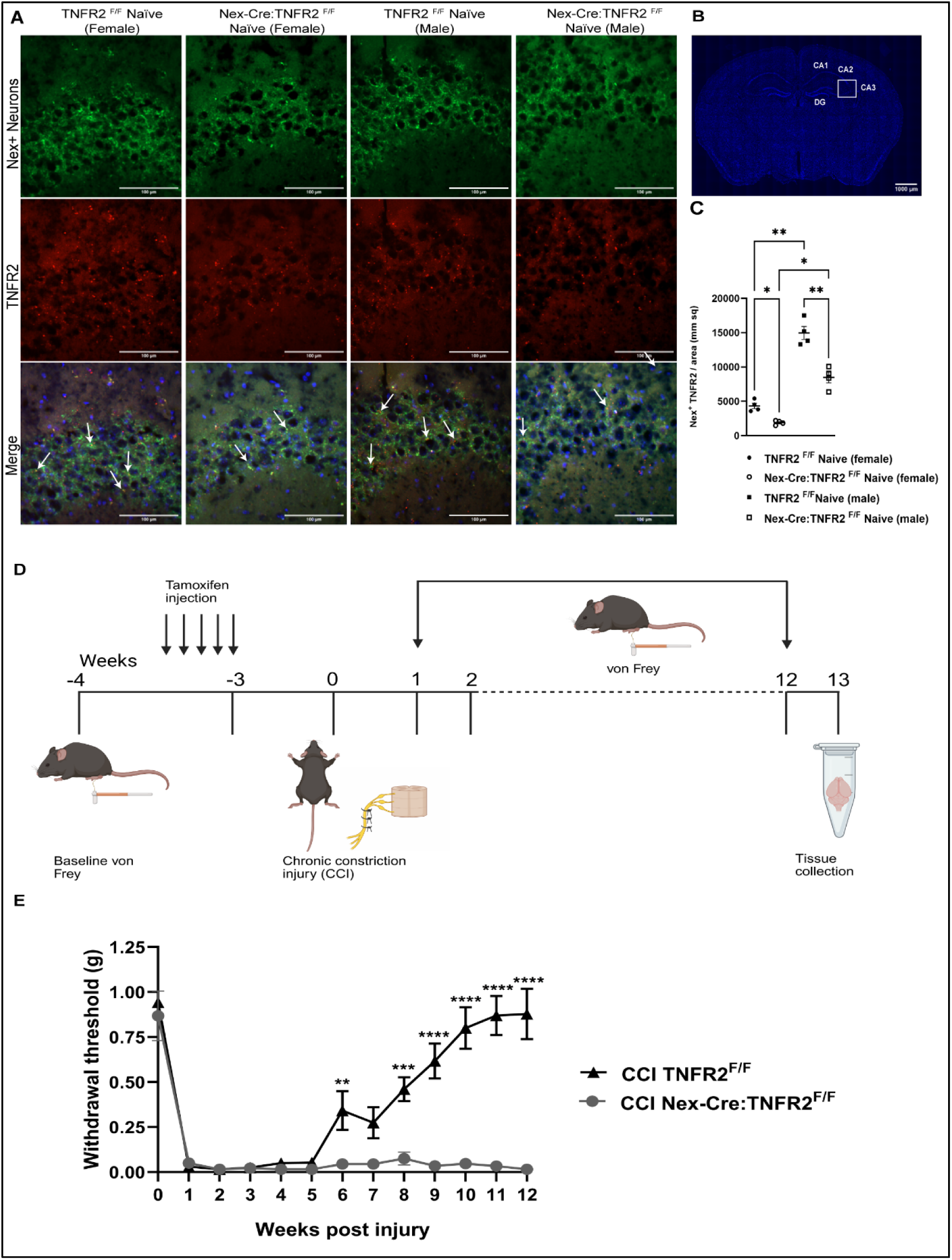
Nex+ TNFR2 signaling is necessary for chronic neuropathic pain recovery. **(A)** mRNA expression of Nex+ neurons co-localized with TNFR2 (marked with arrows) in the hippocampal CA3 region using the RNAscope ISH technique. **(B)** Different regions of the hippocampus, including the CA3 region of the mouse brain. **(C)** TNFR2 KO validation in Nex-Cre:TNFR2^F/F^ mice compared to that of the TNFR2^F/F^ mice in the hippocampal CA3 region (n=4 each group/sex). **(D)** Timeline of the experiment (created in Biorender). **(E)** TNFR2^F/F^ (WT) mice fully recover from CCI, and not the neuronal KO Nex-Cre:TNFR2^F/F^ mice. Withdrawal threshold of TNFR2^F/F^ (WT) and Nex-Cre:TNFR2^F/F^ (KO), both sexes, was measured by von Frey (n=10 each group). Data are represented as mean ± SEM; *\*P* < 0.05, *\*\*P* < 0.01, *\*\*\*P* < 0.001, *\*\*\*\*P* < 0.0001.

### TNFR2 agonist alleviation of chronic neuropathic pain requires Nex neuron TNFR2

Additional work by our group demonstrated that a novel TNFR2 specific agonist is sufficient to reduce both central and peripheral inflammation and promotes injury induced pain recovery in a Treg dependent manner^10^. We expanded upon this by testing whether the observed TNFR2 agonist mediated recovery responses were further dependent on Nex neuron TNFR2 specific expression. Following tamoxifen induced recombination, male and female NexCre^ERT2^:TNFR2^F/F^ or control TNFR2^F/F^ mice were administered 3 doses of either vehicle (PBS) or TNFR2-agonist on days post-injury (dpi) 7,10, and 13 as shown in the experimental timeline (**Figure 2A**). Agonist-treated CCI TNFR2^F/F^ control mice experienced significant progressive recovery in mechanical allodynia across the five-week study timeline, independent of sex, as shown in **Figure 2B-C**. Contrasting this, NexCre^ERT2^:TNFR2^F/F^ completely abolished TNFR2 agonist efficacy with no observed recovery in mechanical allodynia, similar to what is observed in vehicle treated mice (**Figure 2B-C**). These findings corroborate what we demonstrated in **Figure 1** to further confirm the necessary role of TNFR2 expression specifically in Nex neurons for CNP recovery.

**Figure 2:**
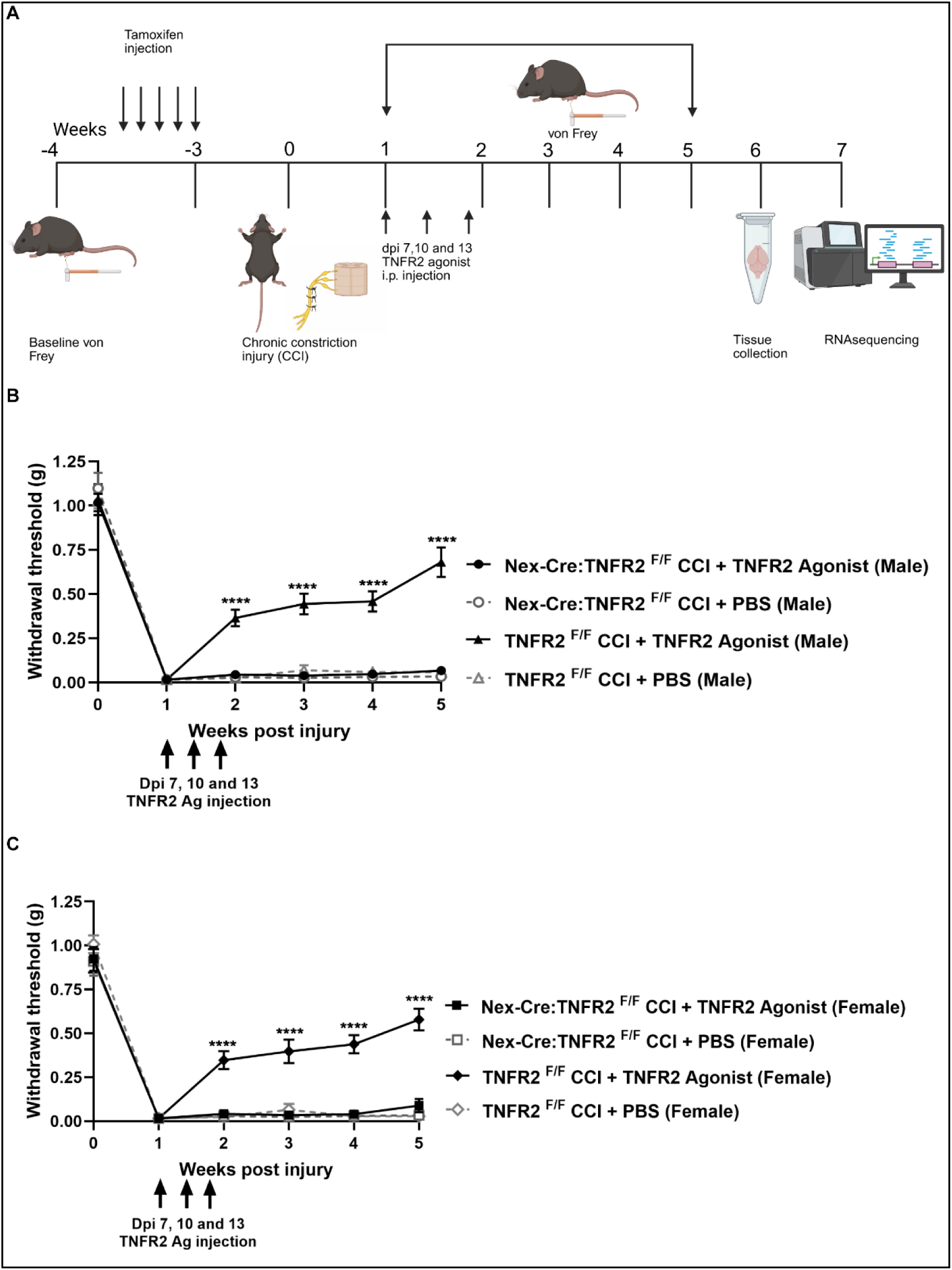
TNFR2 agonism alleviates chronic neuropathic pain in the TNFR2^F/F^ (WT) group but not in the neuronal KO, Nex-Cre:TNFR2^F/F^ group. **(A)** Timeline of the experiment (created in Biorender). After CCI, at Dpi 7,10, and 13, mice were administered (i.p.) either with TNFR2 Agonist (10mg/kg) or PBS. **(B)** and **(C)** Withdrawal threshold of male and female measured by von Frey assessment (n=15 each group/sex). Data are represented as mean ± SEM; *\*\*\*\*P* < 0.0001.

### Hippocampal tissue bulk RNA sequencing implicates TNFR2 agonist engagement of canonical nociceptive control pathways

Expanding on our findings, which showed that Nex neuron TNFR2 expression is necessary for TNFR2 agonist mediated recovery from CCI induced CNP, we performed bulk RNA-sequencing of hippocampal tissue at the endpoint of our previous experiment, detailed in **Figure 2A**, 6 weeks following CCI induction. We specifically focused on transcriptional changes in the hippocampus of male mice since our previous studies did not find any differences between sexes with respect to Nex neuron specific TNFR2 expression on either spontaneous CNP recovery or TNFR2 agonist treatment.

To further resolve changes in gene expression that may be more specifically implicated in Nex TNFR2 specific pain resolution outcomes, we first aimed to identify changes in reciprocally expressed genes between male TNFR2 agonist treated TNFR2^F/F^ control and NexCre^ERT2^:TNFR2^F/F^ mice. We first evaluated gene expression changes in vehicle and agonist treated control TNFR2^F/F^ mice to identify the influence of agonist treatment on gene expression changes following injury only. Next, we evaluated the specific influence of Nex neuron TNFR2 on agonist induced gene expression following CCI by comparing agonist treated TNFR2^F/F^ and NexCre^ERT2^:TNFR2^F/F^. In order to more specifically isolate differences in gene expression that may be more specifically influenced by Nex neuron TNFR2, we evaluated statistically significant enriched reciprocally expressed genes between the two comparison groups, as shown in **Figure 3A**. This approach was used to specifically interrogate whether the specific loss of TNFR2 in Nex neurons would have an influence on genes previously shown to be involved in neuroinflammatory pathways and endogenous opioid or endocannabinoid signaling pathways.

**Figure 3:**
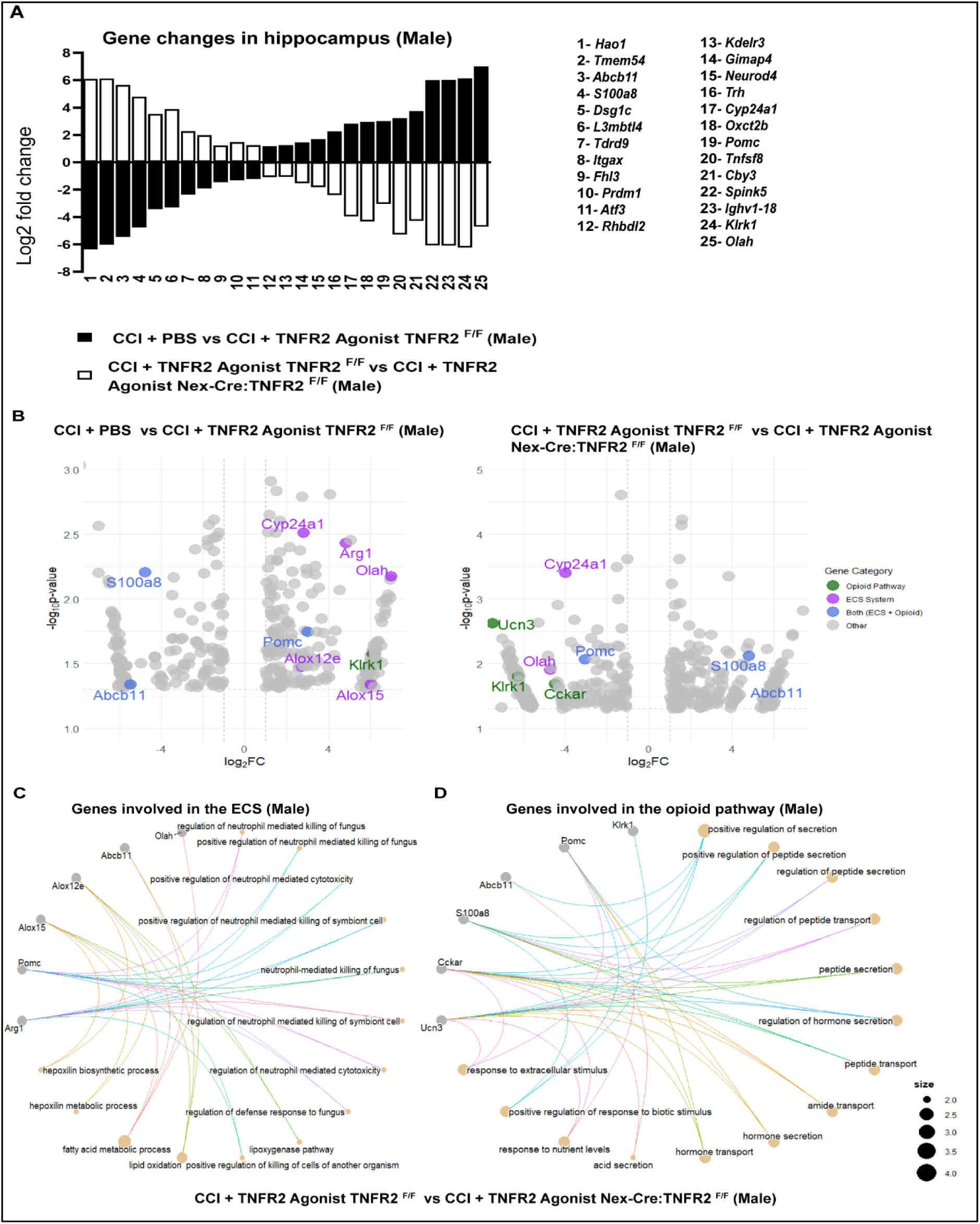
Hippocampal bulk RNA sequencing data reveal endocannabinoid and opioid transcriptional programs in male mice. **(A)** Differential expression of genes in male hippocampus, comparing TNFR2^F/F^ and Nex-Cre:TNFR2^F/F^ mice treated with TNFR2 Agonist.Genes involved in both pathways were reciprocally regulated in TNFR2^F/F^ and Nex-Cre:TNFR2^F/F^ mice treated with TNFR2 Agonist. **(B)** Transcriptome-wide volcano plots of CCI + PBS vs CCI + TNFR2 Agonist of TNFR2^F/F^ mice (left) and CCI + TNFR2 Agonist TNFR2 ^F/F^ vs CCI + TNFR2 Agonist Nex-Cre:TNFR2^F/F^ mice (right) show upregulation and downregulation of genes involved in the ECS and opioid pathway (*P value* < 0.05 and |log2 fold change| > 1). GO pathway enrichment of biological processes show involvement of genes in the **(C)** hormone secretion, transport and regulation, **(D)** fatty acid metabolic process, lipid oxidation and regulation of neutrophil mediated cytotoxicity. (n=4-5 each group)

We identified several reciprocally expressed genes specifically related to neuroinflammation and immune regulation, with specific downregulation of *S100a8* and *Itgax* in TNFR2 receptor expressing CCI mice following agonist treatment^32–34^. In addition, we observed noted changes in genes involved in lipid handling, most notably through an increase in *Olah*, providing a link to TNFR2 agonist mediated pain resolution that may utilize downstream engagement of the endocannabinoid system. Most notably, however, we found stark changes in the reciprocal expression of *Pomc* following TNFR2 agonist treatment, where *Pomc* expression is significantly upregulated in TNFR2 expressing CCI mice and significantly downregulated in mice lacking TNFR2 in Nex neurons (**Figure 3A**). Volcano plots were used to further evaluate differential gene expression of genes that may be either directly or indirectly involved in engaging endogenous opioid or endocannabinoid signaling pathways in **Figure 3B**. Volcano plots comparing differential gene expression across all experimental conditions are shown in **Supplemental Figures 1**. Gene ontology analyses of endocannabinoid or endogenous opioid signaling-related genes revealed enrichment for secretory and peptide handling processes, responses to extracellular stimuli, as well as immune-effector terms **(Figure 3C-D)**. Taken together, our data suggest TNFR2 agonism may reorganize neuroinflammatory, endocannabinoid, and opioid transcriptional programs in a Nex neuron TNFR2 specific manner in the hippocampus of male mice in order to facilitate pain resolution.

### Nex neuron TNFR2 expression mediates TNFR2 agonist induced increases in CA3 POMC and **β**-endorphin protein expression

Prior studies show that POMC helps alleviate synaptic plasticity impairment in neurodegenerative disease models^25,35^. Since our transcriptomics analyses identified *Pomc* as a significant, reciprocally expressed gene following TNFR2 agonist treatment, we validated this in our model by evaluating POMC protein expression within the hippocampal CA3 region following CCI and TNFR2 agonism. POMC protein levels are significantly reduced in CCI-induced TNFR2^F/F^ control male mice relative to the TNFR2 agonist-treated group; thus, TNFR2 agonist treatment was sufficient to significantly restore POMC protein expression (**Figure 4A**). NexCre^ERT2^:TNFR2^F/F^ male mice exhibited slight but not statistically significant decreases in basal POMC expression (**Supplemental Figure 2A)**. Interestingly, NexCre^ERT2^:TNFR2^F/F^ male CCI mice showed a similar drop in POMC protein expression following injury, and TNFR2 agonism was unable to restore POMC expression, providing further evidence that the pain-resolving effects of TNFR2 agonism in male mice may be POMC dependent (**Figure 4A**). In contrast to male mice, female TNFR2^F/F^ and NexCre^ERT2^:TNFR2^F/F^ did not experience any changes in POMC protein expression following CCI (**Figure 4B** and **Supplemental Figure 2B**). Relatedly, TNFR2 agonism following CCI significantly increased CA3 POMC protein expression in female TNFR2^F/F^ but not NexCre^ERT2^:TNFR2^F/F^ mice (**Figure 4B**).

**Figure 4:**
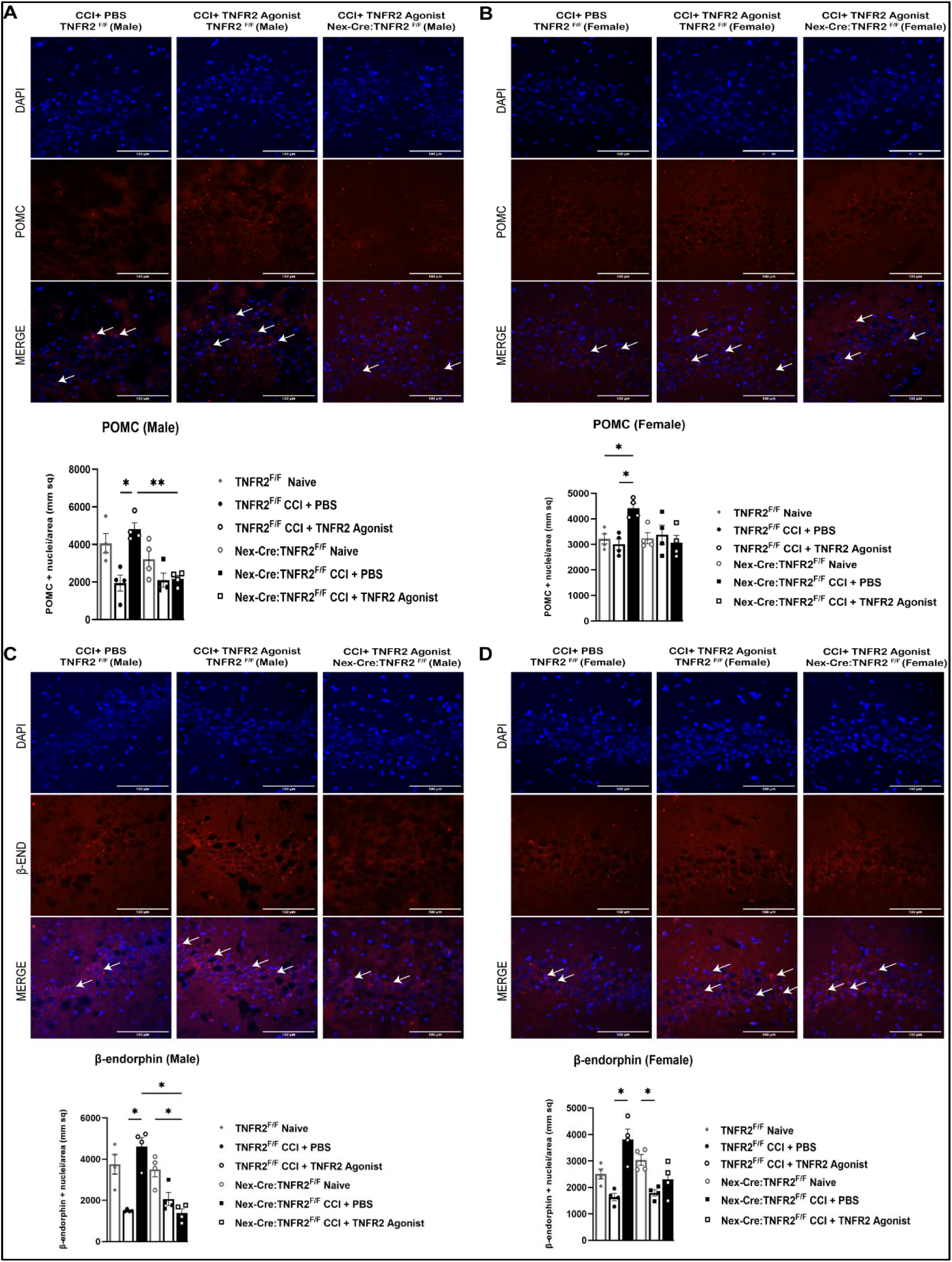
Elevation in protein expression levels of POMC and β-endorphin in the CA3 region of the hippocampus in TNFR2^F/F^ (WT) mice treated with TNFR2 Agonist compared to that of the neuronal KO, Nex-Cre:TNFR2^F/F^ mice. **(A)** and **(B)** POMC expression levels in the hippocampal CA3 region of male and female, represented by immunohistochemistry (n=4 each group/sex). **(C)** and **(D)** β-Endorphin expression in the hippocampal CA3 region of male and female, represented by immunohistochemistry (n=4 each group/sex). Data are represented as mean ± SEM; *\*P* < 0.05, *\*\*P* < 0.01.

Given that the endogenous opioid β-endorphin is a β-lipotropic hormone facilitated post-transcriptional cleavage product of POMC^36^, we tested whether the TNFR2 agonist mediated changes in POMC protein expression would facilitate similar increases in β-endorphin within the CA3. Our data show a similar trend in male mice, where CCI prompts a significant decrease in β-endorphin male TNFR2^F/F^ that is then significantly restored following TNFR2 agonist treatment (**Figure 4C**). This effect was found to be Nex-neuron TNFR2 specific since TNFR2 agonist treatment in CCI induced NexCre^ERT2^:TNFR2^F/F^ male mice did not change the injury induced decreases in β-endorphin protein levels (**Figure 4C** and **Supplemental Figure 2C**). Female mice showed slight decreases in β-endorphin levels following CCI injury, but these decreases were only statistically significant in the NexCre^ERT2^:TNFR2^F/F^ group (**Figure 4D** and **Supplemental Figure 2D**). Similar to what we observe in males, TNFR2 agonist treatment following CCI was able to significantly increase CA3 β-endorphin levels in TNFR2^F/F^ but not NexCre^ERT2^:TNFR2^F/F^ mice (**Figure 4D**). These findings provide additional supportive evidence that suggests that hippocampal Nex neuron specific TNFR2 signaling promotes pain resolution through engagement of endogenous opioid pathways through upregulation of β-endorphin.

### Naltrexone mediated µ-opioid receptor antagonism prevents TNFR2-agonist mediated pain recovery in male but not female mice

To directly test whether TNFR2 in Nex neuron mediates pain recovery by engaging endogenous opioid signaling pathways through β-endorphin, we administered the µ-opioid receptor antagonist Naltrexone following CCI and TNFR2 agonist treatment in TNFR2^F/F^ mice, since β-endorphin primarily signals through the µ-opioid receptor^37,38^. Following CCI, male and female mice were administered vehicle, TNFR2 agonist, or TNFR2 agonist with Naltrexone (**Figure 5A**). Naltrexone was administered subcutaneously, daily starting at dpi 7, concurrent with the first TNFR2 agonist dose and continued through dpi 21. Assessments of pain recovery, as measured through Von-Frey testing of mechanical allodynia, found that naltrexone treatment completely abolished TNFR2 agonist efficacy in male mice, and only partially blunted recovery in female mice (**Figure 5B-C**). These findings suggest that while there are similar TNFR2 agonist mediated changes in β-endorphin expression within the CA3, pain recovery following CCI in male, but not female mice, require engagement of opioid signaling pathways.

**Figure 5:**
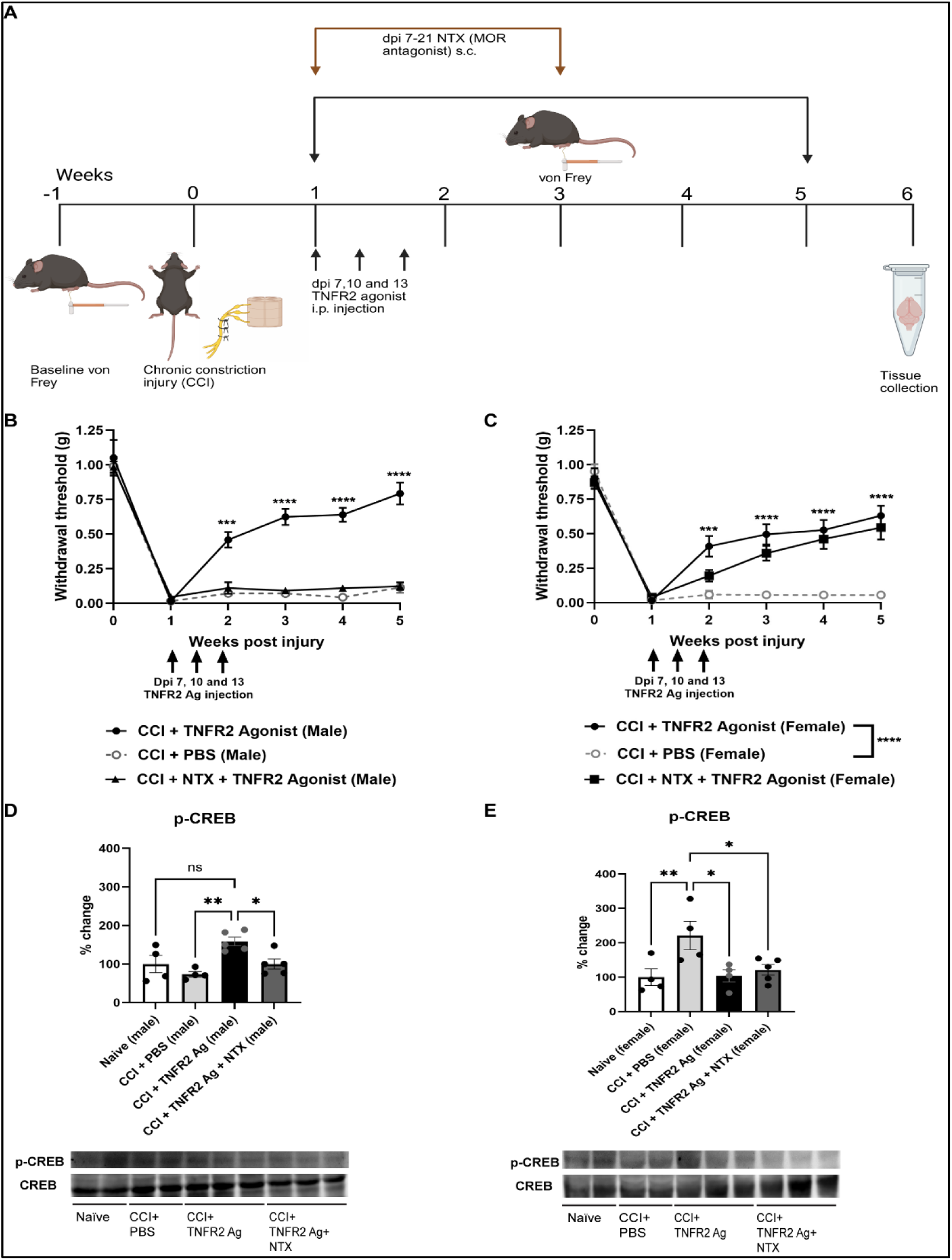
MOR antagonist (Naltrexone) shows sexual dimorphism in CNP recovery when treated with TNFR2 agonist **(A)** Timeline of the experiment (created in Biorender). After CCI, at dpi 7, 10, and 13, mice were administered (i.p.) either with TNFR2 Agonist (10mg/kg) or PBS, and the μ-opioid receptor antagonist Naltrexone (NTX) was delivered s.c. from dpi 7-21. Longitudinal mechanical sensitivity (von Frey withdrawal thresholds) following CCI in males that do not recover from CNP **(B)** and females show similar recovery when treated with TNFR2 Agonist **(C)** (n=10 each group/sex). Representative hippocampal immunoblots for phospho-CREB (p-CREB) from **(D)** males (n=4-5 each group); p-CREB levels were elevated after TNFR2 agonism and reduced after CCI and NTX co-treatment with TNFR2 agonist, and **(E)** females (n=4-5 each group); p-CREB levels were elevated after CCI and reduced by TNFR2 agonism and NTX co-treatment in females. Data are represented as mean ± SEM; *\*P* < 0.05, *\*\*P* < 0.01, *\*\*\*P* < 0.001, *\*\*\*\*P* < 0.0001.

To further validate sex-dependent alterations in TNFR2 agonist mediated engagement of opioid signaling on pain recovery, we used western blotting to evaluate changes in hippocampal p-CREB at the study endpoint, 6 weeks post-CCI. CREB (cAMP response element–binding protein) is a kinase-activated transcription factor; when phosphorylated at Ser133, it converts neuronal and inflammatory signals into coordinated changes in gene expression^39,40^. CREB can go up or down depending on whether the system is in a central-sensitization (pro-nociceptive) state^41^ or a pro-resolving (analgesic) state^42,43^. We observed sexual dimorphism in the p-CREB changes across the group. In male mice, p-CREB was significantly upregulated following TNFR2 agonist treatment and was reduced to baseline levels following CCI injury and Naltrexone co-treatment. (**Figure 5D**). Interestingly, in female mice, we observed a stark increase in p-CREB expression following CCI injury that was reduced back to baseline levels following TNFR2 agonist treatment, irrespective of Naltrexone (**Figure 5E**). Together, these data indicate that TNFR2-driven recovery from mechanical allodynia requires μ-opioid signaling with sex-biased magnitude, and that hippocampal p-CREB mirrors the direction of behavioral changes in both male and female mice.

### AM251 mediated CB1R antagonism prevents TNFR2 agonist mediated pain recovery in both male and female mice

While we were able to directly show that Naltrexone mediated µ-opioid receptor antagonism was sufficient to abolish TNFR2 agonist mediated pain recovery in male but not female mice, we next tested whether female mice may preferentially engage endocannabinoid signaling pathways to promote pain recovery. AM251 antagonism of the CB1-receptor was conducted by weekly injections starting at dpi 7, concurrent with the first dose of TNFR2 agonist (**Figure 6A**). Following CCI, as expected, systemic TNFR2 agonism increased von Frey withdrawal thresholds in both males and females (**Figure 6B-C**). However, AM251 antagonism was sufficient to completely abolish TNFR2 agonist mediated pain recovery in both male and female mice (**Figure 6B-C**).

**Figure 6:**
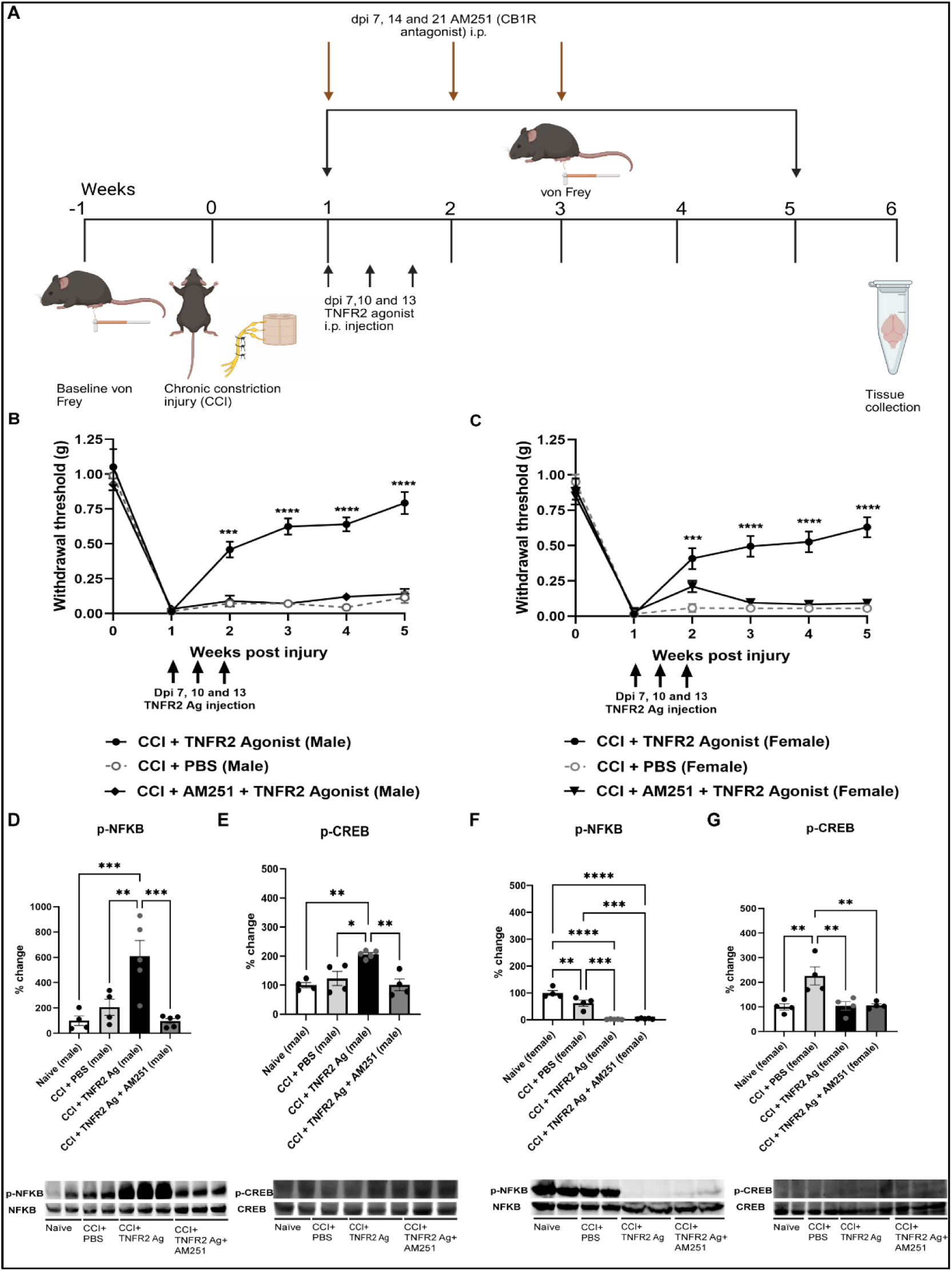
CB1 receptor blockade using AM251 prevents TNFR2 Agonist-driven recovery after CCI in both sexes and reveals sex-divergent hippocampal NF-κB/CREB signaling. **(A)** Timeline of the experiment (created in Biorender). After CCI, at dpi 7, 10, and 13, mice were administered (i.p.) either with TNFR2 Agonist (10mg/kg) or PBS, and the CB1 receptor antagonist AM251 (i.p.) was given at dpi 7, 14, and 21. Longitudinal von Frey withdrawal thresholds after CCI in males **(B)** and females **(C)** (n=10 each group/sex). (D-E) Hippocampal western blots and quantification for males: phospho-NF-κB **(D)** and phospho-CREB **(E)**. In males, TNFR2 agonism elevated p-p65 and p-CREB relative to Naïve and CCI+PBS; AM251 reduced both toward or below baseline (n=4-5 each group). (F-G) Hippocampal western blots and quantification for females phospho-NF-κB **(F)** and p-CREB **(G)**. In females, TNFR2 agonism and AM251 co-treatment reduced p-p65 relative to Naïve and CCI+PBS,; p-CREB levels were elevated after CCI and reduced by TNFR2 agonism and AM251 co-treatment (n=4-5 each group). Data are represented as mean ± SEM; *P<0.05, **P<0.01, ***P<0.001, ****P<0.0001.

Endpoint, 6-week post CCI, western blotting of hippocampal tissues found a significant increase in p-NFκB following TNFR2 agonist treatment that is significantly attenuated by AM251 treatment (**Figure 6D**). Similarly, we observed a significant increase in p-CREB following TNFR2 agonist treatment, which is likewise significantly decreased following AM251 antagonist administration (**Figure 6E**). Interestingly, in female mice, we find that CCI results in slight but significant decreases in p-NFκB levels, that are further decreased following TNFR2 agonist treatment with or without AM251. P-CREB was increased significantly following CCI injury, which was reduced back to baseline levels following TNFR2 agonist treatment, irrespective of AM251 (**Figure 6F**). Together, these data indicate that CB1 receptor signaling is necessary for the behavioral benefits of TNFR2 agonism after nerve injury and that hippocampal NFκB/CREB pathways are engaged in a sex-divergent manner, enhanced in males and suppressed in females, providing a mechanistic correlate of the CB1 dependence observed behaviorally.

In this study, we show that TNFR2 signaling in (1) hippocampal CA3 Nex/Neurod6^+^ pyramidal neurons is required for recovery from CCI-induced chronic neuropathic pain; genetic deletion of TNFR2 in these neurons prevents spontaneous CNP recovery, (2) whereas systemic TNFR2 agonism restores mechanical thresholds in wild-type but not NexCre^ERT2^:TNFR2^F/F^ mice. (3) Hippocampal transcriptomics and immunostaining identify involvement of endogenous analgesic systems; upregulation of *Pomc* with increased CA3 POMC/β-endorphin protein, and induction of lipid-handling genes (e.g., *Olah/Alox15*) consistent with endocannabinoid pathway activation. (4) Mechanistically, naltrexone reveals a sexual dimorphism in CNP recovery; μ-opioid signaling (blocked by naltrexone) is required for TNFR2-driven recovery in males but not females, while CB1 receptor antagonism (AM251) prevents TNFR2-mediated analgesia in both sexes. (5) Interestingly, at the supraspinal level, signaling also diverges by sex, with TNFR2 agonism elevating hippocampal p-CREB and NF-κB programs in males, but reducing injury-evoked p-CREB and NF-κB in females, defining a neuronal TNFR2 axis that resolves pain by differentially recruiting opioid and endocannabinoid pathways.

## Discussion

### Nex neuron specific TNFR2 expression is necessary for the spontaneous recovery of chronic neuropathic pain

Here, we have demonstrated that Nex^+^ (Neurod6) hippocampal neuron TNFR2 signaling is required for the spontaneous resolution of CCI-induced CNP. Our data showed that loss of TNFR2 expression in excitatory Nex neurons resulted in the persistence of pain across the study period, as demonstrated by lack of improvement in mechanical allodynia. This aligns with previous work from our group highlighting TNFR2 expression as a key mediator of long-term pain resolution^10^. We previously showed in global TNFR2 knockout mice that loss of TNFR2 results in chronic, non-resolving pain following nerve injury, whereas wild-type TNFR2 expressing mice experience spontaneous pain recovery^10^. Likewise, prior work by our group has also shown that inhibition of TNFR1 signaling, sex-dependently, mitigates the development and maintenance of neuropathic pain specifically in male mice^44^. The findings of these studies, when taken together, further highlight a functional dichotomy between the two TNF receptors: signaling through TNFR1 drives pro-nociceptive programs, and signaling through TNFR2 promotes engagement of endogenous analgesic programs to facilitate spontaneous pain recovery. Our work builds on this by identifying hippocampal Nex expressing neurons as the specific TNFR2 expressing population within the CNS driving TNFR2 mediated spontaneous neuropathic pain recovery following nerve injury.

We have also previously demonstrated that supraspinal Nex neuronal TNFR1/p38αMAPK signaling is critical to pain chronification in a sex-specific manner^18^. Specifically, we found that loss of TNFR1 in Nex neurons prevented the onset of chronic pain in male mice and facilitated recovery from mechanical allodynia following nerve injury. Building on this, we show that TNFR2 expression in this same population of supraspinal neurons is necessary for spontaneous pain recovery at later timepoints following injury. Our results provide further evidence for a temporally resolved preferential engagement of TNFR signaling during the chronification and spontaneous recovery from injury induced neuropathic pain, where, at least in male mice, early post-injury Nex dependent TNFR1 signaling facilitates initiation of injury induced chronic pain. This is then followed by Nex dependent TNFR2 signaling at later timepoints that promotes the spontaneous resolution of injury induced CNP. The CA3 region of the hippocampus is densely populated with excitatory pyramidal neurons, which might serve as a hub where pro-inflammatory signals (such as soluble TNF in chronic pain) are counter-balanced by TNFR2-driven protective pathways. As we have shown in our studies, the absence of the Nex neuronal TNFR2 may serve to disrupt this balance, thereby providing a shift towards persistent pain hypersensitivity. Our findings also show that both male and female mice require intact TNFR2 expression to spontaneously recover from pain, which is consistent with the previous work on TNFR2-dependent mechanisms of pain resolution in both sexes^10^. Prior work from our group has also shown that estrogen can prolong neuropathic pain in females by enhancing TNF/TNFR1 signaling^18^. Thus, activating TNFR2 may help counteract this bias and promote recovery in both sexes.

### Nex neuron TNFR2 expression is necessary for TNFR2 agonist mediated chronic neuropathic pain resolution

We found that pharmacologically mediated TNFR2 specific agonism markedly alleviated neuropathic pain only when TNFR2 expression in Nex⁺ neurons was preserved. TNFR2^F/F^ mice treated with a TNFR2 agonist showed significant reversal of mechanical allodynia, whereas Nex specific Cre driven TNFR2 loss prevented agonist mediated therapeutic benefits. This result provides strong evidence that the therapeutic effect of TNFR2 agonism is mediated specifically through Nex^+^ neuronal TNFR2 signaling.

Previous work has demonstrated that systemically administered TNFR2 agonist mediated neuropathic pain recovery was dependent on T-reg TNFR2 expression and subsequent suppression of inflammation^10,45^. This suggests a dual mode of action for TNFR2 agonism: one enhancing Treg function to resolve inflammation peripherally, and another, acting within CNS pain circuits through neuronal TNFR2. Consistent with this, a study in CNS autoimmunity models has demonstrated that exogenous TNFR2 activation simultaneously reduces inflammation and promotes neuronal recovery^45^. Our present findings provide further confirmation, specifically showing that supraspinal neuron TNFR2 expression is essential for pain relief.

These data are encouraging from a therapeutic perspective. TNFR2 agonism engages endogenous analgesic networks within the brain to promote lasting recovery from chronic neuropathic pain, independent of exogenous opioids. The efficacy observed in both male and female wild-type mice suggests broad therapeutic potential across sexes, while the lack of recovery in knockout animals confirms target specificity. By activating Nex^+^ neuronal TNFR2 signaling, TNFR2 agonism may provide a therapeutic strategy for permanent treatment of pain recovery through endogenous analgesia versus temporary pain blockage.

### Hippocampal tissue bulk RNA sequencing implicates TNFR2 agonist engagement of canonical nociceptive control pathways

To understand how TNFR2 signaling in hippocampal neurons facilitates pain resolution, we performed bulk RNA sequencing of hippocampal tissue, which revealed significant transcriptional changes associated with TNFR2 agonist mediated CNP recovery. Agonist treatment of TNFR2 expressing mice following injury uncovered significant alterations in gene expression relative to agonist treatment in mice lacking Nex neuron TNFR2, suggesting neuronal TNFR2-dependent control of pro-analgesic transcriptional programs.

Among these, *Pomc*, encoding the prepropeptide POMC that can be post-translationally processed to release the endogenous opioid β-endorphin, was statistically upregulated in TNFR2^F/F^ mice but not in NexCre^ERT2^:TNFR2^F/F^, suggesting that TNFR2 signaling activates an opioid pathway within the hippocampal Nex TNFR2 expressing cells. This is remarkable, as *Pomc* expression has been more classically characterized within hypothalamic neurons^46^. These findings provided evidence for the existence of a POMC/opioid signaling axis within hippocampal neurons that are specifically engaged following TNFR2 agonism after CCI induced neuropathic pain, suggesting a mechanism through which TNFR2 signaling can modulate endogenous opioid production or release to facilitate pain recovery.

Furthermore we observed potential genes linked to endocannabinoid signaling may also be differentially expressed. TNFR2 activation increased *Olah,* an enzyme that generates oleic acid, a precursor of oleoylethanolamide (OEA)^47,48^. Thus, *Olah* may be indirectly involved in promoting anti-inflammatory and analgesic signaling via PPAR-α (Peroxisome Proliferator-Activated Receptor alpha)^49^. Similarly, *Alox15* (arachidonate 15-lipoxygenase) and its isoform *Alox12e* (arachidonate 12-lipoxygenase, epidermal type) were upregulated, both involved in oxygenating endocannabinoids such as anandamide (AEA) and 2-arachidonoylglycerol (2-AG) to form pro-resolving oxylipins^50–52^. This suggests that TNFR2 agonist mediated signaling may promote endocannabinoid related synthetic pathways and subsequently engage inflammation or pain resolving pathways in our model.

Another gene that showed transcriptional change is *Arg1* (Arginase-1), a marker of anti-inflammatory “M2” microglia^53^. *Arg1* is indirectly linked to the endocannabinoid system (ECS), as microglial CB2 receptor activation promotes Arg1⁺ phenotypes and supports endocannabinoid-mediated analgesia^54,55^. Other relevant upregulated genes included *Abcb11*, a bile acid transporter that can indirectly enhance anandamide synthesis through NAPE-PLD activation^56^. *Cyp24a1* is a vitamin D-metabolizing enzyme potentially linked to cannabinoid-steroid crosstalk^57,58^. Upregulation of *Klrk1* (encoding NKG2D) may reflect immune modulation supporting opioid-driven recovery^59,60^, while *S100a8* downregulation by TNFR2 could balance cannabinoid anti-inflammatory effects and TLR4-linked opioid tolerance^32,61^. *Abcb11* induces TGR5 receptor activation, further validating engagement of opioid-dependent analgesia^62^.

Together, these results indicate that agonist mediated TNFR2 signaling activates two intrinsic analgesic signaling pathways, the endogenous opioid pathway via *Pomc*/β*-endorphin* and the endocannabinoid pathway through several enzymes, including *Olah*. This dual transcriptional activation likely facilitates a coordinated neuroimmune shift toward pain resolution.

### Nex neuron TNFR2 expression mediates TNFR2 agonist induced increases in CA3 POMC and **β**-endorphin protein expression

Consistent with our transcriptomic data, we observed a significant change in the protein levels of both POMC and its peptide product β-endorphin in the CA3 region following TNFR2 activation. In TNFR2^F/F^ mice, TNFR2 agonism following CCI induction produced a significant elevation of both POMC and β-endorphin protein levels. In contrast, Nex specific TNFR2 knockout mice showed little to no change, confirming that hippocampal TNFR2 signaling directly engages the translation of endogenous opioids in the context of CCI.

β-Endorphin is a potent endogenous neuropeptide and MOR agonist and provides a direct mechanism for downstream TNFR2 engaged pain relief^63^. Increases in CA3 β-endorphin imply local synthesis and release by hippocampal neurons, which may act on MORs within hippocampal circuits. Another possibility is that β-endorphin released in the hippocampus could diffuse through cerebrospinal fluid to reach distant brain regions, including the brainstem^64^. In either case, TNFR2-induced β-endorphin engages with supraspinal MOR and reduces pain perception.

The upregulation of β-endorphin by TNFR2 agonism connects our work to classical pain regulatory mechanisms. Immune cells such as T cells and monocytes release β-endorphin to activate opioid receptors on peripheral nerves to ultimately reduce pain^65^. Our findings suggest a parallel central process where activation of hippocampal Nex neuron TNFR2 serves as a central source of β-endorphin production which may be utilized as an immune-neural bridge facilitating Treg-mediated peripheral analgesia^10^ while promoting central neuronal opioid release. Interestingly, β-endorphin has effects beyond analgesia, where MOR activation may also serve to reduce stress and neuroinflammatory responses^66–68^. Expanding on this, it is possible that TNFR2 agonist-induced β-endorphin contributes to a self-reinforcing recovery loop that relieves pain, dampens inflammation, and further inhibits pain chronification. Because POMC is a precursor to several peptides, including ACTH and α-MSH, the observed CA3 upregulation may have broader implications. α-MSH, in particular, exerts anti-inflammatory effects through melanocortin receptors^69^ and could complement β-endorphin-mediated analgesia.

### Naltrexone mediated µ-opioid receptor antagonism prevents TNFR2-agonist driven pain recovery in male but not female mice

Here we uncovered a novel sex-dependent role for MOR signaling in TNFR2-mediated recovery from neuropathic pain. In males, TNFR2 agonism significantly reversed mechanical allodynia, but this effect was blocked by naltrexone mediated MOR antagonism, indicating that TNFR2-driven analgesia in males is specifically dependent on μ-opioid receptor signaling. This observation aligns with prior evidence that TNFR2-expressing immune cells can release opioid peptides such as β-endorphin and enkephalins to suppress pain^70,71^. In contrast, females retained pain recovery despite naltrexone treatment, suggesting that TNFR2 activation in females engages alternative, MOR-independent mechanisms. The persistence of recovery in females implies that males and females engage different neuroimmune mechanisms for pain modulation^72^.

At the molecular level, phosphorylated CREB (p-CREB) patterns in the hippocampus have reflected these sex-specific differences (**Figure 5**). In females, CCI elevated hippocampal p-CREB, which has been described as a known marker of injury-induced plasticity in pain pathways^73^, while TNFR2 agonism normalized p-CREB levels similar to naïve mice, mirroring behavioral recovery. Co-treatment with naltrexone also showed a similar p-CREB level to that of naïve and TNFR2 agonist treated mice, further validating the behavioral data and suggesting that TNFR2 reduces p-CREB through MOR-independent mechanisms.

In males, TNFR2 agonism increased hippocampal p-CREB, and naltrexone administration returned p-CREB levels towards baseline, demonstrating that CREB phosphorylation in males is MOR-dependent. Mechanistically, TNFR2 activation likely stimulates β-endorphin release, which engages hippocampal MORs and triggers MAPK/ERK signaling^74,75^. ERK then activates downstream kinases, including MSK1/2, which phosphorylate CREB at Ser133^76,77^. Phosphorylated CREB can promote expression of plasticity-related genes associated with pain resolution, linking MOR activation to adaptive neural remodeling. It is therefore possible that blocking MOR signaling via naltrexone administration disrupts this cascade, preventing CREB phosphorylation and thereby abolishing TNFR2-driven recovery in males. Together, these findings imply a sexual dimorphism where TNFR2-induced analgesia involves dampening of CREB activity in the female hippocampus, thus reflecting reduced neuronal excitability or stress, while in males, TNFR2 signaling promotes MOR-dependent CREB phosphorylation and expression of plasticity-related genes that facilitate recovery.

### Antagonism of CB1R through AM251 administration prevents TNFR2 agonist mediated pain recovery in both male and female mice

Our data reveal that cannabinoid receptor signaling is essential for TNFR2-mediated analgesia, with distinct downstream mechanisms found in both male and female mice. Systemic administration AM251, a CB1 receptor antagonist, prevented TNFR2 agonist mediated improvement of mechanical allodynia in both sexes. This observation is consistent with the established role of the endocannabinoid system in modulating pain across central and peripheral signals^78–80^. Thus, TNFR2 activation likely engages endogenous cannabinoids to act on CB1R and initiate analgesia, highlighting an important crosstalk between TNFR2 signaling and the endocannabinoid system.

Although CB1R signaling is necessary for behavioral recovery in both male and female mice, the hippocampal signaling downstream of TNFR2 activation was strikingly different. In males, TNFR2 agonism increased phosphorylated NF-κB (p-NF-κB) and CREB (p-CREB), whereas AM251 reduced these levels. These results suggest that TNFR2 activates a CB1R-dependent cascade that elevates NF-κB and CREB activity in male hippocampal neurons. While NF-κB is best known for its pro-inflammatory roles, it can also context specifically mediate neuroprotective and pro-resolving transcriptional programs^81–85^. Similarly, CREB can promote either sensitization or recovery depending on context, supporting both pain and anti-inflammatory gene expression^42,86–91^. Blocking CB1R likely disrupts these dual transcriptional programs by removing ERK and AKT signaling support^92,93^, leading to a collapse of NF-κB and CREB phosphorylation^94^ and thereby promoting loss of TNFR2-driven analgesia. In contrast, females exhibited the opposite pattern. TNFR2 agonism decreased hippocampal p-NF-κB and p-CREB levels relative to injury controls, suggesting suppression of inflammatory and stress-linked signaling. Notably, AM251 co-treatment did not reverse these decreases, implying that TNFR2 signaling in females occurs through CB1R-independent mechanisms. Together, these findings reveal that CB1R signaling is required for TNFR2-mediated pain relief in both sexes, but the molecular direction of this regulation diverges: TNFR2 enhances NF-κB/CREB activation in males but suppresses it in females.

## Conclusions

In summary, this study identifies a novel dual-analgesic mechanism orchestrated by neuronal TNFR2 signaling within hippocampal CA3 pyramidal neurons. Activation of TNFR2 in these cells engages two complementary intrinsic pathways: an opioid axis driven by the upregulation of Pomc and its peptide product β-endorphin, and an endocannabinoid axis centered on downstream CB1R mediated signaling cascades. Together, these pathways promote the resolution of chronic neuropathic pain through activation of endogenous β-endorphin release and separately, cannabinoid receptor activity. By uncovering this intrinsic TNFR2-dependent opioid-endocannabinoid axis, our work provides direct evidence that TNFR2 agonism recruits analgesic systems to resolve neuropathic pain. These findings establish a mechanistic basis for TNFR2 as a therapeutic target capable of engaging neuroimmune mediated pain recovery programs. TNFR2 agonism has already been proposed as a therapeutic approach for neuropathic pain and other inflammatory disorders^95^. Our findings further support this by showing that the analgesic effects of TNFR2 agonisms occurs through activation of endogenous pain-relief pathways within the CNS. This work not only elucidates a novel neuroimmune mechanism through which hippocampus Nex neuron specific TNFR2 functions to resolves chronic pain but also provides a promising therapeutic strategy for restoring homeostatic balance in the context of chronic pain disorders.

## 2. Materials and Methods

### 2.1 Mice

Male and female mice were used in all experiments. Knockout (KO) animals (NexCre^ERT2^:TNFR2^F/F^) were generated in-house by crossing Neurod6^tm2.1(cre/ERT2)^ ^Kan^ animals (MGI:5308766; provided by Dr. K.-A. Nave) with TNFR2^loxP/loxP^ (TNFR2^F/F^) (EMMA ID: 05925; provided by Institut Clinique de la Souris). Age-matched C57BL/6J wild-type (WT) mice (JAX #000664) were purchased at ∼10 weeks of age from The Jackson Laboratory (Bar Harbor, ME) as indicated for specific experiments. Animals were housed in a specific-pathogen-free vivarium on a 12:12 h light/dark cycle with ad libitum access to food and water and were acclimated for ≥1 week before experiments. All procedures were approved by The George Washington University IACUC (protocols #A2023-066 and #A2023-061) and conducted in accordance with U.S. Public Health Service Policy.

### 2.2 Tamoxifen injections

Cre recombination was induced with tamoxifen (Sigma-Aldrich T5684; 75 mg/kg, i.p.) administered once daily for 5 consecutive days to NexCre^ERT2^:TNFR2^F/F^ mice and their TNFR2^F/F^ littermate controls at 10 weeks of age. Downstream procedures began 21 days after the final injection to allow sufficient time for recombination and tamoxifen clearance. The regimen was identical across sexes and genotypes.

### 2.3 Chronic constriction injury (CCI)

CCI was performed as described previously^96^. Briefly, mice were anesthetized with ketamine/xylazine (100/10 mg/kg, i.p.), and through a small mid-thigh incision, the right sciatic nerve was exposed, and three loose 6-0 silk ligatures (Oasis MV-711-V; Oasis Medical, Mettawa, IL) were placed ∼1.0-1.5 mm apart. Skin was closed with 5-0 nylon (Oasis MV-661). Animals recovered on a warming pad, returned to home cages upon regaining sternal posture, and received daily postoperative monitoring.

### 2.4 Mechanical Allodynia

Mechanical sensitivity (von Frey) was assessed 3 days before CCI (baseline) and weekly thereafter. Mice were placed individually on an elevated wire mesh under a transparent enclosure and habituated for 45-60 min. Calibrated monofilaments (Touch-Test Sensory Evaluators; 0.02-2.0 g) were applied perpendicularly to the plantar surface (ipsilateral, then contralateral) until the filament just bent, held ∼4-5 s, and removed. Testing used a staircase paradigm beginning at 0.4 g. Within each trial, a filament was applied five times at ∼60 s intervals; a response prompted testing with the next lower force, and no response prompted the next higher force. If withdrawal occurred at 0.02 g, the next trial used the same 0.02 g cutoff; if no response at 2.0 g, the next trial used the 2.0 g cutoff. Each session comprised six trials per paw. Responses were scored only when withdrawal was accompanied by directed attention/looking, guarding, licking, or clear avoidance; movements related to locomotion or weight-shifting were not counted. For each paw and time point, the withdrawal threshold (g) was the force eliciting a response in each trial, averaged across six trials.

### 2.5 Drugs and dosing

A TNFR2 agonist (10 mg/kg, i.p.), composed of a covalently stabilized TNFR2-selective single-chain TNF trimer fused to an effector-deficient Fc domain diluted in sterile PBS (Thermo Fisher J67670.K2). Doses were administered on post-injury days (dpi) 7, 10, and 13. The CB1 receptor antagonist AM251 (MedChemExpress HY-15443; 10 mg/kg, i.p.) was prepared in DMSO (Sigma-Aldrich 34869) and corn oil (Thermo Fisher 8001-30-7) and given on dpi 14, 21, and 28. The μ-opioid receptor antagonist naltrexone (Sigma-Aldrich N3136; 10 mg/kg, s.c.) was diluted in DMSO and PBS and administered every day from dpi 7–21. Injections were weight-normalized and performed at a consistent time of day. Vehicle controls (PBS; DMSO+corn oil; or DMSO+PBS) were administered on matched schedules. Animals were randomized to treatment.

### 2.6 RNA isolation and Bulk RNA sequencing

Six weeks after CCI, mice were perfused with PBS and the hippocampus dissected. Total RNA was extracted with TRIzol (Thermo Fisher Scientific, Waltham, MA) according to the manufacturer’s instructions. RNA quantity and purity were assessed by NanoDrop, and integrity was confirmed using an Agilent Bioanalyzer 2100. Samples with RIN ≥ 7.0 were used for library preparation.

Hippocampal tissue was collected from male mice (n = 4-5 per group) across six conditions: naïve TNFR2^F/F^ and NexCre^ERT2^:TNFR2^F/F^; vehicle-treated CCI TNFR2^F/F^ and CCI NexCre^ERT2^:TNFR2^F/F^; TNFR2 agonist-treated CCI TNFR2^F/F^ and CCI NexCre^ERT2^:TNFR2^F/F^. RNA libraries were prepared using the Illumina TruSeq Stranded mRNA kit and sequenced at MedGenome on an Illumina NovaSeq platform (100 bp paired-end, ∼45-60 million reads/sample). Demultiplexed FASTQ files were used for downstream analysis.

### 2.7 RNA Sequencing Data Analysis

#### Read Processing and Alignment

Raw reads were quality-checked with FastQC, and adapters and low-quality bases were removed using Cutadapt. High-quality reads were aligned to the Mus musculus reference genome (GRCm38.p6, Ensembl release 92) using the STAR two-pass alignment pipeline. Gene counts were quantified using HTSeq, and differential expression analysis was performed using DESeq2. Genes with |log2 fold change| ≥ 1 and adjusted p < 0.05 were considered significantly differentially expressed.

#### Volcano Plot Analysis

Volcano plots were generated in R (ggplot2 and ggrepel) to visualize differential expression across key comparisons. Genes previously implicated in opioid signaling, endocannabinoid (ECS) pathways, or both were selectively color-coded and labeled to highlight pathway-relevant expression changes. Significance thresholds were indicated at log2FC ±1 and p = 0.05. All volcano plots used the same standardized plotting function.

#### Reciprocal Differential Expression Analysis

To identify transcriptional changes specifically dependent on TNFR2 signaling in Nex-expressing excitatory neurons, we performed a reciprocal DEG analysis. First, DEGs induced by TNFR2 agonist treatment in TNFR2^F/F^ mice (vehicle-treated CCI TNFR2^F/F^ vs TNFR2 agonist-treated CCI TNFR2^F/F^) were identified. These were compared with DEGs altered by agonist treatment in NexCre^ERT2^:TNFR2^F/F^ mice (TNFR2 agonist-treated CCI TNFR2^F/F^ vs TNFR2 agonist-treated CCI NexCre^ERT2^:TNFR2^F/F^). Genes showing opposite-direction regulation between these two comparisons (adjusted p < 0.05) were classified as reciprocally expressed. Log2 fold-change values for these genes were analyzed.

#### GO Enrichment and CNETplot Analysis

Gene Ontology (GO) enrichment analysis was performed using clusterProfiler (enrichGO) with Benjamini–Hochberg correction (p < 0.05, q < 0.20). Enriched GO Biological Process terms were visualized using cnetplots (enrichplot), allowing visualization of gene–pathway relationships within reciprocally regulated gene sets. Analyses were run using mouse gene annotations (org.Mm.eg.db), and only terms meeting enrichment criteria were included.

### 2.8 Immunohistochemistry and RNAscope

Mice were perfused transcardially with 4% paraformaldehyde (PFA; Sigma-Aldrich) in PBS. Brains were post-fixed overnight in 4% PFA, cryoprotected in 25% sucrose for 48 h at 4 °C, embedded in OCT (Tissue-Tek), and stored at −80 °C. Cryosections (15 µm) were cut (Leica) and permeabilized in PBS/0.3% Triton X-100 (10 min, room temperature). Nonspecific binding was blocked with PBS/4% BSA (30 min). Primary antibodies were applied overnight at 4 °C in blocking buffer: POMC (goat, 1:200; Novus) and β-endorphin (rat, 1:200; Phoenix Pharmaceuticals). After three 10-minute PBS washes, species-specific secondary antibodies (Alexa Fluor 594, 1:500; Invitrogen) were applied in 1% normal goat serum/PBS for 45 min at room temperature. Slides were washed (3 × 10 min), counterstained with Hoechst (1:10,000 in PBS, 10 min), washed, mounted with Fluoromount (Electron Microscopy Sciences; Fluoro-Gel with TES), and covered by a coverslip. Images were acquired on a Zeiss spinning-disk microscope at the GW Nanofabrication and Imaging Center at 25X magnification (NIH S10OD010710).

Fixed-frozen sections were processed according to the manufacturer’s protocol (ACDbio). Briefly, slides were incubated at −20 °C for 120 min, rinsed in PBS, dry-baked for 60 min, and post-fixed in 4% PFA for 15 min. Sections were dehydrated through 50%, 70%, and 100% ethanol; treated with hydrogen peroxide; and subjected to target retrieval (heated Target Retrieval Buffer, Cat. #322000) and Protease III (Cat. #322381). Probes were applied for Tnfrsf1a-C1 (Cat. #438941), Tnfrsf1b-C2 (Cat. #438951-C2), and Neurod6-C3 (Cat. #444851). Amplification used sequential AMP1/AMP2/AMP3 (Cat. #323110). Channel-specific HRP was applied for C1, followed by Opal 690 (1:1000). HRP was quenched, and the process was repeated for C2 (Opal 570, 1:500) and C3 (Opal 520, 1:1000). Slides were counterstained, covered by a coverslip, and stored at 4 °C until imaging.

For each animal, four coronal hippocampal sections containing the CA3 region were analyzed, spaced 90 µm apart. Within these sections, POMC, β-endorphin, and TNFR2 were manually counted. For immunohistochemistry, POMC and β-endorphin puncta adjacent to DAPI-labeled nuclei were considered positive. For RNAscope, TNFR2 puncta were counted only when colocalized with DAPI and Nex-labeled neurons. All quantification was performed by an observer blinded to group and sex.

### 2.9 Western Blot

Proteins were separated by SDS-polyacrylamide gel electrophoresis (SDS-PAGE) and transferred to nitrocellulose using a Trans-Blot Turbo system (Bio-Rad). Membranes were blocked for 1 h at room temperature in 5% BSA (Cell Signaling) prepared in TBS-T (10 mM Tris-HCl, pH 7.5; 150 mM NaCl; 0.1% Tween-20). Primary antibodies diluted in the same blocking buffer were applied overnight at 4 °C with gentle rocking: anti-p-CREB (mouse, 1:750, Cell Signaling #9196), anti-CREB (rabbit, 1:1000, Cell Signaling #9197), anti-NF-κB p65 (rabbit, 1:1000, Abcam ab16502), and anti-phospho-NF-κB p65 (rabbit, 1:750, Abcam ab76302). After washing, membranes were incubated with species-specific HRP-conjugated secondary antibodies and developed by chemiluminescence (West Pico; Thermo Fisher Scientific). Band intensities were quantified in FIJI (ImageJ; NIH). For loading control, total protein per lane was assessed by Ponceau S staining (Thermo Fisher) and then normalized to the control conditions.

### 2.10 Statistics

Data were analyzed in GraphPad Prism 10.0 (GraphPad Software, San Diego, CA, USA) and are presented as mean ± SEM. Normality was assessed before analysis. Two-group comparisons used t-tests; multiple groups used one-way or two-way ANOVA with appropriate multiple-comparison procedures. P < 0.05 was considered significant.

### Abbreviations

CNP: Chronic neuropathic pain
CNS: Central nervous system
CCI: Chronic constriction injury
TNF: Tumor necrosis factor
tmTNF: Transmembrane TNF
TNFR2: Tumor necrosis factor receptor 2
Hc: Hippocampus
POMC: Proopiomelanocortin
β-End: β-endorphins
MOR: Mu-opioid receptor
CA1 and CA3: Cornu ammonis 1 and 3
ECS: Endocannabinoid system
CB_1_R and CB_2_R: Cannabinoid receptors
OLAH: Oleoyl ACP hydrolase
Dpi: Days post injury
NTX: Naltrexone

## Author Contribution

S.A., P.F., R.F., V.J.T., and J.R.B. designed research; S.A., E.J., K.L.N., and V.B.R. performed research; J.R.B. contributed new reagents/analytic tools; S.A., and S.G. analyzed data, and S.A., P.F., and A.M. wrote the paper.

## Acknowledgments

This work was supported by the Department of Defense (DOD) #CP200074 (J.R.B)

## References

(1) Costigan, M.; Scholz, J.; Woolf, C. J. Neuropathic Pain. Annu Rev Neurosci 2009, 32, 1–32. 10.1146/annurev.neuro.051508.135531.

(2) Rikard, S. M. Chronic Pain Among Adults — United States, 2019–2021. MMWR Morb Mortal Wkly Rep 2023, 72. 10.15585/mmwr.mm7215a1.

(3) Jensen, T. S.; Finnerup, N. B. Allodynia and Hyperalgesia in Neuropathic Pain: Clinical Manifestations and Mechanisms. The Lancet Neurology 2014, 13 (9), 924–935. 10.1016/S1474-4422(14)70102-4.

(4) CDC. SUDORS Dashboard: Fatal Drug Overdose Data. Overdose Prevention. https://www.cdc.gov/overdose-prevention/data-research/facts-stats/sudors-dashboard-fatal-overdose-data.html (accessed 2025-11-20).

(5) Harden, N.; Cohen, M. Unmet Needs in the Management of Neuropathic Pain. Journal of Pain and Symptom Management 2003, 25 (5), S12–S17. 10.1016/S0885-3924(03)00065-4.

(6) Garnett, M. F.; Miniño, A. M. Drug Overdose Deaths in the United States, 2003-2023. 10.15620/cdc/170565.

(7) Lindenlaub, T.; Sommer, C. Cytokines in Sural Nerve Biopsies from Inflammatory and Non-Inflammatory Neuropathies. Acta Neuropathol 2003, 105 (6), 593–602. 10.1007/s00401-003-0689-y.

(8) Fischer, R.; Kontermann, R. E.; Maier, O. Targeting sTNF/TNFR1 Signaling as a New Therapeutic Strategy. Antibodies 2015, 4 (1), 48–70. 10.3390/antib4010048.

(9) Probert, L. TNF and Its Receptors in the CNS: The Essential, the Desirable and the Deleterious Effects. Neuroscience 2015, 302, 2–22. 10.1016/j.neuroscience.2015.06.038.

(10) Fischer, R.; Sendetski, M.; del Rivero, T.; Martinez, G. F.; Bracchi-Ricard, V.; Swanson, K. A.; Pruzinsky, E. K.; Delguercio, N.; Rosalino, M. J.; Padutsch, T.; Kontermann, R. E.; Pfizenmaier, K.; Bethea, J. R. TNFR2 Promotes Treg-Mediated Recovery from Neuropathic Pain across Sexes. Proceedings of the National Academy of Sciences 2019, 116 (34), 17045–17050. 10.1073/pnas.1902091116.

(11) Baliki, M. N.; Apkarian, A. V. Nociception, Pain, Negative Moods and Behavior Selection. Neuron 2015, 87 (3), 474–491. 10.1016/j.neuron.2015.06.005.

(12) Tajerian, M.; Hung, V.; Nguyen, H.; Lee, G.; Joubert, L.-M.; Malkovskiy, A. V.; Zou, B.; Xie, S.; Huang, T.-T.; Clark, J. D. The Hippocampal Extracellular Matrix Regulates Pain and Memory after Injury. Mol Psychiatry 2018, 23 (12), 2302–2313. 10.1038/s41380-018-0209-z.

(13) Kalman, E.; Keay, K. A. Different Patterns of Morphological Changes in the Hippocampus and Dentate Gyrus Accompany the Differential Expression of Disability Following Nerve Injury. Journal of Anatomy 2014, 225 (6), 591–603. 10.1111/joa.12238.

(14) Mutso, A. A.; Radzicki, D.; Baliki, M. N.; Huang, L.; Banisadr, G.; Centeno, M. V.; Radulovic, J.; Martina, M.; Miller, R. J.; Apkarian, A. V. Abnormalities in Hippocampal Functioning with Persistent Pain. J Neurosci 2012, 32 (17), 5747–5756. 10.1523/JNEUROSCI.0587-12.2012.

(15) Mutso, A. A.; Petre, B.; Huang, L.; Baliki, M. N.; Torbey, S.; Herrmann, K. M.; Schnitzer, T. J.; Apkarian, A. V. Reorganization of Hippocampal Functional Connectivity with Transition to Chronic Back Pain. J Neurophysiol 2014, 111 (5), 1065–1076. 10.1152/jn.00611.2013.

(16) Zheng, J.; Jiang, Y.-Y.; Xu, L.-C.; Ma, L.-Y.; Liu, F.-Y.; Cui, S.; Cai, J.; Liao, F.-F.; Wan, Y.; Yi, M. Adult Hippocampal Neurogenesis along the Dorsoventral Axis Contributes Differentially to Environmental Enrichment Combined with Voluntary Exercise in Alleviating Chronic Inflammatory Pain in Mice. J Neurosci 2017, 37 (15), 4145–4157. 10.1523/JNEUROSCI.3333-16.2017.

(17) Wang, Y.; Liu, N.; Ma, L.; Yue, L.; Cui, S.; Liu, F.-Y.; Yi, M.; Wan, Y. Ventral Hippocampal CA1 Pyramidal Neurons Encode Nociceptive Information. Neurosci. Bull. 2024, 40 (2), 201–217. 10.1007/s12264-023-01086-x.

(18) Swanson, K. A.; Nguyen, K. L.; Gupta, S.; Ricard, J.; Bethea, J. R. TNFR1/p38αMAPK Signaling in Nex+ Supraspinal Neurons Regulates Estrogen-Dependent Chronic Neuropathic Pain. Brain Behav Immun 2024, 119, 261–271. 10.1016/j.bbi.2024.03.050.

(19) Agarwal, A.; Dibaj, P.; Kassmann, C. M.; Goebbels, S.; Nave, K.-A.; Schwab, M. H. In Vivo Imaging and Noninvasive Ablation of Pyramidal Neurons in Adult NEX-CreERT2 Mice. Cereb Cortex 2012, 22 (7), 1473–1486. 10.1093/cercor/bhr214.

(20) Palacios-Filardo, J.; Mellor, J. R. Neuromodulation of Hippocampal Long-Term Synaptic Plasticity. Curr Opin Neurobiol 2019, 54, 37–43. 10.1016/j.conb.2018.08.009.

(21) Carney, B. N.; Illiano, P.; Pohl, T. M.; Desu, H. L.; Mini, A.; Mudalegundi, S.; Asencor, A. I.; Jwala, S.; Ascona, M. C.; Singh, P. K.; Titus, D. J.; Pazarlar, B. A.; Wang, L.; Bianchi, L.; Mikkelsen, J. D.; Atkins, C. M.; Lambertsen, K. L.; Brambilla, R. Astroglial TNFR2 Signaling Regulates Hippocampal Synaptic Function and Plasticity in a Sex Dependent Manner. Brain, Behavior, and Immunity 2025, 129, 757–777. 10.1016/j.bbi.2025.07.006.

(22) Harno, E.; Gali Ramamoorthy, T.; Coll, A. P.; White, A. POMC: The Physiological Power of Hormone Processing. Physiol Rev 2018, 98 (4), 2381–2430. 10.1152/physrev.00024.2017.

(23) Pandey, V.; Yadav, V.; Singh, R.; Srivastava, A.; Subhashini. β-Endorphin (an Endogenous Opioid) Inhibits Inflammation, Oxidative Stress and Apoptosis via Nrf-2 in Asthmatic Murine Model. Sci Rep 2023, 13 (1), 12414. 10.1038/s41598-023-38366-5.

(24) YEARS OF POMC: Lipotropin and beta-endorphin: a perspective in: Journal of Molecular Endocrinology Volume 56 Issue 4 (2016). https://jme.bioscientifica.com/view/journals/jme/56/4/T13.xml (accessed 2025-08-08).

(25) Shen, Y.; Tian, M.; Zheng, Y.; Gong, F.; Fu, A. K. Y.; Ip, N. Y. Stimulation of the Hippocampal POMC/MC4R Circuit Alleviates Synaptic Plasticity Impairment in an Alzheimer’s Disease Model. Cell Reports 2016, 17 (7), 1819–1831. 10.1016/j.celrep.2016.10.043.

(26) Woodhams, S. G.; Sagar, D. R.; Burston, J. J.; Chapman, V. The Role of the Endocannabinoid System in Pain. In Pain Control; Schaible, H.-G., Ed.; Springer: Berlin, Heidelberg, 2015; pp 119–143. 10.1007/978-3-662-46450-2_7.

(27) Jia, X.; Crawford, J. C.; Gebregzabher, D.; Monson, E. A.; Mettelman, R. C.; Wan, Y.; Ren, Y.; Chou, J.; Novak, T.; McQuilten, H. A.; Clarke, M.; Bachem, A.; Foo, I. J.; Fritzlar, S.; Montoya, J. C.; Trenerry, A. M.; Nie, S.; Leeming, M. G.; Nguyen, T. H. O.; Kedzierski, L.; Littler, D. R.; Kueh, A.; Cardamone, T.; Wong, C. Y.; Hensen, L.; Cabug, A.; Laguna, J. G.; Agrawal, M.; Flerlage, T.; Boyd, D. F.; Velde, L.-A. V. de; Habel, J. R.; Loh, L.; Koay, H.-F.; Sandt, C. E. van de; Konstantinov, I. E.; Berzins, S. P.; Flanagan, K. L.; Wakim, L. M.; Herold, M. J.; Green, A. M.; Smallwood, H. S.; Rossjohn, J.; Thwaites, R. S.; Chiu, C.; Scott, N. E.; Mackenzie, J. M.; Bedoui, S.; Reading, P. C.; Londrigan, S. L.; Helbig, K. J.; Randolph, A. G.; Thomas, P. G.; Xu, J.; Wang, Z.; Chua, B. Y.; Kedzierska, K. High Expression of Oleoyl-ACP Hydrolase Underpins Life-Threatening Respiratory Viral Diseases. Cell 2024, 187 (17), 4586–4604.e20. 10.1016/j.cell.2024.07.026.

(28) Merkler, D. J.; Chew, G. H.; Gee, A. J.; Merkler, K. A.; Sorondo, J.-P. O.; Johnson, M. E. Oleic Acid Derived Metabolites in Mouse Neuroblastoma N18TG2 Cells. Biochemistry 2004, 43 (39), 12667–12674. 10.1021/bi049529p.

(29) Hiley, C. R.; Hoi, P. M. Oleamide: A Fatty Acid Amide Signaling Molecule in the Cardiovascular System? Cardiovascular Drug Reviews 2007, 25, 46–60. 10.1111/j.1527-3466.2007.00004.x.

(30) Kędziora, M.; Boccella, S.; Marabese, I.; Mlost, J.; Infantino, R.; Maione, S.; Starowicz, K. Inhibition of Anandamide Breakdown Reduces Pain and Restores LTP and Monoamine Levels in the Rat Hippocampus via the CB1 Receptor Following Osteoarthritis. Neuropharmacology 2023, 222, 109304. 10.1016/j.neuropharm.2022.109304.

(31) Dong, Y.; Fischer, R.; Naudé, P. J. W.; Maier, O.; Nyakas, C.; Duffey, M.; Van der Zee, E. A.; Dekens, D.; Douwenga, W.; Herrmann, A.; Guenzi, E.; Kontermann, R. E.; Pfizenmaier, K.; Eisel, U. L. M. Essential Protective Role of Tumor Necrosis Factor Receptor 2 in Neurodegeneration. Proc Natl Acad Sci U S A 2016, 113 (43), 12304–12309. 10.1073/pnas.1605195113.

(32) Ma, L.; Sun, P.; Zhang, J.-C.; Zhang, Q.; Yao, S.-L. Proinflammatory Effects of S100A8/A9 via TLR4 and RAGE Signaling Pathways in BV-2 Microglial Cells. Int J Mol Med 2017, 40 (1), 31–38. 10.3892/ijmm.2017.2987.

(33) Maganin, A. G.; Souza, G. R.; Fonseca, M. D.; Lopes, A. H.; Guimarães, R. M.; Dagostin, A.; Cecilio, N. T.; Mendes, A. S.; Gonçalves, W. A.; Silva, C. E. A.; Fernandes Gomes, F. I.; Mauriz Marques, L. M.; Silva, R. L.; Arruda, L. M.; Santana, D. A.; Lemos, H.; Huang, L.; Davoli-Ferreira, M.; Santana-Coelho, D.; Sant’Anna, M. B.; Kusuda, R.; Talbot, J.; Pacholczyk, G.; Buqui, G. A.; Lopes, N. P.; Alves-Filho, J. C.; Leão, R. M.; O’Connor, J. C.; Cunha, F. Q.; Mellor, A.; Cunha, T. M. Meningeal Dendritic Cells Drive Neuropathic Pain through Elevation of the Kynurenine Metabolic Pathway in Mice. J Clin Invest 132 (23), e153805. 10.1172/JCI153805.

(34) Wu, J.; Wu, H.; An, J.; Ballantyne, C. M.; Cyster, J. G. Critical Role of Integrin CD11c in Splenic Dendritic Cell Capture of Missing-Self CD47 Cells to Induce Adaptive Immunity. Proc Natl Acad Sci U S A 2018, 115 (26), 6786–6791. 10.1073/pnas.1805542115.

(35) Lau, J. K. Y.; Tian, M.; Shen, Y.; Lau, S.-F.; Fu, W.-Y.; Fu, A. K. Y.; Ip, N. Y. Melanocortin Receptor Activation Alleviates Amyloid Pathology and Glial Reactivity in an Alzheimer’s Disease Transgenic Mouse Model. Sci Rep 2021, 11, 4359. 10.1038/s41598-021-83932-4.

(36) Tanaka, S. Comparative Aspects of Intracellular Proteolytic Processing of Peptide Hormone Precursors: Studies of Proopiomelanocortin Processing. jzoo 2003, 20 (10), 1183–1198. 10.2108/zsj.20.1183.

(37) Facchinetti, F.; Petraglia, F.; Genazzani, A. R. Localization and Expression of the Three Opioid Systems. Seminars in Reproductive Endocrinology 2008, 5, 103–113. 10.1055/s-2007-1021858.

(38) Raynor, K.; Kong, H.; Chen, Y.; Yasuda, K.; Yu, L.; Bell, G. I.; Reisine, T. Pharmacological Characterization of the Cloned Kappa-, Delta-, and Mu-Opioid Receptors. Mol Pharmacol 1994, 45 (2), 330–334.

(39) Gonzalez, G. A.; Montminy, M. R. Cyclic AMP Stimulates Somatostatin Gene Transcription by Phosphorylation of CREB at Serine 133. Cell 1989, 59 (4), 675–680. 10.1016/0092-8674(89)90013-5.

(40) Sheng, M.; Thompson, M. A.; Greenberg, M. E. CREB: A Ca2+-Regulated Transcription Factor Phosphorylated by Calmodulin-Dependent Kinases. Science 1991, 252 (5011), 1427–1430. 10.1126/science.1646483.

(41) Ma, W.; Quirion, R. Increased Phosphorylation of Cyclic AMP Response Element-Binding Protein (CREB) in the Superficial Dorsal Horn Neurons Following Partial Sciatic Nerve Ligation. Pain 2001, 93 (3), 295–301. 10.1016/S0304-3959(01)00335-9.

(42) Wen, A. Y.; Sakamoto, K. M.; Miller, L. S. The Role of the Transcription Factor CREB in Immune Function. J Immunol 2010, 185 (11), 6413–6419. 10.4049/jimmunol.1001829.

(43) Avni, D.; Ernst, O.; Philosoph, A.; Zor, T. Role of CREB in Modulation of TNFα and IL-10 Expression in LPS-Stimulated RAW264.7 Macrophages. Molecular Immunology 2010, 47 (7), 1396–1403. 10.1016/j.molimm.2010.02.015.

(44) del Rivero, T.; Fischer, R.; Yang, F.; Swanson, K. A.; Bethea, J. R. Tumor Necrosis Factor Receptor 1 Inhibition Is Therapeutic for Neuropathic Pain in Males but Not in Females. PAIN 2019, 160 (4), 922. 10.1097/j.pain.0000000000001470.

(45) Fischer, R.; Padutsch, T.; Bracchi-Ricard, V.; Murphy, K. L.; Martinez, George. F.; Delguercio, N.; Elmer, N.; Sendetski, M.; Diem, R.; Eisel, U. L. M.; Smeyne, R. J.; Kontermann, R. E.; Pfizenmaier, K.; Bethea, J. R. Exogenous Activation of Tumor Necrosis Factor Receptor 2 Promotes Recovery from Sensory and Motor Disease in a Model of Multiple Sclerosis. Brain Behav Immun 2019, 81, 247–259. 10.1016/j.bbi.2019.06.021.

(46) Biglari, N.; Gaziano, I.; Schumacher, J.; Radermacher, J.; Paeger, L.; Klemm, P.; Chen, W.; Corneliussen, S.; Wunderlich, C. M.; Sue, M.; Vollmar, S.; Klöckener, T.; Sotelo-Hitschfeld, T.; Abbasloo, A.; Edenhofer, F.; Reimann, F.; Gribble, F. M.; Fenselau, H.; Kloppenburg, P.; Wunderlich, F. T.; Brüning, J. C. Functionally Distinct POMC-Expressing Neuron Subpopulations in Hypothalamus Revealed by Intersectional Targeting. Nat Neurosci 2021, 24 (7), 913–929. 10.1038/s41593-021-00854-0.

(47) Schwartz, G. J.; Fu, J.; Astarita, G.; Li, X.; Gaetani, S.; Campolongo, P.; Cuomo, V.; Piomelli, D. The Lipid Messenger OEA Links Dietary Fat Intake to Satiety. Cell Metab 2008, 8 (4), 281–288. 10.1016/j.cmet.2008.08.005.

(48) Guijarro, A.; Fu, J.; Astarita, G.; Piomelli, D. CD36 Gene Deletion Decreases Oleoylethanolamide Levels in Small Intestine of Free-Feeding Mice. Pharmacological Research 2010, 61 (1), 27–33. 10.1016/j.phrs.2009.09.003.

(49) Yang, L.; Guo, H.; Li, Y.; Meng, X.; Yan, L.; Dan Zhang; Wu, S.; Zhou, H.; Peng, L.; Xie, Q.; Jin, X. Oleoylethanolamide Exerts Anti-Inflammatory Effects on LPS-Induced THP-1 Cells by Enhancing PPARα Signaling and Inhibiting the NF-κB and ERK1/2/AP-1/STAT3 Pathways. Sci Rep 2016, 6, 34611. 10.1038/srep34611.

(50) van Zadelhoff, G.; van der Stelt, M. Oxygenation of Anandamide by Lipoxygenases. In Endocannabinoid Signaling: Methods and Protocols; Maccarrone, M., Ed.; Springer: New York, NY, 2016; pp 217–225. 10.1007/978-1-4939-3539-0_22.

(51) Ivanov, I.; Kakularam, K. R.; Shmendel, E. V.; Rothe, M.; Aparoy, P.; Heydeck, D.; Kuhn, H. Oxygenation of Endocannabinoids by Mammalian Lipoxygenase Isoforms. Biochimica et Biophysica Acta (BBA) - Molecular and Cell Biology of Lipids 2021, 1866 (6), 158918. 10.1016/j.bbalip.2021.158918.

(52) Kozak, K. R.; Gupta, R. A.; Moody, J. S.; Ji, C.; Boeglin, W. E.; DuBois, R. N.; Brash, A. R.; Marnett, L. J. 15-Lipoxygenase Metabolism of 2-Arachidonylglycerol: GENERATION OF A PEROXISOME PROLIFERATOR-ACTIVATED RECEPTOR α AGONIST *. Journal of Biological Chemistry 2002, 277 (26), 23278–23286. 10.1074/jbc.M201084200.

(53) Zhou, X.; Spittau, B.; Krieglstein, K. TGFβ Signalling Plays an Important Role in IL4-Induced Alternative Activation of Microglia. Journal of Neuroinflammation 2012, 9 (1), 210. 10.1186/1742-2094-9-210.

(54) Ferrisi, R.; Gado, F.; Ricardi, C.; Polini, B.; Manera, C.; Chiellini, G. The Interplay between Cannabinoid Receptors and Microglia in the Pathophysiology of Alzheimer’s Disease. Journal of Clinical Medicine 2023, 12 (23), 7201. 10.3390/jcm12237201.

(55) Mecha, M.; Feliú, A.; Carrillo-Salinas, F. J.; Rueda-Zubiaurre, A.; Ortega-Gutiérrez, S.; de Sola, R. G.; Guaza, C. Endocannabinoids Drive the Acquisition of an Alternative Phenotype in Microglia. Brain, Behavior, and Immunity 2015, 49, 233–245. 10.1016/j.bbi.2015.06.002.

(56) Margheritis, E.; Castellani, B.; Magotti, P.; Peruzzi, S.; Romeo, E.; Natali, F.; Mostarda, S.; Gioiello, A.; Piomelli, D.; Garau, G. Bile Acid Recognition by NAPE-PLD. ACS Chem Biol 2016, 11 (10), 2908–2914. 10.1021/acschembio.6b00624.

(57) Trojan, V.; Landa, L.; Šulcová, A.; Slíva, J.; Hřib, R. The Main Therapeutic Applications of Cannabidiol (CBD) and Its Potential Effects on Aging with Respect to Alzheimer’s Disease. Biomolecules 2023, 13 (10), 1446. 10.3390/biom13101446.

(58) Trivedi, M. K.; Mondal, S.; Gangwar, M.; Jana, S. Effects of Cannabidiol Interactions with CYP2R1, CYP27B1, CYP24A1, and Vitamin D3 Receptors on Spatial Memory, Pain, Inflammation, and Aging in Vitamin D3 Deficiency Diet-Induced Rats. Cannabis and Cannabinoid Research 2023, 8 (6), 1019–1029. 10.1089/can.2021.0240.

(59) Maher, D. P.; Walia, D.; Heller, N. M. Suppression of Human Natural Killer Cells by Different Classes of Opioids. Anesth Analg 2019, 128 (5), 1013–1021. 10.1213/ANE.0000000000004058.

(60) Liang, X.; Liu, R.; Chen, C.; Ji, F.; Li, T. Opioid System Modulates the Immune Function: A Review. Transl Perioper Pain Med 2016, 1 (1), 5–13.

(61) Hutchinson, M. R.; Northcutt, A. L.; Hiranita, T.; Wang, X.; Lewis, S. S.; Thomas, J.; van Steeg, K.; Kopajtic, T. A.; Loram, L. C.; Sfregola, C.; Galer, E.; Miles, N. E.; Bland, S. T.; Amat, J.; Rozeske, R. R.; Maslanik, T.; Chapman, T. R.; Strand, K. A.; Fleshner, M.; Bachtell, R. K.; Somogyi, A. A.; Yin, H.; Katz, J. L.; Rice, K. C.; Maier, S. F.; Watkins, L. R. Opioid Activation of Toll-Like Receptor 4 Contributes to Drug Reinforcement. J Neurosci 2012, 32 (33), 11187–11200. 10.1523/JNEUROSCI.0684-12.2012.

(62) Alemi, F.; Kwon, E.; Poole, D. P.; Lieu, T.; Lyo, V.; Cattaruzza, F.; Cevikbas, F.; Steinhoff, M.; Nassini, R.; Materazzi, S.; Guerrero-Alba, R.; Valdez-Morales, E.; Cottrell, G. S.; Schoonjans, K.; Geppetti, P.; Vanner, S. J.; Bunnett, N. W.; Corvera, C. U. The TGR5 Receptor Mediates Bile Acid–Induced Itch and Analgesia. J Clin Invest 2013, 123 (4), 1513–1530. 10.1172/JCI64551.

(63) Loh, H. H.; Tseng, L. F.; Wei, E.; Li, C. H. Beta-Endorphin Is a Potent Analgesic Agent. Proc Natl Acad Sci U S A 1976, 73 (8), 2895–2898. 10.1073/pnas.73.8.2895.

(64) De Riu, P. L.; Petruzzi, V.; Caria, M. A.; Mameli, O.; Casu, A. R.; Nuvoli, S.; Spanu, A.; Madeddu, G. β-Endorphin and Cortisol Levels in Plasma and CSF Following Acute Experimental Spinal Traumas. Physiology & Behavior 1997, 62 (1), 1–5. 10.1016/S0031-9384(97)00099-1.

(65) Mousa, S. A.; Zhang, Q.; Sitte, N.; Ji, R.-R.; Stein, C. β-Endorphin-Containing Memory-Cells and μ-Opioid Receptors Undergo Transport to Peripheral Inflamed Tissue. Journal of Neuroimmunology 2001, 115 (1), 71–78. 10.1016/S0165-5728(01)00271-5.

(66) Brunton, P. J.; Meddle, S. L.; Ma, S.; Ochedalski, T.; Douglas, A. J.; Russell, J. A. Endogenous Opioids and Attenuated Hypothalamic-Pituitary-Adrenal Axis Responses to Immune Challenge in Pregnant Rats. J Neurosci 2005, 25 (21), 5117–5126. 10.1523/JNEUROSCI.0866-05.2005.

(67) Sacerdote, P.; Di San Secondo, V. E. M. R.; Sirchia, G.; Manfredi, B.; Panerai, A. E. Endogenous Opioids Modulate Allograft Rejection Time in Mice: Possible Relation with Th1/Th2 Cytokines. Clin Exp Immunol 1998, 113 (3), 465–469. 10.1046/j.1365-2249.1998.00680.x.

(68) Spinal microglial β-endorphin signaling mediates IL-10 and exenatide-induced inhibition of synaptic plasticity in neuropathic pain - Ma - 2021 - CNS Neuroscience & Therapeutics - Wiley Online Library. https://onlinelibrary.wiley.com/doi/10.1111/cns.13694 (accessed 2025-11-02).

(69) Rheins, L. A.; Cotleur, A. L.; Kleier, R. S.; Hoppenjans, W. B.; Saunder, D. N.; Nordlund, J. J. Alpha-Melanocyte Stimulating Hormone Modulates Contact Hypersensitivity Responsiveness in C57/BL6 Mice. J Invest Dermatol 1989, 93 (4), 511–517. 10.1111/1523-1747.ep12284064.

(70) Labuz, D.; Schmidt, Y.; Schreiter, A.; Rittner, H. L.; Mousa, S. A.; Machelska, H. Immune Cell–Derived Opioids Protect against Neuropathic Pain in Mice. J Clin Invest 2009, 119 (2), 278–286. 10.1172/JCI36246.

(71) Laumet, G.; Ma, J.; Robison, A. J.; Kumari, S.; Heijnen, C. J.; Kavelaars, A. T Cells as an Emerging Target for Chronic Pain Therapy. Front. Mol. Neurosci. 2019, 12. 10.3389/fnmol.2019.00216.

(72) Sorge, R. E.; Mapplebeck, J. C. S.; Rosen, S.; Beggs, S.; Taves, S.; Alexander, J. K.; Martin, L. J.; Austin, J.-S.; Sotocinal, S. G.; Chen, D.; Yang, M.; Shi, X. Q.; Huang, H.; Pillon, N. J.; Bilan, P. J.; Tu, Y. S.; Klip, A.; Ji, R.-R.; Zhang, J.; Salter, M. W.; Mogil, J. S. Different Immune Cells Mediate Mechanical Pain Hypersensitivity in Male and Female Mice. Nat Neurosci 2015, 18 (8), 1081–1083. 10.1038/nn.4053.

(73) Descalzi, G.; Fukushima, H.; Suzuki, A.; Kida, S.; Zhuo, M. Genetic Enhancement of Neuropathic and Inflammatory Pain by Forebrain Upregulation of CREB-Mediated Transcription. Mol Pain 2012, 8, 1744–8069-8–90. 10.1186/1744-8069-8-90.

(74) Apryani, E.; Ali, U.; Wang, Z.-Y.; Wu, H.-Y.; Mao, X.-F.; Ahmad, K. A.; Li, X.-Y.; Wang, Y.-X. The Spinal Microglial IL-10/β-Endorphin Pathway Accounts for Cinobufagin-Induced Mechanical Antiallodynia in Bone Cancer Pain Following Activation of Α7-Nicotinic Acetylcholine Receptors. Journal of Neuroinflammation 2020, 17 (1), 75. 10.1186/s12974-019-1616-z.

(75) Duraffourd, C.; Kumala, E.; Anselmi, L.; Brecha, N. C.; Sternini, C. Opioid-Induced Mitogen-Activated Protein Kinase Signaling in Rat Enteric Neurons Following Chronic Morphine Treatment. PLOS ONE 2014, 9 (10), e110230. 10.1371/journal.pone.0110230.

(76) Xing, J.; Kornhauser, J. M.; Xia, Z.; Thiele, E. A.; Greenberg, M. E. Nerve Growth Factor Activates Extracellular Signal-Regulated Kinase and P38 Mitogen-Activated Protein Kinase Pathways To Stimulate CREB Serine 133 Phosphorylation. Mol Cell Biol 1998, 18 (4), 1946–1955. 10.1128/mcb.18.4.1946.

(77) Simon, J.; Arthur, C.; Fong, A. L.; Dwyer, J. M.; Davare, M.; Reese, E.; Obrietan, K.; Impey, S. Mitogen-and Stress-Activated Protein Kinase 1 Mediates cAMP Response Element-Binding Protein Phosphorylation and Activation by Neurotrophins. J Neurosci 2004, 24 (18), 4324–4332. 10.1523/JNEUROSCI.5227-03.2004.

(78) Hohmann, A. G.; Briley, E. M.; Herkenham, M. Pre- and Postsynaptic Distribution of Cannabinoid and Mu Opioid Receptors in Rat Spinal Cord. Brain Research 1999, 822 (1), 17–25. 10.1016/S0006-8993(98)01321-3.

(79) Hohmann, A. G.; Herkenham, M. Localization of Central Cannabinoid CB1 Receptor Messenger RNA in Neuronal Subpopulations of Rat Dorsal Root Ganglia: A Double-Label in Situ Hybridization Study. Neuroscience 1999, 90 (3), 923–931. 10.1016/S0306-4522(98)00524-7.

(80) Brusberg, M.; Arvidsson, S.; Kang, D.; Larsson, H.; Lindström, E.; Martinez, V. CB1 Receptors Mediate the Analgesic Effects of Cannabinoids on Colorectal Distension-Induced Visceral Pain in Rodents. J. Neurosci. 2009, 29 (5), 1554–1564. 10.1523/JNEUROSCI.5166-08.2009.

(81) Liu, D.; Zhong, Z.; Karin, M. NF-κB: A Double-Edged Sword Controlling Inflammation. Biomedicines 2022, 10 (6), 1250. 10.3390/biomedicines10061250.

(82) Dresselhaus, E. C.; Meffert, M. K. Cellular Specificity of NF-κB Function in the Nervous System. Front. Immunol. 2019, 10. 10.3389/fimmu.2019.01043.

(83) Gorbacheva, L.; Pinelis, V.; Ishiwata, S.; Strukova, S.; Reiser, G. Activated Protein C Prevents Glutamate- and Thrombin-Induced Activation of Nuclear Factor-κB in Cultured Hippocampal Neurons. Neuroscience 2010, 165 (4), 1138–1146. 10.1016/j.neuroscience.2009.11.027.

(84) Gorbacheva, L.; Strukova, S.; Pinelis, V.; Ishiwata, S.; Stricker, R.; Reiser, G. NF-κB-Dependent and - Independent Pathways in the Protective Effects of Activated Protein C in Hippocampal and Cortical Neurons at Excitotoxicity. Neurochemistry International 2013, 63 (2), 101–111. 10.1016/j.neuint.2013.05.008.

(85) Fridmacher, V.; Kaltschmidt, B.; Goudeau, B.; Ndiaye, D.; Rossi, F. M.; Pfeiffer, J.; Kaltschmidt, C.; Israël, A.; Mémet, S. Forebrain-Specific Neuronal Inhibition of Nuclear Factor-κB Activity Leads to Loss of Neuroprotection. J Neurosci 2003, 23 (28), 9403–9408. 10.1523/JNEUROSCI.23-28-09403.2003.

(86) Ananieva, O.; Darragh, J.; Johansen, C.; Carr, J. M.; McIlrath, J.; Park, J. M.; Wingate, A.; Monk, C. E.; Toth, R.; Santos, S. G.; Iversen, L.; Arthur, J. S. C. The Kinases MSK1 and MSK2 Act as Negative Regulators of Toll-like Receptor Signaling. Nat Immunol 2008, 9 (9), 1028–1036. 10.1038/ni.1644.

(87) Hu, X.; Paik, P. K.; Chen, J.; Yarilina, A.; Kockeritz, L.; Lu, T. T.; Woodgett, J. R.; Ivashkiv, L. B. IFN-γ Suppresses IL-10 Production and Synergizes with TLR2 by Regulating GSK3 and CREB/AP-1 Proteins. Immunity 2006, 24 (5), 563–574. 10.1016/j.immuni.2006.02.014.

(88) Wang, X.; Ni, L.; Chang, D.; Lu, H.; Jiang, Y.; Kim, B.-S.; Wang, A.; Liu, X.; Zhong, B.; Yang, X.; Dong, C. Cyclic AMP-Responsive Element-Binding Protein (CREB) Is Critical in Autoimmunity by Promoting Th17 but Inhibiting Treg Cell Differentiation. eBioMedicine 2017, 25, 165–174. 10.1016/j.ebiom.2017.10.010.

(89) Goel, R.; Bhat, S. A.; Hanif, K.; Nath, C.; Shukla, R. Angiotensin II Receptor Blockers Attenuate Lipopolysaccharide-Induced Memory Impairment by Modulation of NF-κB-Mediated BDNF/CREB Expression and Apoptosis in Spontaneously Hypertensive Rats. Mol Neurobiol 2018, 55 (2), 1725–1739. 10.1007/s12035-017-0450-5.

(90) Ma, W.; Quirion, R. Increased Phosphorylation of Cyclic AMP Response Element-Binding Protein (CREB) in the Superficial Dorsal Horn Neurons Following Partial Sciatic Nerve Ligation. Pain 2001, 93 (3), 295–301. 10.1016/S0304-3959(01)00335-9.

(91) Song, X.-S.; Cao, J.-L.; Xu, Y.-B.; He, J.-H.; Zhang, L.-C.; Zeng, Y.-M. Activation of ERK/CREB Pathway in Spinal Cord Contributes to Chronic Constrictive Injury-Induced Neuropathic Pain in Rats. Acta Pharmacol Sin 2005, 26 (7), 789–798. 10.1111/j.1745-7254.2005.00123.x.

(92) Martinez-Torres, S.; Mesquida-Veny, F.; Del Rio, J. A.; Hervera, A. Injury-Induced Activation of the Endocannabinoid System Promotes Axon Regeneration. iScience 2023, 26 (6), 106814. 10.1016/j.isci.2023.106814.

(93) Blázquez, C.; Chiarlone, A.; Bellocchio, L.; Resel, E.; Pruunsild, P.; García-Rincón, D.; Sendtner, M.; Timmusk, T.; Lutz, B.; Galve-Roperh, I.; Guzmán, M. The CB1 Cannabinoid Receptor Signals Striatal Neuroprotection via a PI3K/Akt/mTORC1/BDNF Pathway. Cell Death Differ 2015, 22 (10), 1618–1629. 10.1038/cdd.2015.11.

(94) Derkinderen, P.; Valjent, E.; Toutant, M.; Corvol, J.-C.; Enslen, H.; Ledent, C.; Trzaskos, J.; Caboche, J.; Girault, J.-A. Regulation of Extracellular Signal-Regulated Kinase by Cannabinoids in Hippocampus. J. Neurosci. 2003, 23 (6), 2371–2382. 10.1523/JNEUROSCI.23-06-02371.2003.

(95) Gupta, S.; Arnab, S.; Nguyen, K. L.; Reed, M.; Fathi, P.; Tammen, K.; Turner, E.; Jones, E.; Fischer, R.; Mendelowitz, D.; Bethea, J. R. Sex-Chromosome Complement and Activin-A Shape the Therapeutic Potential of TNFR2 Activation in a Model of MS and CNP. Proceedings of the National Academy of Sciences 2025, 122 (20), e2426771122. 10.1073/pnas.2426771122.

(96) Dellarole, A.; Morton, P.; Brambilla, R.; Walters, W.; Summers, S.; Bernardes, D.; Grilli, M.; Bethea, J. R. Neuropathic Pain-Induced Depressive-like Behavior and Hippocampal Neurogenesis and Plasticity Are Dependent on TNFR1 Signaling. Brain, Behavior, and Immunity 2014, 41, 65–81. 10.1016/j.bbi.2014.04.003.

