## Supplementary figure1-2 and table1-9 for "Hippocampal CA3 Nex/Neurod6^+^ neuron-specific TNFR2 alleviates chronic neuropathic pain by sex-dependently engaging opioid and endocannabinoid pathways"

**Supplementary Figures:**

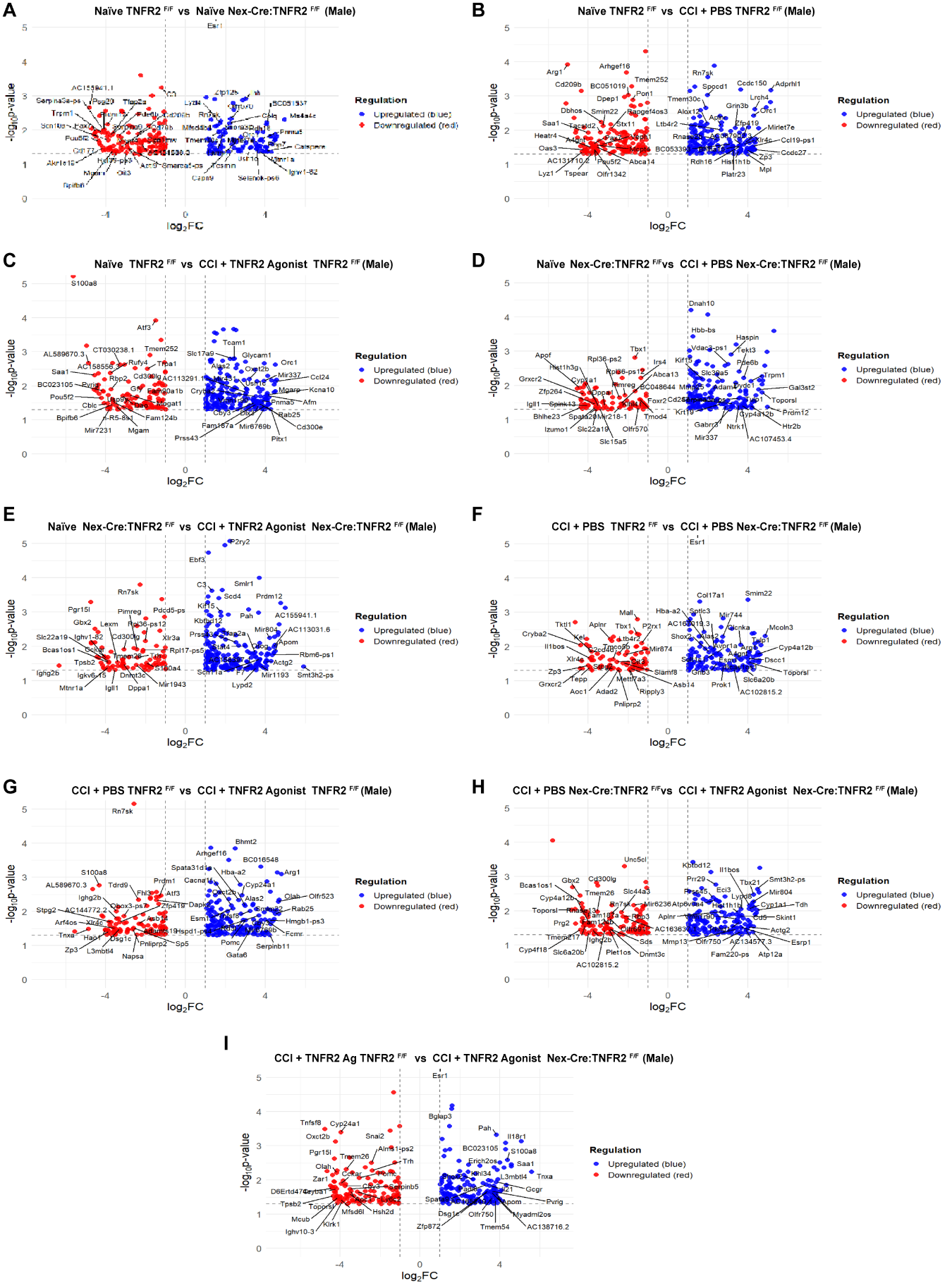

**Supplementary Figure 1:** Hippocampal bulk RNA sequencing data in male mice. Differential expression of protein-coding genes of **(A)** Naïve TNFR2^F/F^ vs Naïve Nex-Cre:TNFR2^F/F^ **(B)**  Naïve TNFR2^F/F^ vs CCI + PBS TNFR2^F/F^  **(C)** Naïve TNFR2^F/F^ vs CCI + TNFR2 Agonist TNFR2^F/F^  **(D)** Naïve Nex-Cre:TNFR2^F/F^ vs CCI + PBS Nex-Cre:TNFR2^F/F^ **(E)** Naïve Nex-Cre:TNFR2^F/F^ vs CCI + TNFR2 Agonist Nex-Cre:TNFR2^F/F^ **(F)** CCI + PBS TNFR2^F/F^ vs CCI + PBS Nex-Cre:TNFR2^F/F^ **(G)** CCI + PBS TNFR2^F/F^ vs CCI + TNFR2 Agonist TNFR2^F/F^ **(H)** CCI + PBS Nex-Cre:TNFR2^F/F^ vs CCI + TNFR2 Agonist Nex-Cre:TNFR2^F/F^ **(I)** CCI + TNFR2 Agonist TNFR2^F/F^ vs CCI + TNFR2 Agonist Nex-Cre:TNFR2^F/F^ (n=4-5 each group) (|log2 fold change| > 1).

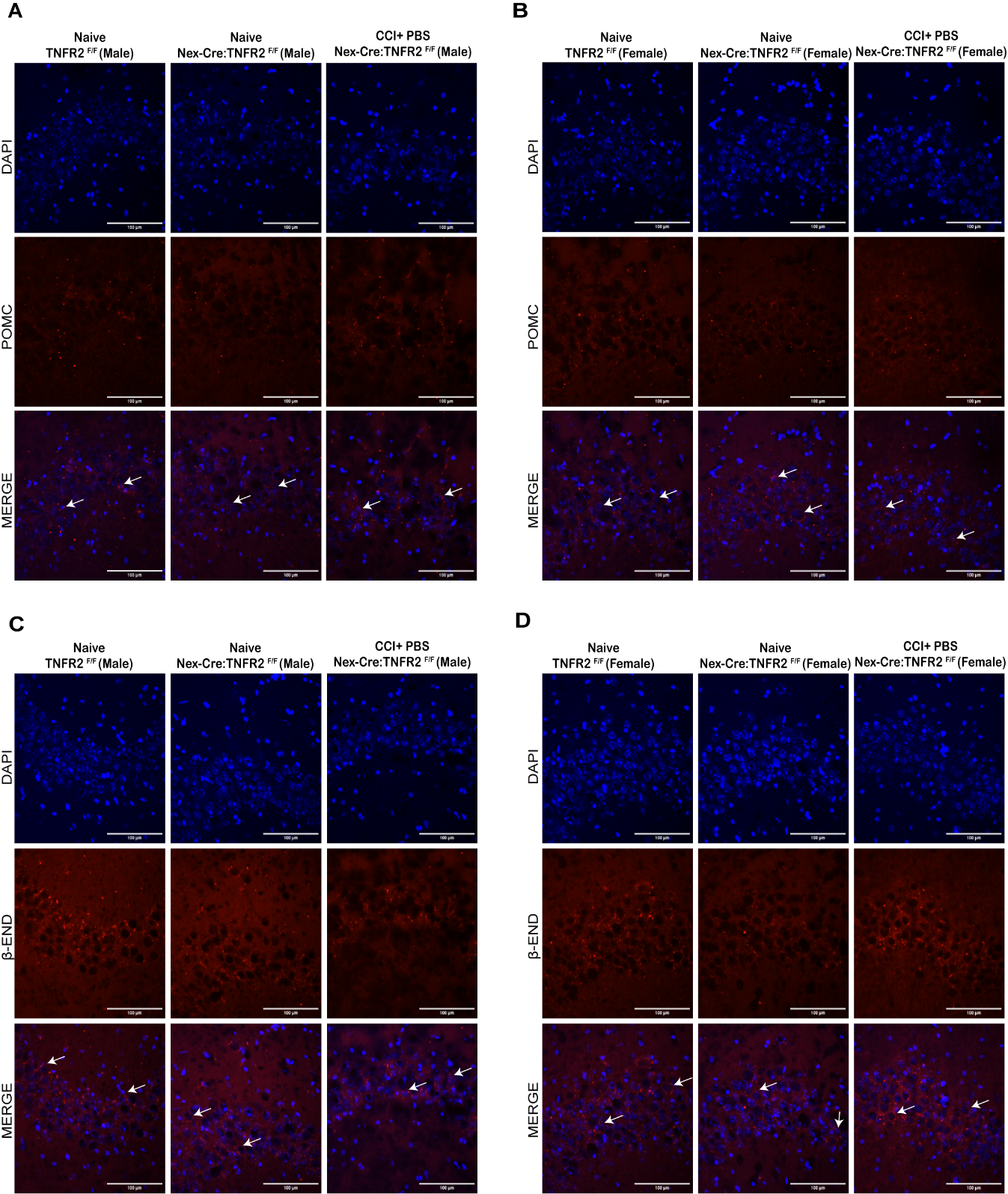

**Supplementary Figure 2:** Protein expression levels of POMC and β-endorphin in the CA3 region of the hippocampus in TNFR2^F/F^ (WT) Naïve mice, Nex-Cre:TNFR2^F/F^ Naïve mice, and Nex-Cre:TNFR2^F/F^ CCI + PBS mice. **(A)** and **(B)** POMC expression levels in the hippocampal CA3 region of male and female, represented by immunohistochemistry (n=4 each group/sex). **(C)** and **(D)** β-Endorphin expression in the hippocampal CA3 region of male and female, represented by immunohistochemistry (n=4 each group/sex). Data are represented as mean ± SEM; **P* < 0.05.

**Supplementary Tables: Differential expression of protein-coding genes**

**Supplementary Table 1: Naïve TNFR2^F/F^ vs Naïve Nex-Cre:TNFR2^F/F^**

| **S.No** | **Gene_ID** | **Gene_Name** | **Gene_Biotype** | **baseMean** | **log2FoldChange** | **lfcSE** | **stat** | **pvalue** | **padj** |
| --- | --- | --- | --- | --- | --- | --- | --- | --- | --- |
| 1 | ENSMUSG00000069044 | Usp9y | protein_coding | 3.22853 | -7.48649 | 3.792486 | -1.97403 | 0.048378 |  |
| 2 | ENSMUSG00000027857 | Tshb | protein_coding | 2.063315 | -6.8408 | 2.999867 | -2.28037 | 0.022586 |  |
| 3 | ENSMUSG00000093668 | Pou5f2 | protein_coding | 1.867564 | -6.68949 | 2.714223 | -2.46461 | 0.013716 |  |
| 4 | ENSMUSG00000096003 | Gm3500 | protein_coding | 1.78716 | -6.6329 | 2.722484 | -2.43634 | 0.014837 |  |
| 5 | ENSMUSG00000052212 | Cd177 | protein_coding | 1.723858 | -6.57491 | 2.737074 | -2.40217 | 0.016298 |  |
| 6 | ENSMUSG00000026415 | Fcamr | protein_coding | 1.624398 | -6.48816 | 2.908407 | -2.23083 | 0.025693 |  |
| 7 | ENSMUSG00000071551 | Akr1c19 | protein_coding | 1.599682 | -6.4653 | 2.753041 | -2.34842 | 0.018853 |  |
| 8 | ENSMUSG00000090894 | Olfr110 | protein_coding | 1.531868 | -6.41072 | 2.927175 | -2.19007 | 0.028519 |  |
| 9 | ENSMUSG00000047061 | Gm9817 | protein_coding | 1.462775 | -6.3392 | 2.944397 | -2.15297 | 0.031321 |  |
| 10 | ENSMUSG00000090176 | Cd200r2 | protein_coding | 1.395537 | -6.27704 | 2.964274 | -2.11756 | 0.034212 |  |
| 11 | ENSMUSG00000056148 | Rdh9 | protein_coding | 1.307489 | -6.17976 | 2.994427 | -2.06375 | 0.039041 |  |
| 12 | ENSMUSG00000030159 | Clec1b | protein_coding | 1.15671 | -6.0047 | 2.927839 | -2.0509 | 0.040277 |  |
| 13 | ENSMUSG00000052850 | Tas2r137 | protein_coding | 1.112604 | -5.95562 | 2.818229 | -2.11325 | 0.034579 |  |
| 14 | ENSMUSG00000075610 | Tmem92 | protein_coding | 1.090481 | -5.93007 | 2.934654 | -2.0207 | 0.04331 |  |
| 15 | ENSMUSG00000028736 | Pax7 | protein_coding | 3.440821 | -4.83216 | 2.032841 | -2.37705 | 0.017452 |  |
| 16 | ENSMUSG00000030523 | Trpm1 | protein_coding | 2.869012 | -4.47685 | 1.925784 | -2.32469 | 0.020089 |  |
| 17 | ENSMUSG00000023159 | Psg29 | protein_coding | 2.638036 | -4.33755 | 1.844707 | -2.35135 | 0.018705 |  |
| 18 | ENSMUSG00000034533 | Scn10a | protein_coding | 2.273461 | -4.20907 | 1.942391 | -2.16695 | 0.030238 |  |
| 19 | ENSMUSG00000026931 | 1700019N19Rik | protein_coding | 2.395182 | -4.14642 | 1.875049 | -2.21137 | 0.02701 |  |
| 20 | ENSMUSG00000079466 | Prdm12 | protein_coding | 4.10647 | -3.94388 | 1.461908 | -2.69776 | 0.006981 |  |
| 21 | ENSMUSG00000021359 | Tfap2a | protein_coding | 3.562927 | -3.73802 | 1.54547 | -2.41869 | 0.015576 |  |
| 22 | ENSMUSG00000029491 | Pde6b | protein_coding | 3.263388 | -3.60532 | 1.458143 | -2.47254 | 0.013416 |  |
| 23 | ENSMUSG00000061068 | Mcpt4 | protein_coding | 3.202876 | -3.58969 | 1.650417 | -2.17502 | 0.029629 |  |
| 24 | ENSMUSG00000058260 | Serpina9 | protein_coding | 4.305923 | -3.35237 | 1.362725 | -2.46005 | 0.013892 |  |
| 25 | ENSMUSG00000040592 | Cd79b | protein_coding | 2.543465 | -3.27245 | 1.527402 | -2.14249 | 0.032154 |  |
| 26 | ENSMUSG00000009654 | Oit3 | protein_coding | 3.134085 | -2.98179 | 1.483487 | -2.00999 | 0.044433 |  |
| 27 | ENSMUSG00000092519 | Actl9 | protein_coding | 2.962009 | -2.82849 | 1.337119 | -2.11536 | 0.034399 |  |
| 28 | ENSMUSG00000062939 | Stat4 | protein_coding | 3.334649 | -2.58161 | 1.302323 | -1.98231 | 0.047444 |  |
| 29 | ENSMUSG00000065987 | Cd209b | protein_coding | 5.680115 | -2.38029 | 0.947811 | -2.51136 | 0.012027 |  |
| 30 | ENSMUSG00000023279 | Bmp15 | protein_coding | 3.586106 | -2.34822 | 1.188916 | -1.9751 | 0.048257 |  |
| 31 | ENSMUSG00000045392 | Olfr1033 | protein_coding | 4.482342 | -2.33295 | 1.082272 | -2.15561 | 0.031114 |  |
| 32 | ENSMUSG00000074210 | E130208F15Rik | protein_coding | 4.201774 | -2.22507 | 1.042007 | -2.13537 | 0.032731 |  |
| 33 | ENSMUSG00000031394 | Opn1mw | protein_coding | 9.620842 | -2.15127 | 0.923694 | -2.32899 | 0.01986 |  |
| 34 | ENSMUSG00000039865 | Slc44a3 | protein_coding | 5.665176 | -2.0337 | 0.87276 | -2.3302 | 0.019796 |  |
| 35 | ENSMUSG00000085139 | A730046J19Rik | protein_coding | 11.2353 | -1.86446 | 0.686471 | -2.71601 | 0.006607 |  |
| 36 | ENSMUSG00000024670 | Cd6 | protein_coding | 10.48771 | -1.8416 | 0.790605 | -2.32935 | 0.01984 |  |
| 37 | ENSMUSG00000086564 | Cd101 | protein_coding | 7.155131 | -1.79404 | 0.821668 | -2.18341 | 0.029005 |  |
| 38 | ENSMUSG00000042379 | Esm1 | protein_coding | 5.66928 | -1.75921 | 0.88102 | -1.99679 | 0.045848 |  |
| 39 | ENSMUSG00000056071 | S100a9 | protein_coding | 14.48891 | -1.75281 | 0.805381 | -2.17638 | 0.029527 |  |
| 40 | ENSMUSG00000028523 | Tctex1d1 | protein_coding | 10.3392 | -1.6649 | 0.763046 | -2.18191 | 0.029116 |  |
| 41 | ENSMUSG00000060678 | Hist1h4c | protein_coding | 8.376771 | -1.64754 | 0.700169 | -2.35307 | 0.018619 |  |
| 42 | ENSMUSG00000009876 | Cox4i2 | protein_coding | 15.82553 | -1.58066 | 0.609568 | -2.59308 | 0.009512 |  |
| 43 | ENSMUSG00000027485 | Bpifb1 | protein_coding | 12.21236 | -1.49044 | 0.683312 | -2.18121 | 0.029168 |  |
| 44 | ENSMUSG00000061742 | Slc22a12 | protein_coding | 10.03337 | -1.4795 | 0.64802 | -2.2831 | 0.022424 |  |
| 45 | ENSMUSG00000032446 | Eomes | protein_coding | 12.66584 | -1.47074 | 0.61256 | -2.40097 | 0.016352 |  |
| 46 | ENSMUSG00000010830 | Kdelr3 | protein_coding | 17.98139 | -1.34582 | 0.536862 | -2.50684 | 0.012182 |  |
| 47 | ENSMUSG00000036768 | Kif15 | protein_coding | 10.88204 | -1.33194 | 0.615752 | -2.16311 | 0.030532 |  |
| 48 | ENSMUSG00000079391 | Gm2974 | protein_coding | 10.25034 | -1.27284 | 0.610141 | -2.08615 | 0.036965 |  |
| 49 | ENSMUSG00000104713 | Gbp6 | protein_coding | 20.61889 | -1.24066 | 0.516978 | -2.39983 | 0.016403 |  |
| 50 | ENSMUSG00000063594 | Gng8 | protein_coding | 17.768 | -1.23599 | 0.556913 | -2.21936 | 0.026462 |  |
| 51 | ENSMUSG00000024164 | C3 | protein_coding | 50.28645 | -1.20924 | 0.347731 | -3.47751 | 0.000506 |  |
| 52 | ENSMUSG00000033187 | BC016579 | protein_coding | 14.75688 | -1.15286 | 0.51328 | -2.24606 | 0.0247 |  |
| 53 | ENSMUSG00000032860 | P2ry2 | protein_coding | 14.23042 | -1.11747 | 0.568895 | -1.96428 | 0.049498 |  |
| 54 | ENSMUSG00000031022 | BC051019 | protein_coding | 13.51632 | -1.10401 | 0.520008 | -2.12307 | 0.033748 |  |
| 55 | ENSMUSG00000000686 | Abhd15 | protein_coding | 33.38401 | -1.10363 | 0.367743 | -3.00108 | 0.00269 |  |
| 56 | ENSMUSG00000028558 | Calr4 | protein_coding | 25.35285 | -1.09707 | 0.455793 | -2.40696 | 0.016086 |  |
| 57 | ENSMUSG00000054320 | Lrrc36 | protein_coding | 40.44135 | -1.02676 | 0.364461 | -2.81721 | 0.004844 |  |
| 58 | ENSMUSG00000017861 | Mybl2 | protein_coding | 21.31283 | -1.02658 | 0.472843 | -2.17108 | 0.029925 |  |
| 59 | ENSMUSG00000032643 | Fhl3 | protein_coding | 18.38785 | -1.00626 | 0.447906 | -2.24658 | 0.024667 |  |
| 60 | ENSMUSG00000000301 | Pemt | protein_coding | 47.1728 | 1.034851 | 0.314184 | 3.293774 | 0.000989 |  |
| 61 | ENSMUSG00000089798 | 1700028K03Rik | protein_coding | 20.07675 | 1.179165 | 0.558019 | 2.113126 | 0.03459 |  |
| 62 | ENSMUSG00000037705 | Tecta | protein_coding | 26.56629 | 1.186866 | 0.452241 | 2.624411 | 0.00868 |  |
| 63 | ENSMUSG00000033114 | Slc35d2 | protein_coding | 17.92414 | 1.192257 | 0.579308 | 2.058072 | 0.039583 |  |
| 64 | ENSMUSG00000020447 | Npc1l1 | protein_coding | 18.90418 | 1.199056 | 0.510799 | 2.347412 | 0.018904 |  |
| 65 | ENSMUSG00000079364 | Gm3558 | protein_coding | 15.30577 | 1.20243 | 0.543109 | 2.213975 | 0.02683 |  |
| 66 | ENSMUSG00000100916 | Lhb | protein_coding | 13.80104 | 1.319255 | 0.596529 | 2.21155 | 0.026998 |  |
| 67 | ENSMUSG00000019768 | Esr1 | protein_coding | 311.0309 | 1.516727 | 0.179747 | 8.438141 | 3.22423989672433e-17 | 2.23391461244545e-13 |
| 68 | ENSMUSG00000053268 | Dspp | protein_coding | 6.918622 | 1.553088 | 0.741033 | 2.095843 | 0.036096 |  |
| 69 | ENSMUSG00000038522 | Mfsd4b1 | protein_coding | 14.09228 | 1.589235 | 0.586043 | 2.711804 | 0.006692 |  |
| 70 | ENSMUSG00000005220 | Corin | protein_coding | 8.984903 | 1.635082 | 0.810043 | 2.018513 | 0.043538 |  |
| 71 | ENSMUSG00000022753 | Tmem30c | protein_coding | 16.58186 | 1.640693 | 0.642243 | 2.554629 | 0.01063 |  |
| 72 | ENSMUSG00000043664 | Tmem221 | protein_coding | 8.144278 | 1.691263 | 0.854043 | 1.980301 | 0.04767 |  |
| 73 | ENSMUSG00000002930 | Ppp1r17 | protein_coding | 12.51151 | 1.709637 | 0.703387 | 2.430578 | 0.015075 |  |
| 74 | ENSMUSG00000032530 | Lyzl4 | protein_coding | 12.15639 | 1.731673 | 0.625852 | 2.766906 | 0.005659 |  |
| 75 | ENSMUSG00000075405 | 9430097D07Rik | protein_coding | 7.949264 | 1.761434 | 0.802022 | 2.196242 | 0.028075 |  |
| 76 | ENSMUSG00000068697 | Myoz1 | protein_coding | 6.912871 | 1.791155 | 0.878935 | 2.037871 | 0.041563 |  |
| 77 | ENSMUSG00000045106 | Ccdc73 | protein_coding | 7.508711 | 1.861092 | 0.79688 | 2.335474 | 0.019519 |  |
| 78 | ENSMUSG00000053338 | Tarm1 | protein_coding | 7.564703 | 1.88681 | 0.754306 | 2.501387 | 0.012371 |  |
| 79 | ENSMUSG00000057606 | Colq | protein_coding | 8.544756 | 2.209412 | 0.801731 | 2.7558 | 0.005855 |  |
| 80 | ENSMUSG00000073964 | Olfr570 | protein_coding | 13.63276 | 2.263268 | 0.790594 | 2.862742 | 0.0042 |  |
| 81 | ENSMUSG00000024905 | Tesmin | protein_coding | 5.080516 | 2.445941 | 1.140466 | 2.144686 | 0.031978 |  |
| 82 | ENSMUSG00000031981 | Capn9 | protein_coding | 5.24765 | 2.507681 | 1.086571 | 2.307886 | 0.021005 |  |
| 83 | ENSMUSG00000033255 | Gm5134 | protein_coding | 3.04755 | 2.838761 | 1.334292 | 2.127541 | 0.033375 |  |
| 84 | ENSMUSG00000006538 | Ihh | protein_coding | 6.756938 | 3.002336 | 0.922023 | 3.256248 | 0.001129 |  |
| 85 | ENSMUSG00000030838 | Ush1c | protein_coding | 2.717604 | 3.258326 | 1.520224 | 2.14332 | 0.032087 |  |
| 86 | ENSMUSG00000023120 | Gm853 | protein_coding | 3.818708 | 3.909817 | 1.543545 | 2.533011 | 0.011309 |  |
| 87 | ENSMUSG00000028587 | Orc1 | protein_coding | 2.126571 | 4.031168 | 1.885726 | 2.137728 | 0.032539 |  |
| 88 | ENSMUSG00000050424 | Pnma5 | protein_coding | 2.260604 | 4.043965 | 1.875598 | 2.156093 | 0.031076 |  |
| 89 | ENSMUSG00000028996 | Rbp7 | protein_coding | 2.394244 | 4.134629 | 1.934 | 2.137864 | 0.032528 |  |
| 90 | ENSMUSG00000069456 | Rdh16 | protein_coding | 2.678583 | 4.325783 | 1.792638 | 2.413083 | 0.015818 |  |
| 91 | ENSMUSG00000049291 | Prss38 | protein_coding | 1.093996 | 5.893312 | 2.92951 | 2.011706 | 0.044251 |  |
| 92 | ENSMUSG00000006389 | Mpl | protein_coding | 1.244659 | 6.053464 | 2.764091 | 2.190038 | 0.028522 |  |
| 93 | ENSMUSG00000029369 | Afm | protein_coding | 1.283109 | 6.075635 | 2.923528 | 2.078186 | 0.037692 |  |
| 94 | ENSMUSG00000112611 | Glipr1l3 | protein_coding | 1.328544 | 6.169754 | 2.851221 | 2.163899 | 0.030472 |  |
| 95 | ENSMUSG00000037161 | Mgarp | protein_coding | 1.391366 | 6.192141 | 2.79316 | 2.216895 | 0.02663 |  |
| 96 | ENSMUSG00000024770 | Lipn | protein_coding | 1.378868 | 6.204001 | 2.994135 | 2.072051 | 0.038261 |  |
| 97 | ENSMUSG00000016758 | Bik | protein_coding | 1.376493 | 6.216089 | 2.804242 | 2.216673 | 0.026645 |  |
| 98 | ENSMUSG00000037053 | Azgp1 | protein_coding | 1.404164 | 6.228298 | 3.015067 | 2.065724 | 0.038855 |  |
| 99 | ENSMUSG00000047592 | Nxpe5 | protein_coding | 1.406879 | 6.254936 | 2.882164 | 2.170222 | 0.02999 |  |
| 100 | ENSMUSG00000054764 | Mtnr1a | protein_coding | 1.84639 | 6.61613 | 2.85974 | 2.313543 | 0.020693 |  |
| 101 | ENSMUSG00000024675 | Ms4a4c | protein_coding | 2.867856 | 7.270923 | 2.645731 | 2.748172 | 0.005993 |  |

**Supplementary Table 2: Naïve vs CCI + PBS TNFR2^F/F^**

| **S.No** | **Gene_ID** | **Gene_Name** | **Gene_Biotype** | **baseMean** | **log2FoldChange** | **lfcSE** | **stat** | **pvalue** | **padj** |
| --- | --- | --- | --- | --- | --- | --- | --- | --- | --- |
| 1 | ENSMUSG00000069044 | Usp9y | protein_coding | 3.265602 | -7.51953 | 3.792628 | -1.98267 | 0.047404 |  |
| 2 | ENSMUSG00000069515 | Lyz1 | protein_coding | 1.771955 | -6.63067 | 2.859014 | -2.31922 | 0.020383 |  |
| 3 | ENSMUSG00000032661 | Oas3 | protein_coding | 1.625336 | -6.53017 | 2.797161 | -2.33457 | 0.019566 |  |
| 4 | ENSMUSG00000110266 | Gm32742 | protein_coding | 1.470768 | -6.36248 | 2.781561 | -2.28738 | 0.022174 |  |
| 5 | ENSMUSG00000069581 | Tspear | protein_coding | 1.266984 | -6.1573 | 2.738547 | -2.24838 | 0.024552 |  |
| 6 | ENSMUSG00000039508 | Fam26d | protein_coding | 1.190919 | -6.07664 | 3.014094 | -2.01608 | 0.043792 |  |
| 7 | ENSMUSG00000040485 | Lrrc52 | protein_coding | 1.203701 | -6.07162 | 2.812096 | -2.15911 | 0.030842 |  |
| 8 | ENSMUSG00000046774 | 8030474K03Rik | protein_coding | 1.197037 | -6.06779 | 3.018882 | -2.00995 | 0.044437 |  |
| 9 | ENSMUSG00000022832 | Ropn1 | protein_coding | 1.191814 | -6.0621 | 3.066893 | -1.97663 | 0.048084 |  |
| 10 | ENSMUSG00000031603 | Fgf20 | protein_coding | 1.167658 | -6.05049 | 3.042395 | -1.98873 | 0.046731 |  |
| 11 | ENSMUSG00000048981 | Krt31 | protein_coding | 1.167658 | -6.05049 | 3.042395 | -1.98873 | 0.046731 |  |
| 12 | ENSMUSG00000046623 | Gjb4 | protein_coding | 1.178733 | -6.04824 | 3.071533 | -1.96913 | 0.048939 |  |
| 13 | ENSMUSG00000090509 | Sfta2 | protein_coding | 1.029321 | -5.85485 | 2.981331 | -1.96384 | 0.049549 |  |
| 14 | ENSMUSG00000004872 | Pax3 | protein_coding | 1.024099 | -5.84796 | 2.93286 | -1.99394 | 0.046158 |  |
| 15 | ENSMUSG00000019987 | Arg1 | protein_coding | 4.97751 | -5.29926 | 1.633563 | -3.24399 | 0.001179 |  |
| 16 | ENSMUSG00000074115 | Saa1 | protein_coding | 2.472013 | -4.39811 | 1.899183 | -2.31579 | 0.02057 |  |
| 17 | ENSMUSG00000065987 | Cd209b | protein_coding | 5.062624 | -4.39227 | 1.314885 | -3.34042 | 0.000837 |  |
| 18 | ENSMUSG00000090843 | Heatr4 | protein_coding | 2.205336 | -4.25559 | 2.074146 | -2.05173 | 0.040196 |  |
| 19 | ENSMUSG00000037953 | A4gnt | protein_coding | 2.299086 | -4.07474 | 1.939777 | -2.10062 | 0.035674 |  |
| 20 | ENSMUSG00000043383 | Olfr1342 | protein_coding | 2.021247 | -3.99468 | 2.005776 | -1.99159 | 0.046416 |  |
| 21 | ENSMUSG00000062017 | Abca14 | protein_coding | 1.897458 | -3.89748 | 1.921742 | -2.0281 | 0.04255 |  |
| 22 | ENSMUSG00000093668 | Pou5f2 | protein_coding | 2.035419 | -3.88049 | 1.972829 | -1.96697 | 0.049187 |  |
| 23 | ENSMUSG00000028736 | Pax7 | protein_coding | 3.651171 | -3.70275 | 1.6629 | -2.22668 | 0.025969 |  |
| 24 | ENSMUSG00000061068 | Mcpt4 | protein_coding | 3.23094 | -3.61829 | 1.63919 | -2.20736 | 0.027289 |  |
| 25 | ENSMUSG00000051397 | Tacstd2 | protein_coding | 4.085398 | -3.39613 | 1.33346 | -2.54686 | 0.01087 |  |
| 26 | ENSMUSG00000027831 | Veph1 | protein_coding | 2.789598 | -3.30052 | 1.491508 | -2.21288 | 0.026906 |  |
| 27 | ENSMUSG00000096215 | Smim22 | protein_coding | 3.843143 | -3.2627 | 1.204046 | -2.70978 | 0.006733 |  |
| 28 | ENSMUSG00000068860 | Gm128 | protein_coding | 3.937808 | -3.25348 | 1.476907 | -2.2029 | 0.027602 |  |
| 29 | ENSMUSG00000049709 | Nlrp10 | protein_coding | 3.617967 | -3.17702 | 1.4244 | -2.23043 | 0.025719 |  |
| 30 | ENSMUSG00000050108 | Bpifc | protein_coding | 2.540951 | -3.14671 | 1.505799 | -2.08973 | 0.036642 |  |
| 31 | ENSMUSG00000040694 | Apobec2 | protein_coding | 3.564838 | -2.70675 | 1.314713 | -2.05881 | 0.039512 |  |
| 32 | ENSMUSG00000039232 | Stx11 | protein_coding | 5.609596 | -2.70639 | 1.044949 | -2.58997 | 0.009598 |  |
| 33 | ENSMUSG00000057933 | Gsta2 | protein_coding | 4.015283 | -2.53222 | 1.130935 | -2.23905 | 0.025153 |  |
| 34 | ENSMUSG00000061815 | Rufy4 | protein_coding | 5.026216 | -2.50876 | 1.164781 | -2.15385 | 0.031252 |  |
| 35 | ENSMUSG00000049537 | Tecrl | protein_coding | 6.103279 | -2.41117 | 1.027431 | -2.34679 | 0.018936 |  |
| 36 | ENSMUSG00000042379 | Esm1 | protein_coding | 5.375417 | -2.24742 | 0.9126 | -2.46266 | 0.013791 |  |
| 37 | ENSMUSG00000030666 | Calcb | protein_coding | 4.430098 | -2.20661 | 1.035967 | -2.13 | 0.033172 |  |
| 38 | ENSMUSG00000033427 | Upb1 | protein_coding | 5.08869 | -2.19251 | 1.063206 | -2.06216 | 0.039192 |  |
| 39 | ENSMUSG00000019278 | Dpep1 | protein_coding | 12.96937 | -2.18825 | 0.661038 | -3.31032 | 0.000932 |  |
| 40 | ENSMUSG00000023439 | Gnb3 | protein_coding | 4.821104 | -2.16719 | 1.101134 | -1.96815 | 0.049051 |  |
| 41 | ENSMUSG00000056987 | Fam71d | protein_coding | 6.957411 | -2.16174 | 0.929892 | -2.32472 | 0.020087 |  |
| 42 | ENSMUSG00000030116 | Mfap5 | protein_coding | 7.445083 | -2.14896 | 0.909568 | -2.36262 | 0.018146 |  |
| 43 | ENSMUSG00000029032 | Arhgef16 | protein_coding | 15.6694 | -2.10011 | 0.558812 | -3.75816 | 0.000171 |  |
| 44 | ENSMUSG00000030911 | Zp2 | protein_coding | 8.506352 | -2.02934 | 0.814201 | -2.49243 | 0.012687 |  |
| 45 | ENSMUSG00000031022 | BC051019 | protein_coding | 11.74449 | -2.00429 | 0.595789 | -3.3641 | 0.000768 |  |
| 46 | ENSMUSG00000023943 | Sult1c1 | protein_coding | 8.928138 | -1.9478 | 0.91588 | -2.12669 | 0.033446 |  |
| 47 | ENSMUSG00000046223 | Plaur | protein_coding | 6.830028 | -1.94037 | 0.885854 | -2.19039 | 0.028496 |  |
| 48 | ENSMUSG00000054988 | Agtr1b | protein_coding | 5.528563 | -1.89946 | 0.96317 | -1.97209 | 0.0486 |  |
| 49 | ENSMUSG00000055333 | Fat2 | protein_coding | 17.85495 | -1.84915 | 0.625382 | -2.95684 | 0.003108 |  |
| 50 | ENSMUSG00000041301 | Cftr | protein_coding | 15.05976 | -1.84407 | 0.586533 | -3.14403 | 0.001666 |  |
| 51 | ENSMUSG00000048572 | Tmem252 | protein_coding | 19.53552 | -1.83282 | 0.522349 | -3.50881 | 0.00045 |  |
| 52 | ENSMUSG00000002588 | Pon1 | protein_coding | 23.55797 | -1.77407 | 0.548642 | -3.23356 | 0.001223 |  |
| 53 | ENSMUSG00000040380 | Cbln3 | protein_coding | 21.13877 | -1.71181 | 0.551693 | -3.10283 | 0.001917 |  |
| 54 | ENSMUSG00000039092 | Sptlc3 | protein_coding | 16.2814 | -1.63047 | 0.555659 | -2.9343 | 0.003343 |  |
| 55 | ENSMUSG00000040950 | Mgl2 | protein_coding | 10.80223 | -1.62475 | 0.646393 | -2.51357 | 0.011952 |  |
| 56 | ENSMUSG00000025804 | Ccr1 | protein_coding | 9.935442 | -1.53984 | 0.702376 | -2.19233 | 0.028356 |  |
| 57 | ENSMUSG00000085139 | A730046J19Rik | protein_coding | 12.27928 | -1.4625 | 0.683851 | -2.13862 | 0.032466 |  |
| 58 | ENSMUSG00000037868 | Egr2 | protein_coding | 81.90663 | -1.44646 | 0.667308 | -2.16761 | 0.030188 | 0.446146493 |
| 59 | ENSMUSG00000025064 | Col17a1 | protein_coding | 24.18426 | -1.38009 | 0.437831 | -3.15211 | 0.001621 |  |
| 60 | ENSMUSG00000010830 | Kdelr3 | protein_coding | 18.04851 | -1.34412 | 0.521958 | -2.57516 | 0.010019 |  |
| 61 | ENSMUSG00000032725 | Folr2 | protein_coding | 13.07757 | -1.32698 | 0.617246 | -2.14984 | 0.031568 |  |
| 62 | ENSMUSG00000015401 | Tmem27 | protein_coding | 14.56599 | -1.31602 | 0.562438 | -2.33985 | 0.019292 |  |
| 63 | ENSMUSG00000035165 | Kcne3 | protein_coding | 9.25484 | -1.31321 | 0.639364 | -2.05393 | 0.039983 |  |
| 64 | ENSMUSG00000019787 | Trdn | protein_coding | 12.36283 | -1.29254 | 0.638356 | -2.02479 | 0.042889 |  |
| 65 | ENSMUSG00000064356 | mt-Atp8 | protein_coding | 14.92781 | -1.27611 | 0.564468 | -2.26073 | 0.023776 |  |
| 66 | ENSMUSG00000006642 | Tcf23 | protein_coding | 11.97069 | -1.25448 | 0.572972 | -2.18942 | 0.028566 |  |
| 67 | ENSMUSG00000079941 | Gm11273 | protein_coding | 21.14841 | -1.21276 | 0.495914 | -2.44551 | 0.014465 |  |
| 68 | ENSMUSG00000071656 | Lrrn4cl | protein_coding | 24.41321 | -1.16634 | 0.460061 | -2.5352 | 0.011238 |  |
| 69 | ENSMUSG00000003355 | Fkbp11 | protein_coding | 15.57385 | -1.15892 | 0.552511 | -2.09755 | 0.035945 |  |
| 70 | ENSMUSG00000019368 | Sec14l4 | protein_coding | 30.1841 | -1.13803 | 0.400047 | -2.84474 | 0.004445 |  |
| 71 | ENSMUSG00000028195 | Cyr61 | protein_coding | 157.5505 | -1.13287 | 0.279487 | -4.05338 | 5.05E-05 | 0.026936662 |
| 72 | ENSMUSG00000069899 | Gm12166 | protein_coding | 15.57905 | -1.12534 | 0.542578 | -2.07406 | 0.038074 |  |
| 73 | ENSMUSG00000021250 | Fos | protein_coding | 546.0452 | -1.08269 | 0.412075 | -2.62741 | 0.008604 | 0.277309115 |
| 74 | ENSMUSG00000022032 | Scara5 | protein_coding | 70.85181 | -1.04298 | 0.328458 | -3.17539 | 0.001496 | 0.13519811 |
| 75 | ENSMUSG00000051079 | Rgs13 | protein_coding | 21.27396 | -1.00742 | 0.479954 | -2.099 | 0.035817 |  |
| 76 | ENSMUSG00000052631 | Sh2d6 | protein_coding | 14.4072 | 1.052082 | 0.512983 | 2.050912 | 0.040275 |  |
| 77 | ENSMUSG00000034881 | Tbxa2r | protein_coding | 21.29304 | 1.066068 | 0.436976 | 2.439651 | 0.014701 |  |
| 78 | ENSMUSG00000045102 | Poln | protein_coding | 17.61988 | 1.069966 | 0.485802 | 2.202476 | 0.027632 |  |
| 79 | ENSMUSG00000025592 | Dach2 | protein_coding | 27.66219 | 1.088422 | 0.472843 | 2.301867 | 0.021343 |  |
| 80 | ENSMUSG00000033114 | Slc35d2 | protein_coding | 17.2821 | 1.091134 | 0.546672 | 1.995957 | 0.045939 |  |
| 81 | ENSMUSG00000091568 | Gm8206 | protein_coding | 20.91101 | 1.173065 | 0.441688 | 2.655869 | 0.00791 |  |
| 82 | ENSMUSG00000094152 | Slc6a16 | protein_coding | 17.8134 | 1.18338 | 0.580815 | 2.037448 | 0.041605 |  |
| 83 | ENSMUSG00000078157 | 4931440F15Rik | protein_coding | 10.53694 | 1.240248 | 0.625788 | 1.981896 | 0.047491 |  |
| 84 | ENSMUSG00000021768 | Dusp13 | protein_coding | 13.76048 | 1.269242 | 0.562547 | 2.256243 | 0.024055 |  |
| 85 | ENSMUSG00000020447 | Npc1l1 | protein_coding | 20.17796 | 1.305629 | 0.482185 | 2.707733 | 0.006774 |  |
| 86 | ENSMUSG00000074635 | 3110070M22Rik | protein_coding | 24.20167 | 1.347459 | 0.527569 | 2.554091 | 0.010647 |  |
| 87 | ENSMUSG00000079105 | C7 | protein_coding | 15.36247 | 1.374423 | 0.603187 | 2.278602 | 0.022691 |  |
| 88 | ENSMUSG00000046110 | Serinc4 | protein_coding | 11.84833 | 1.380905 | 0.661946 | 2.086132 | 0.036967 |  |
| 89 | ENSMUSG00000037661 | Gpr160 | protein_coding | 17.20026 | 1.435361 | 0.594836 | 2.413036 | 0.01582 |  |
| 90 | ENSMUSG00000100235 | Gm28557 | protein_coding | 11.32807 | 1.48434 | 0.61618 | 2.408938 | 0.015999 |  |
| 91 | ENSMUSG00000038522 | Mfsd4b1 | protein_coding | 14.26095 | 1.59589 | 0.59217 | 2.694987 | 0.007039 |  |
| 92 | ENSMUSG00000069041 | Slc25a31 | protein_coding | 6.759413 | 1.616746 | 0.803297 | 2.012638 | 0.044153 |  |
| 93 | ENSMUSG00000040367 | Lrrd1 | protein_coding | 7.418194 | 1.632568 | 0.813763 | 2.006196 | 0.044835 |  |
| 94 | ENSMUSG00000074213 | Gm10642 | protein_coding | 6.949076 | 1.688835 | 0.808483 | 2.088895 | 0.036717 |  |
| 95 | ENSMUSG00000022034 | Esco2 | protein_coding | 5.982908 | 1.6917 | 0.856808 | 1.974423 | 0.048334 |  |
| 96 | ENSMUSG00000021499 | Catsper3 | protein_coding | 7.233957 | 1.772773 | 0.769927 | 2.302522 | 0.021306 |  |
| 97 | ENSMUSG00000034127 | Tspan8 | protein_coding | 6.594142 | 1.79097 | 0.839292 | 2.133907 | 0.03285 |  |
| 98 | ENSMUSG00000031312 | Itgb1bp2 | protein_coding | 9.995179 | 1.826855 | 0.699972 | 2.609898 | 0.009057 |  |
| 99 | ENSMUSG00000053368 | Rxfp2 | protein_coding | 7.512742 | 1.846735 | 0.811543 | 2.275586 | 0.022871 |  |
| 100 | ENSMUSG00000022753 | Tmem30c | protein_coding | 19.2422 | 1.889607 | 0.614379 | 3.075638 | 0.002101 |  |
| 101 | ENSMUSG00000000320 | Alox12 | protein_coding | 10.16163 | 1.955672 | 0.709526 | 2.756308 | 0.005846 |  |
| 102 | ENSMUSG00000028784 | Spocd1 | protein_coding | 11.80218 | 1.95882 | 0.583317 | 3.358073 | 0.000785 |  |
| 103 | ENSMUSG00000057606 | Colq | protein_coding | 7.590945 | 1.996085 | 0.772533 | 2.583817 | 0.009771 |  |
| 104 | ENSMUSG00000030577 | Cd22 | protein_coding | 4.57811 | 1.996357 | 1.007572 | 1.981355 | 0.047551 |  |
| 105 | ENSMUSG00000020712 | Tcam1 | protein_coding | 7.269612 | 2.122428 | 0.899191 | 2.360377 | 0.018256 |  |
| 106 | ENSMUSG00000020698 | Cct6b | protein_coding | 5.233362 | 2.251172 | 1.116194 | 2.016828 | 0.043713 |  |
| 107 | ENSMUSG00000022491 | Glycam1 | protein_coding | 4.639775 | 2.40837 | 1.151616 | 2.091296 | 0.036502 |  |
| 108 | ENSMUSG00000059606 | Rnase2b | protein_coding | 4.16585 | 2.473831 | 1.038547 | 2.382011 | 0.017218 |  |
| 109 | ENSMUSG00000091648 | C2cd4d | protein_coding | 4.154582 | 2.487093 | 1.266105 | 1.964366 | 0.049488 |  |
| 110 | ENSMUSG00000021898 | Asb14 | protein_coding | 4.307577 | 2.492007 | 1.114237 | 2.236514 | 0.025318 |  |
| 111 | ENSMUSG00000040432 | Ltb4r2 | protein_coding | 4.744991 | 2.658567 | 1.014996 | 2.619289 | 0.008811 |  |
| 112 | ENSMUSG00000044338 | Aplnr | protein_coding | 6.370912 | 2.662004 | 0.985736 | 2.700524 | 0.006923 |  |
| 113 | ENSMUSG00000073492 | Gm10521 | protein_coding | 5.674874 | 2.821242 | 1.137546 | 2.480113 | 0.013134 |  |
| 114 | ENSMUSG00000046974 | BC053393 | protein_coding | 3.72833 | 3.120298 | 1.388728 | 2.246875 | 0.024648 |  |
| 115 | ENSMUSG00000035745 | Grin3b | protein_coding | 4.262618 | 3.360936 | 1.267815 | 2.650968 | 0.008026 |  |
| 116 | ENSMUSG00000025983 | Ccdc150 | protein_coding | 6.892456 | 3.642907 | 1.06077 | 3.434209 | 0.000594 |  |
| 117 | ENSMUSG00000069456 | Rdh16 | protein_coding | 2.030678 | 3.869611 | 1.907847 | 2.028261 | 0.042534 |  |
| 118 | ENSMUSG00000052271 | Bhlha15 | protein_coding | 1.996388 | 3.874432 | 1.902905 | 2.036062 | 0.041744 |  |
| 119 | ENSMUSG00000031362 | Xlr4c | protein_coding | 2.281041 | 4.04341 | 1.945076 | 2.078792 | 0.037636 |  |
| 120 | ENSMUSG00000028587 | Orc1 | protein_coding | 3.158016 | 4.601288 | 1.789635 | 2.571076 | 0.010138 |  |
| 121 | ENSMUSG00000093445 | Lrch4 | protein_coding | 3.298281 | 4.60513 | 1.730426 | 2.66127 | 0.007785 |  |
| 122 | ENSMUSG00000057465 | Saa2 | protein_coding | 0.962946 | 5.673247 | 2.884723 | 1.966653 | 0.049223 |  |
| 123 | ENSMUSG00000065952 | C330021F23Rik | protein_coding | 1.072813 | 5.844779 | 2.935821 | 1.99085 | 0.046497 |  |
| 124 | ENSMUSG00000097025 | Gm26558 | protein_coding | 1.135513 | 5.903249 | 2.922313 | 2.02006 | 0.043377 |  |
| 125 | ENSMUSG00000032292 | Nr2e3 | protein_coding | 1.166691 | 5.950563 | 2.890422 | 2.058718 | 0.039521 |  |
| 126 | ENSMUSG00000068604 | Gm8225 | protein_coding | 1.203051 | 6.020872 | 2.918354 | 2.063105 | 0.039103 |  |
| 127 | ENSMUSG00000042861 | Kcna10 | protein_coding | 1.249669 | 6.039384 | 3.002141 | 2.011693 | 0.044252 |  |
| 128 | ENSMUSG00000031204 | Asb12 | protein_coding | 1.266162 | 6.045166 | 3.049518 | 1.982335 | 0.047442 |  |
| 129 | ENSMUSG00000115768 | DPEP2NB | protein_coding | 1.301095 | 6.097747 | 3.007253 | 2.02768 | 0.042593 |  |
| 130 | ENSMUSG00000057135 | Scimp | protein_coding | 1.333288 | 6.170787 | 3.014373 | 2.047121 | 0.040646 |  |
| 131 | ENSMUSG00000024837 | Dmrt1 | protein_coding | 1.388074 | 6.193726 | 2.834323 | 2.185257 | 0.02887 |  |
| 132 | ENSMUSG00000004948 | Zp3 | protein_coding | 1.443534 | 6.270349 | 2.717349 | 2.307524 | 0.021026 |  |
| 133 | ENSMUSG00000016758 | Bik | protein_coding | 1.491457 | 6.283097 | 2.814206 | 2.232636 | 0.025573 |  |
| 134 | ENSMUSG00000006389 | Mpl | protein_coding | 1.501052 | 6.291199 | 2.795491 | 2.250481 | 0.024418 |  |
| 135 | ENSMUSG00000039492 | Ccdc27 | protein_coding | 1.912418 | 6.659602 | 2.830636 | 2.352687 | 0.018638 |  |
| 136 | ENSMUSG00000031448 | Adprhl1 | protein_coding | 3.315016 | 7.469102 | 2.511181 | 2.974339 | 0.002936 |  |

**Supplementary Table 3: Naïve vs CCI + TNFR2 Ag TNFR2^F/F^**

| **S.No** | **Gene_ID** | **Gene_Name** | **Gene_Biotype** | **baseMean** | **log2FoldChange** | **lfcSE** | **stat** | **pvalue** | **padj** |
| --- | --- | --- | --- | --- | --- | --- | --- | --- | --- |
| 1 | ENSMUSG00000074115 | Saa1 | protein_coding | 1.981737 | -6.88289 | 2.383977 | -2.88714 | 0.003888 |  |
| 2 | ENSMUSG00000093668 | Pou5f2 | protein_coding | 1.573324 | -6.54732 | 2.516522 | -2.60173 | 0.009275 |  |
| 3 | ENSMUSG00000040525 | Cblc | protein_coding | 1.257413 | -6.23478 | 2.502345 | -2.49158 | 0.012718 |  |
| 4 | ENSMUSG00000087610 | Gm16253 | protein_coding | 1.16228 | -6.11977 | 2.522748 | -2.42584 | 0.015273 |  |
| 5 | ENSMUSG00000042477 | Tfap2e | protein_coding | 1.147935 | -6.09349 | 3.020965 | -2.01707 | 0.043689 |  |
| 6 | ENSMUSG00000056054 | S100a8 | protein_coding | 6.35244 | -6.08518 | 1.655164 | -3.67648 | 0.000236 |  |
| 7 | ENSMUSG00000039508 | Fam26d | protein_coding | 0.991731 | -5.89904 | 2.867003 | -2.05756 | 0.039632 |  |
| 8 | ENSMUSG00000028786 | Tmem54 | protein_coding | 0.976468 | -5.86319 | 2.595646 | -2.25886 | 0.023892 |  |
| 9 | ENSMUSG00000021622 | Ckmt2 | protein_coding | 0.895138 | -5.75067 | 2.929686 | -1.96289 | 0.049658 |  |
| 10 | ENSMUSG00000084897 | Gm14226 | protein_coding | 0.882796 | -5.71641 | 2.790777 | -2.04832 | 0.040528 |  |
| 11 | ENSMUSG00000026866 | Kynu | protein_coding | 0.869355 | -5.69577 | 2.666951 | -2.13569 | 0.032705 |  |
| 12 | ENSMUSG00000090509 | Sfta2 | protein_coding | 0.856915 | -5.67834 | 2.828354 | -2.00765 | 0.044681 |  |
| 13 | ENSMUSG00000004872 | Pax3 | protein_coding | 0.853228 | -5.67237 | 2.780985 | -2.0397 | 0.04138 |  |
| 14 | ENSMUSG00000109713 | Pvrig | protein_coding | 2.221633 | -4.50633 | 1.82571 | -2.46826 | 0.013577 |  |
| 15 | ENSMUSG00000068009 | Bpifb6 | protein_coding | 1.676973 | -4.09509 | 2.00745 | -2.03995 | 0.041356 |  |
| 16 | ENSMUSG00000068587 | Mgam | protein_coding | 1.475238 | -4.02377 | 2.006042 | -2.00582 | 0.044875 |  |
| 17 | ENSMUSG00000015962 | 1700016C15Rik | protein_coding | 2.680679 | -3.93657 | 1.843541 | -2.13533 | 0.032734 |  |
| 18 | ENSMUSG00000069044 | Usp9y | protein_coding | 2.938871 | -3.93601 | 1.698104 | -2.31789 | 0.020456 |  |
| 19 | ENSMUSG00000032454 | Rbp2 | protein_coding | 2.693456 | -3.81863 | 1.559564 | -2.44852 | 0.014344 |  |
| 20 | ENSMUSG00000061815 | Rufy4 | protein_coding | 3.835776 | -3.78149 | 1.309192 | -2.88842 | 0.003872 |  |
| 21 | ENSMUSG00000043230 | Fam124b | protein_coding | 2.553544 | -3.62921 | 1.710871 | -2.12127 | 0.033899 |  |
| 22 | ENSMUSG00000042096 | Dao | protein_coding | 2.609433 | -3.15198 | 1.333796 | -2.36316 | 0.01812 |  |
| 23 | ENSMUSG00000095276 | Gfy | protein_coding | 3.934089 | -2.91738 | 1.213859 | -2.40339 | 0.016244 |  |
| 24 | ENSMUSG00000073608 | Gal3st2c | protein_coding | 2.901712 | -2.89571 | 1.381543 | -2.096 | 0.036082 |  |
| 25 | ENSMUSG00000017309 | Cd300lg | protein_coding | 4.057469 | -2.86894 | 1.057878 | -2.71198 | 0.006688 |  |
| 26 | ENSMUSG00000063522 | 2010109I03Rik | protein_coding | 2.258792 | -2.84991 | 1.446748 | -1.96987 | 0.048853 |  |
| 27 | ENSMUSG00000012187 | Mogat1 | protein_coding | 3.025543 | -2.82813 | 1.189841 | -2.37689 | 0.017459 |  |
| 28 | ENSMUSG00000044429 | Cryga | protein_coding | 2.328156 | -2.78923 | 1.363146 | -2.04617 | 0.04074 |  |
| 29 | ENSMUSG00000073574 | Grxcr2 | protein_coding | 3.803221 | -2.60395 | 1.207718 | -2.15609 | 0.031076 |  |
| 30 | ENSMUSG00000090451 | Gm6133 | protein_coding | 6.3777 | -2.53695 | 0.846786 | -2.99597 | 0.002736 |  |
| 31 | ENSMUSG00000027833 | Shox2 | protein_coding | 5.746748 | -2.50569 | 1.097326 | -2.28345 | 0.022404 |  |
| 32 | ENSMUSG00000023279 | Bmp15 | protein_coding | 3.129889 | -2.45155 | 1.152024 | -2.12804 | 0.033334 |  |
| 33 | ENSMUSG00000032484 | Ngp | protein_coding | 5.043166 | -2.43452 | 1.043104 | -2.33391 | 0.0196 |  |
| 34 | ENSMUSG00000043549 | Fam90a1b | protein_coding | 5.792743 | -2.27311 | 0.891126 | -2.55082 | 0.010747 |  |
| 35 | ENSMUSG00000065987 | Cd209b | protein_coding | 5.107185 | -2.27048 | 1.0806 | -2.10113 | 0.035629 |  |
| 36 | ENSMUSG00000030116 | Mfap5 | protein_coding | 6.564673 | -2.06453 | 0.953282 | -2.16571 | 0.030333 |  |
| 37 | ENSMUSG00000024114 | Prss41 | protein_coding | 5.256822 | -1.93378 | 0.886211 | -2.18208 | 0.029104 |  |
| 38 | ENSMUSG00000032769 | Trpa1 | protein_coding | 20.15728 | -1.92683 | 0.651641 | -2.95689 | 0.003108 |  |
| 39 | ENSMUSG00000018752 | Tnfsfm13 | protein_coding | 6.412581 | -1.80062 | 0.771723 | -2.33325 | 0.019635 |  |
| 40 | ENSMUSG00000048572 | Tmem252 | protein_coding | 17.20183 | -1.77673 | 0.545642 | -3.25621 | 0.001129 |  |
| 41 | ENSMUSG00000103800 | Pcdha8 | protein_coding | 9.000795 | -1.76832 | 0.723435 | -2.44434 | 0.014512 |  |
| 42 | ENSMUSG00000049537 | Tecrl | protein_coding | 5.880763 | -1.74714 | 0.813017 | -2.14896 | 0.031638 |  |
| 43 | ENSMUSG00000051906 | Cd209f | protein_coding | 9.316893 | -1.70964 | 0.604748 | -2.82703 | 0.004698 |  |
| 44 | ENSMUSG00000037548 | H2-DMb2 | protein_coding | 6.709271 | -1.70205 | 0.804399 | -2.11593 | 0.034351 |  |
| 45 | ENSMUSG00000030650 | Tmc5 | protein_coding | 6.630004 | -1.69455 | 0.830987 | -2.0392 | 0.041431 |  |
| 46 | ENSMUSG00000067144 | Slc22a7 | protein_coding | 7.40213 | -1.62464 | 0.798338 | -2.03502 | 0.041848 |  |
| 47 | ENSMUSG00000070423 | Olfr558 | protein_coding | 11.15907 | -1.54581 | 0.58749 | -2.63122 | 0.008508 |  |
| 48 | ENSMUSG00000056987 | Fam71d | protein_coding | 6.738904 | -1.53764 | 0.74419 | -2.06619 | 0.03881 |  |
| 49 | ENSMUSG00000026628 | Atf3 | protein_coding | 21.73359 | -1.50543 | 0.385134 | -3.90885 | 9.27E-05 |  |
| 50 | ENSMUSG00000005131 | 4930550C14Rik | protein_coding | 11.8608 | -1.45205 | 0.604699 | -2.40128 | 0.016338 |  |
| 51 | ENSMUSG00000032643 | Fhl3 | protein_coding | 15.41592 | -1.36543 | 0.515144 | -2.65058 | 0.008035 |  |
| 52 | ENSMUSG00000040380 | Cbln3 | protein_coding | 19.95575 | -1.36308 | 0.59849 | -2.27754 | 0.022754 |  |
| 53 | ENSMUSG00000025804 | Ccr1 | protein_coding | 9.328481 | -1.32145 | 0.671091 | -1.96911 | 0.048941 |  |
| 54 | ENSMUSG00000037868 | Egr2 | protein_coding | 76.35897 | -1.23546 | 0.351867 | -3.51116 | 0.000446 |  |
| 55 | ENSMUSG00000018698 | Lhx1 | protein_coding | 11.37602 | -1.21778 | 0.606715 | -2.00717 | 0.044732 |  |
| 56 | ENSMUSG00000032380 | Dapk2 | protein_coding | 13.22957 | -1.19841 | 0.539664 | -2.22067 | 0.026373 |  |
| 57 | ENSMUSG00000060678 | Hist1h4c | protein_coding | 8.298137 | -1.1934 | 0.601572 | -1.9838 | 0.047278 |  |
| 58 | ENSMUSG00000042622 | Maff | protein_coding | 22.53504 | -1.17017 | 0.439714 | -2.66122 | 0.007786 |  |
| 59 | ENSMUSG00000089840 | Gm4491 | protein_coding | 20.33369 | -1.15046 | 0.438823 | -2.62168 | 0.00875 |  |
| 60 | ENSMUSG00000056313 | Tcim | protein_coding | 32.39569 | -1.0675 | 0.341752 | -3.12362 | 0.001786 |  |
| 61 | ENSMUSG00000023484 | Prph | protein_coding | 31.22484 | -1.01852 | 0.349421 | -2.91486 | 0.003558 |  |
| 62 | ENSMUSG00000032611 | 1700102P08Rik | protein_coding | 27.00847 | -1.01631 | 0.346078 | -2.93665 | 0.003318 |  |
| 63 | ENSMUSG00000042345 | Ubash3a | protein_coding | 24.4856 | 1.004466 | 0.469276 | 2.140459 | 0.032318 |  |
| 64 | ENSMUSG00000045102 | Poln | protein_coding | 16.71356 | 1.00946 | 0.492803 | 2.048403 | 0.040521 |  |
| 65 | ENSMUSG00000114278 | AC157931.1 | protein_coding | 26.54265 | 1.03284 | 0.365933 | 2.822488 | 0.004765 |  |
| 66 | ENSMUSG00000028076 | Cd1d1 | protein_coding | 25.98717 | 1.045038 | 0.374419 | 2.791094 | 0.005253 |  |
| 67 | ENSMUSG00000089798 | 1700028K03Rik | protein_coding | 18.45212 | 1.047826 | 0.525795 | 1.992839 | 0.046279 |  |
| 68 | ENSMUSG00000081984 | Dnajb3 | protein_coding | 22.88177 | 1.049692 | 0.437424 | 2.399714 | 0.016408 |  |
| 69 | ENSMUSG00000029847 | Slc23a4 | protein_coding | 15.18705 | 1.078541 | 0.508238 | 2.122117 | 0.033828 |  |
| 70 | ENSMUSG00000071104 | Ccdc110 | protein_coding | 17.40799 | 1.089725 | 0.458401 | 2.377231 | 0.017443 |  |
| 71 | ENSMUSG00000063383 | Zfp947 | protein_coding | 27.93466 | 1.126092 | 0.431 | 2.612741 | 0.008982 |  |
| 72 | ENSMUSG00000037705 | Tecta | protein_coding | 25.51321 | 1.133805 | 0.4602 | 2.463724 | 0.01375 |  |
| 73 | ENSMUSG00000079364 | Gm3558 | protein_coding | 14.74141 | 1.141563 | 0.467759 | 2.440493 | 0.014667 |  |
| 74 | ENSMUSG00000020447 | Npc1l1 | protein_coding | 18.28214 | 1.150246 | 0.446668 | 2.57517 | 0.010019 |  |
| 75 | ENSMUSG00000026147 | Col9a1 | protein_coding | 16.32611 | 1.151015 | 0.473269 | 2.432053 | 0.015014 |  |
| 76 | ENSMUSG00000025592 | Dach2 | protein_coding | 29.21031 | 1.240167 | 0.391379 | 3.168715 | 0.001531 |  |
| 77 | ENSMUSG00000074570 | Cass4 | protein_coding | 21.55141 | 1.242089 | 0.452242 | 2.746511 | 0.006023 |  |
| 78 | ENSMUSG00000063234 | Gpr84 | protein_coding | 25.6405 | 1.243675 | 0.393843 | 3.157791 | 0.00159 |  |
| 79 | ENSMUSG00000046610 | Oacyl | protein_coding | 50.29394 | 1.259185 | 0.406128 | 3.100467 | 0.001932 |  |
| 80 | ENSMUSG00000033114 | Slc35d2 | protein_coding | 18.78147 | 1.289336 | 0.477683 | 2.699146 | 0.006952 |  |
| 81 | ENSMUSG00000062157 | Ifnlr1 | protein_coding | 9.479801 | 1.297579 | 0.637654 | 2.034926 | 0.041858 |  |
| 82 | ENSMUSG00000017817 | Jph2 | protein_coding | 20.29503 | 1.305141 | 0.516217 | 2.528277 | 0.011462 |  |
| 83 | ENSMUSG00000038522 | Mfsd4b1 | protein_coding | 12.20652 | 1.324117 | 0.591448 | 2.238772 | 0.025171 |  |
| 84 | ENSMUSG00000048636 | A730049H05Rik | protein_coding | 9.323039 | 1.340634 | 0.67919 | 1.973873 | 0.048396 |  |
| 85 | ENSMUSG00000009670 | Tex11 | protein_coding | 11.83918 | 1.376044 | 0.584419 | 2.354553 | 0.018545 |  |
| 86 | ENSMUSG00000055976 | Cldn23 | protein_coding | 17.45491 | 1.392417 | 0.523099 | 2.66186 | 0.007771 |  |
| 87 | ENSMUSG00000078157 | 4931440F15Rik | protein_coding | 11.35562 | 1.410473 | 0.610529 | 2.31025 | 0.020874 |  |
| 88 | ENSMUSG00000027318 | Adam33 | protein_coding | 31.30345 | 1.432699 | 0.407966 | 3.511808 | 0.000445 |  |
| 89 | ENSMUSG00000034881 | Tbxa2r | protein_coding | 25.19728 | 1.442228 | 0.391534 | 3.683533 | 0.00023 |  |
| 90 | ENSMUSG00000025519 | Tktl2 | protein_coding | 7.738984 | 1.471286 | 0.68352 | 2.152515 | 0.031357 |  |
| 91 | ENSMUSG00000032068 | Plet1 | protein_coding | 7.403924 | 1.497893 | 0.725153 | 2.065623 | 0.038864 |  |
| 92 | ENSMUSG00000109392 | Gm5737 | protein_coding | 6.999508 | 1.524973 | 0.718425 | 2.122661 | 0.033782 |  |
| 93 | ENSMUSG00000039748 | Exo1 | protein_coding | 12.22501 | 1.546338 | 0.654708 | 2.361873 | 0.018183 |  |
| 94 | ENSMUSG00000031312 | Itgb1bp2 | protein_coding | 8.790817 | 1.606813 | 0.683535 | 2.350741 | 0.018736 |  |
| 95 | ENSMUSG00000033182 | Kbtbd12 | protein_coding | 9.3989 | 1.616824 | 0.790253 | 2.045957 | 0.040761 |  |
| 96 | ENSMUSG00000022033 | Pbk | protein_coding | 10.64605 | 1.708712 | 0.803983 | 2.125307 | 0.033561 |  |
| 97 | ENSMUSG00000032530 | Lyzl4 | protein_coding | 12.01653 | 1.708766 | 0.627381 | 2.72365 | 0.006456 |  |
| 98 | ENSMUSG00000038903 | Ccdc68 | protein_coding | 8.425794 | 1.709894 | 0.842681 | 2.029111 | 0.042447 |  |
| 99 | ENSMUSG00000022034 | Esco2 | protein_coding | 5.97726 | 1.713518 | 0.869402 | 1.970915 | 0.048734 |  |
| 100 | ENSMUSG00000021499 | Catsper3 | protein_coding | 7.039149 | 1.714449 | 0.715615 | 2.39577 | 0.016586 |  |
| 101 | ENSMUSG00000020295 | Hbq1a | protein_coding | 9.441494 | 1.721883 | 0.684609 | 2.515134 | 0.011899 |  |
| 102 | ENSMUSG00000050201 | Otop2 | protein_coding | 9.560318 | 1.734174 | 0.755241 | 2.296187 | 0.021665 |  |
| 103 | ENSMUSG00000114456 | Hist1h2bh | protein_coding | 7.589954 | 1.735536 | 0.720177 | 2.409874 | 0.015958 |  |
| 104 | ENSMUSG00000069917 | Hba-a2 | protein_coding | 13.79281 | 1.737283 | 0.774738 | 2.242413 | 0.024935 |  |
| 105 | ENSMUSG00000005994 | Tyrp1 | protein_coding | 9.18154 | 1.806632 | 0.679238 | 2.659794 | 0.007819 |  |
| 106 | ENSMUSG00000039653 | Baat | protein_coding | 5.780468 | 1.81054 | 0.780075 | 2.320982 | 0.020288 |  |
| 107 | ENSMUSG00000018570 | 2810408A11Rik | protein_coding | 6.10633 | 1.832436 | 0.853084 | 2.148013 | 0.031713 |  |
| 108 | ENSMUSG00000075023 | Accsl | protein_coding | 10.34869 | 1.846999 | 0.585461 | 3.154777 | 0.001606 |  |
| 109 | ENSMUSG00000024041 | Cryaa | protein_coding | 5.262618 | 1.867676 | 0.904821 | 2.064138 | 0.039005 |  |
| 110 | ENSMUSG00000000320 | Alox12 | protein_coding | 9.703697 | 1.881916 | 0.703916 | 2.673494 | 0.007507 |  |
| 111 | ENSMUSG00000045502 | Hcar2 | protein_coding | 4.756048 | 1.968799 | 0.943697 | 2.086261 | 0.036955 |  |
| 112 | ENSMUSG00000048473 | Sult6b2 | protein_coding | 6.773114 | 2.058117 | 0.801181 | 2.568852 | 0.010204 |  |
| 113 | ENSMUSG00000057606 | Colq | protein_coding | 7.929338 | 2.0673 | 0.774583 | 2.668918 | 0.00761 |  |
| 114 | ENSMUSG00000049396 | Gemin4 | protein_coding | 5.034609 | 2.134046 | 0.929909 | 2.294898 | 0.021739 |  |
| 115 | ENSMUSG00000100916 | Lhb | protein_coding | 21.73492 | 2.149001 | 0.992407 | 2.165444 | 0.030354 |  |
| 116 | ENSMUSG00000022245 | Skor1 | protein_coding | 4.526705 | 2.27845 | 1.00275 | 2.272201 | 0.023074 |  |
| 117 | ENSMUSG00000044338 | Aplnr | protein_coding | 5.268808 | 2.3077 | 0.918057 | 2.513679 | 0.011948 |  |
| 118 | ENSMUSG00000043753 | Dmrta1 | protein_coding | 4.800863 | 2.378201 | 1.005169 | 2.365972 | 0.017983 |  |
| 119 | ENSMUSG00000023393 | Slc17a9 | protein_coding | 8.164845 | 2.380314 | 0.746051 | 3.190551 | 0.00142 |  |
| 120 | ENSMUSG00000045802 | Hsf3 | protein_coding | 5.756727 | 2.390008 | 0.959221 | 2.491612 | 0.012716 |  |
| 121 | ENSMUSG00000073492 | Gm10521 | protein_coding | 4.561775 | 2.422396 | 1.090276 | 2.221819 | 0.026296 |  |
| 122 | ENSMUSG00000020712 | Tcam1 | protein_coding | 8.79516 | 2.437067 | 0.763519 | 3.191886 | 0.001413 |  |
| 123 | ENSMUSG00000038567 | Cyp24a1 | protein_coding | 5.189056 | 2.474073 | 0.963677 | 2.567326 | 0.010249 |  |
| 124 | ENSMUSG00000025270 | Alas2 | protein_coding | 15.78231 | 2.524726 | 0.849785 | 2.971016 | 0.002968 |  |
| 125 | ENSMUSG00000022584 | Ly6c2 | protein_coding | 2.732496 | 2.593293 | 1.295676 | 2.001498 | 0.045339 |  |
| 126 | ENSMUSG00000070504 | Fcrl6 | protein_coding | 2.98663 | 2.742195 | 1.297713 | 2.113098 | 0.034592 |  |
| 127 | ENSMUSG00000020660 | Pomc | protein_coding | 747.9298 | 2.937634 | 1.335219 | 2.200114 | 0.027799 | 0.33535983 |
| 128 | ENSMUSG00000024124 | Prss30 | protein_coding | 3.399763 | 2.964991 | 1.277294 | 2.321307 | 0.02027 |  |
| 129 | ENSMUSG00000022491 | Glycam1 | protein_coding | 6.808508 | 3.009652 | 0.964907 | 3.119111 | 0.001814 |  |
| 130 | ENSMUSG00000005836 | Gata6 | protein_coding | 2.617049 | 3.127423 | 1.538497 | 2.032778 | 0.042075 |  |
| 131 | ENSMUSG00000026645 | Olah | protein_coding | 2.604531 | 3.136553 | 1.535247 | 2.043028 | 0.04105 |  |
| 132 | ENSMUSG00000076438 | Oxct2b | protein_coding | 3.97815 | 3.194622 | 1.157001 | 2.761122 | 0.00576 |  |
| 133 | ENSMUSG00000026011 | Ctla4 | protein_coding | 2.778782 | 3.21263 | 1.540505 | 2.085439 | 0.037029 |  |
| 134 | ENSMUSG00000100586 | Vmn1r90 | protein_coding | 3.033582 | 3.423661 | 1.502295 | 2.278953 | 0.02267 |  |
| 135 | ENSMUSG00000006546 | Cryba2 | protein_coding | 3.219729 | 3.518262 | 1.521281 | 2.312697 | 0.020739 |  |
| 136 | ENSMUSG00000030838 | Ush1c | protein_coding | 3.609397 | 3.621316 | 1.428813 | 2.534493 | 0.011261 |  |
| 137 | ENSMUSG00000050087 | Cby3 | protein_coding | 1.858755 | 3.715909 | 1.868165 | 1.989069 | 0.046694 |  |
| 138 | ENSMUSG00000023120 | Gm853 | protein_coding | 3.53766 | 3.740135 | 1.534728 | 2.437002 | 0.01481 |  |
| 139 | ENSMUSG00000058398 | Prss43 | protein_coding | 1.980002 | 3.886448 | 1.953431 | 1.989549 | 0.046641 |  |
| 140 | ENSMUSG00000001510 | Dlx3 | protein_coding | 2.130355 | 3.999516 | 2.029971 | 1.970233 | 0.048812 |  |
| 141 | ENSMUSG00000008601 | Rab25 | protein_coding | 2.177965 | 4.027282 | 2.00027 | 2.013369 | 0.044076 |  |
| 142 | ENSMUSG00000050424 | Pnma5 | protein_coding | 2.366991 | 4.048012 | 1.966059 | 2.058947 | 0.039499 |  |
| 143 | ENSMUSG00000028587 | Orc1 | protein_coding | 3.749604 | 4.830023 | 1.798889 | 2.685004 | 0.007253 |  |
| 144 | ENSMUSG00000044854 | 1700056E22Rik | protein_coding | 1.057508 | 5.742543 | 2.929363 | 1.960338 | 0.049956 |  |
| 145 | ENSMUSG00000022229 | Atp12a | protein_coding | 1.06341 | 5.771148 | 2.808374 | 2.054978 | 0.039881 |  |
| 146 | ENSMUSG00000031448 | Adprhl1 | protein_coding | 1.102384 | 5.791514 | 2.796709 | 2.070832 | 0.038375 |  |
| 147 | ENSMUSG00000004651 | Tyr | protein_coding | 1.110011 | 5.823398 | 2.895958 | 2.010871 | 0.044339 |  |
| 148 | ENSMUSG00000047592 | Nxpe5 | protein_coding | 1.121649 | 5.85346 | 2.895937 | 2.021266 | 0.043252 |  |
| 149 | ENSMUSG00000016758 | Bik | protein_coding | 1.150929 | 5.868474 | 2.772623 | 2.116579 | 0.034296 |  |
| 150 | ENSMUSG00000073967 | Olfr557 | protein_coding | 1.349875 | 6.088042 | 2.845399 | 2.139609 | 0.032386 |  |
| 151 | ENSMUSG00000041754 | Trem3 | protein_coding | 1.387162 | 6.138885 | 2.831586 | 2.168002 | 0.030159 |  |
| 152 | ENSMUSG00000024675 | Ms4a4c | protein_coding | 1.377559 | 6.142082 | 2.805137 | 2.189584 | 0.028554 |  |
| 153 | ENSMUSG00000037390 | Muc3 | protein_coding | 1.475533 | 6.216569 | 2.820062 | 2.204409 | 0.027496 |  |
| 154 | ENSMUSG00000029814 | Igf2bp3 | protein_coding | 1.477257 | 6.255519 | 2.79262 | 2.240018 | 0.02509 |  |
| 155 | ENSMUSG00000039492 | Ccdc27 | protein_coding | 1.499247 | 6.268492 | 2.919147 | 2.147371 | 0.031764 |  |
| 156 | ENSMUSG00000075510 | Fam187a | protein_coding | 1.524745 | 6.271164 | 2.736989 | 2.291264 | 0.021948 |  |
| 157 | ENSMUSG00000048498 | Cd300e | protein_coding | 1.58875 | 6.334312 | 2.754356 | 2.299743 | 0.021463 |  |
| 158 | ENSMUSG00000052471 | Gm9881 | protein_coding | 1.610501 | 6.366766 | 2.665925 | 2.388201 | 0.016931 |  |
| 159 | ENSMUSG00000029369 | Afm | protein_coding | 1.667293 | 6.394763 | 2.632023 | 2.4296 | 0.015116 |  |
| 160 | ENSMUSG00000042861 | Kcna10 | protein_coding | 1.65878 | 6.402145 | 2.626943 | 2.437109 | 0.014805 |  |
| 161 | ENSMUSG00000028314 | Toporsl | protein_coding | 1.673291 | 6.428548 | 2.883951 | 2.229077 | 0.025809 |  |
| 162 | ENSMUSG00000026327 | Serpinb11 | protein_coding | 1.707286 | 6.441293 | 2.883668 | 2.233716 | 0.025502 |  |
| 163 | ENSMUSG00000021506 | Pitx1 | protein_coding | 1.891556 | 6.591597 | 2.899457 | 2.27339 | 0.023003 |  |
| 164 | ENSMUSG00000037161 | Mgarp | protein_coding | 2.105841 | 6.756654 | 2.60991 | 2.588845 | 0.00963 |  |
| 165 | ENSMUSG00000004814 | Ccl24 | protein_coding | 2.206804 | 6.82486 | 2.600953 | 2.623984 | 0.008691 |  |

**Supplementary Table 4: Naïve vs CCI + PBS Nex-Cre:TNFR2^F/F^**

| **S.No** | **Gene_ID** | **Gene_Name** | **Gene_Biotype** | **baseMean** | **log2FoldChange** | **lfcSE** | **stat** | **pvalue** | **padj** |
| --- | --- | --- | --- | --- | --- | --- | --- | --- | --- |
| 1 | ENSMUSG00000047631 | Apof | protein_coding | 1.972493 | -6.75606 | 2.64524 | -2.55404 | 0.010648 | 0.956682702 |
| 2 | ENSMUSG00000045493 | Bhlhe23 | protein_coding | 1.905339 | -6.68332 | 2.876783 | -2.32319 | 0.020169 | 0.956682702 |
| 3 | ENSMUSG00000073551 | Spink13 | protein_coding | 1.883259 | -6.673 | 2.766607 | -2.41198 | 0.015866 | 0.956682702 |
| 4 | ENSMUSG00000056257 | Gm5447 | protein_coding | 1.817636 | -6.61971 | 2.890087 | -2.29049 | 0.021993 | 0.956682702 |
| 5 | ENSMUSG00000073574 | Grxcr2 | protein_coding | 1.711569 | -6.5517 | 2.662121 | -2.46108 | 0.013852 | 0.956682702 |
| 6 | ENSMUSG00000075370 | Igll1 | protein_coding | 1.485264 | -6.32565 | 2.750029 | -2.30021 | 0.021436 | 0.956682702 |
| 7 | ENSMUSG00000047592 | Nxpe5 | protein_coding | 1.434316 | -6.31356 | 2.889883 | -2.18471 | 0.02891 | 0.956682702 |
| 8 | ENSMUSG00000044988 | Ucn3 | protein_coding | 1.433038 | -6.30625 | 3.179523 | -1.98339 | 0.047323 | 0.956682702 |
| 9 | ENSMUSG00000022039 | Adam2 | protein_coding | 1.358736 | -6.20217 | 3.001693 | -2.06622 | 0.038807 | 0.956682702 |
| 10 | ENSMUSG00000021953 | Tdh | protein_coding | 1.255304 | -6.11355 | 3.077945 | -1.98624 | 0.047006 | 0.956682702 |
| 11 | ENSMUSG00000052187 | Hbb-y | protein_coding | 1.210694 | -6.06711 | 3.093669 | -1.96114 | 0.049863 | 0.956682702 |
| 12 | ENSMUSG00000031518 | Spata4 | protein_coding | 1.222269 | -6.06599 | 2.787341 | -2.17627 | 0.029535 | 0.956682702 |
| 13 | ENSMUSG00000032315 | Cyp1a1 | protein_coding | 2.075979 | -3.95424 | 1.909257 | -2.07109 | 0.038351 | 0.956682702 |
| 14 | ENSMUSG00000090872 | Gm3239 | protein_coding | 3.343467 | -3.67233 | 1.453304 | -2.52688 | 0.011508 | 0.956682702 |
| 15 | ENSMUSG00000044378 | Slc15a5 | protein_coding | 2.876899 | -3.45578 | 1.722508 | -2.00625 | 0.04483 | 0.956682702 |
| 16 | ENSMUSG00000064010 | Dppa1 | protein_coding | 2.527201 | -3.17615 | 1.536039 | -2.06775 | 0.038663 | 0.956682702 |
| 17 | ENSMUSG00000072952 | Gm5878 | protein_coding | 3.310066 | -3.01325 | 1.522846 | -1.9787 | 0.04785 | 0.956682702 |
| 18 | ENSMUSG00000079497 | Gm13420 | protein_coding | 2.963432 | -2.78533 | 1.293731 | -2.15294 | 0.031323 | 0.956682702 |
| 19 | ENSMUSG00000075307 | Klhl41 | protein_coding | 4.358611 | -2.26721 | 1.058837 | -2.14123 | 0.032255 | 0.956682702 |
| 20 | ENSMUSG00000020808 | Pimreg | protein_coding | 7.572554 | -2.17246 | 0.920229 | -2.36078 | 0.018237 | 0.956682702 |
| 21 | ENSMUSG00000009097 | Tbx1 | protein_coding | 22.23933 | -1.67026 | 0.522988 | -3.19368 | 0.001405 | 0.790334294 |
| 22 | ENSMUSG00000004668 | Abca13 | protein_coding | 9.175243 | -1.66106 | 0.676888 | -2.45397 | 0.014129 | 0.956682702 |
| 23 | ENSMUSG00000073964 | Olfr570 | protein_coding | 15.56433 | -1.49006 | 0.731017 | -2.03834 | 0.041516 | 0.956682702 |
| 24 | ENSMUSG00000005628 | Tmod4 | protein_coding | 11.38596 | -1.42657 | 0.633856 | -2.25063 | 0.024409 | 0.956682702 |
| 25 | ENSMUSG00000054667 | Irs4 | protein_coding | 38.37725 | -1.35839 | 0.49018 | -2.77121 | 0.005585 | 0.956682702 |
| 26 | ENSMUSG00000024972 | Lgals12 | protein_coding | 13.164 | -1.26751 | 0.621873 | -2.03821 | 0.041529 | 0.956682702 |
| 27 | ENSMUSG00000027377 | Mall | protein_coding | 17.92286 | -1.18331 | 0.555291 | -2.13096 | 0.033092 | 0.956682702 |
| 28 | ENSMUSG00000036091 | Hyal3 | protein_coding | 19.42791 | -1.15423 | 0.550725 | -2.09584 | 0.036097 | 0.956682702 |
| 29 | ENSMUSG00000071665 | Foxr2 | protein_coding | 21.61756 | -1.1035 | 0.459845 | -2.39973 | 0.016407 | 0.956682702 |
| 30 | ENSMUSG00000036523 | Greb1 | protein_coding | 31.12261 | -1.08908 | 0.464825 | -2.34299 | 0.01913 | 0.956682702 |
| 31 | ENSMUSG00000032311 | Nrg4 | protein_coding | 13.45392 | -1.08174 | 0.525863 | -2.05708 | 0.039679 | 0.956682702 |
| 32 | ENSMUSG00000066090 | Insl5 | protein_coding | 20.07824 | 1.016455 | 0.460433 | 2.207607 | 0.027272 | 0.956682702 |
| 33 | ENSMUSG00000026407 | Cacna1s | protein_coding | 21.50281 | 1.018076 | 0.468632 | 2.172444 | 0.029822 | 0.956682702 |
| 34 | ENSMUSG00000046886 | Zfp474 | protein_coding | 29.82854 | 1.021486 | 0.395041 | 2.585775 | 0.009716 | 0.956682702 |
| 35 | ENSMUSG00000084989 | Crocc2 | protein_coding | 62.04349 | 1.068082 | 0.320218 | 3.335485 | 0.000852 | 0.722917851 |
| 36 | ENSMUSG00000040247 | Tbc1d10c | protein_coding | 24.35413 | 1.069317 | 0.491713 | 2.174676 | 0.029654 | 0.956682702 |
| 37 | ENSMUSG00000036972 | Zic4 | protein_coding | 141.5439 | 1.078327 | 0.489938 | 2.200947 | 0.02774 | 0.956682702 |
| 38 | ENSMUSG00000047497 | Adamts12 | protein_coding | 18.23146 | 1.078479 | 0.528318 | 2.041344 | 0.041217 | 0.956682702 |
| 39 | ENSMUSG00000010476 | Ebf3 | protein_coding | 53.24547 | 1.094976 | 0.340613 | 3.21472 | 0.001306 | 0.781516617 |
| 40 | ENSMUSG00000024164 | C3 | protein_coding | 48.72638 | 1.106962 | 0.376497 | 2.940164 | 0.00328 | 0.956682702 |
| 41 | ENSMUSG00000069919 | Hba-a1 | protein_coding | 40.67289 | 1.11134 | 0.46552 | 2.387307 | 0.016972 | 0.956682702 |
| 42 | ENSMUSG00000054320 | Lrrc36 | protein_coding | 43.04292 | 1.11893 | 0.390215 | 2.867471 | 0.004138 | 0.956682702 |
| 43 | ENSMUSG00000038011 | Dnah10 | protein_coding | 95.51269 | 1.142334 | 0.282772 | 4.039767 | 5.35E-05 | 0.330296969 |
| 44 | ENSMUSG00000047671 | Tctex1d4 | protein_coding | 21.11651 | 1.188143 | 0.496495 | 2.39306 | 0.016709 | 0.956682702 |
| 45 | ENSMUSG00000000889 | Dbh | protein_coding | 18.02011 | 1.21505 | 0.566991 | 2.142979 | 0.032115 | 0.956682702 |
| 46 | ENSMUSG00000017861 | Mybl2 | protein_coding | 24.03111 | 1.231353 | 0.436523 | 2.820821 | 0.00479 | 0.956682702 |
| 47 | ENSMUSG00000022033 | Pbk | protein_coding | 12.52195 | 1.233301 | 0.560722 | 2.199486 | 0.027843 | 0.956682702 |
| 48 | ENSMUSG00000052305 | Hbb-bs | protein_coding | 89.32975 | 1.234477 | 0.343768 | 3.591024 | 0.000329 | 0.620123221 |
| 49 | ENSMUSG00000009876 | Cox4i2 | protein_coding | 14.04627 | 1.297252 | 0.587949 | 2.206402 | 0.027356 | 0.956682702 |
| 50 | ENSMUSG00000047220 | Ccdc36 | protein_coding | 12.64845 | 1.332287 | 0.661898 | 2.012827 | 0.044133 | 0.956682702 |
| 51 | ENSMUSG00000063681 | Crb1 | protein_coding | 22.95332 | 1.355542 | 0.505366 | 2.6823 | 0.007312 | 0.956682702 |
| 52 | ENSMUSG00000104301 | Wdr49 | protein_coding | 16.10146 | 1.370617 | 0.586221 | 2.338054 | 0.019384 | 0.956682702 |
| 53 | ENSMUSG00000063594 | Gng8 | protein_coding | 19.58296 | 1.384815 | 0.483571 | 2.863725 | 0.004187 | 0.956682702 |
| 54 | ENSMUSG00000030549 | Rhcg | protein_coding | 10.90827 | 1.440704 | 0.683879 | 2.106665 | 0.035147 | 0.956682702 |
| 55 | ENSMUSG00000031893 | Tsnaxip1 | protein_coding | 29.25315 | 1.467537 | 0.530595 | 2.76583 | 0.005678 | 0.956682702 |
| 56 | ENSMUSG00000039956 | Mrap | protein_coding | 10.17319 | 1.470426 | 0.692161 | 2.124399 | 0.033637 | 0.956682702 |
| 57 | ENSMUSG00000000791 | Il12rb1 | protein_coding | 18.76647 | 1.480135 | 0.627398 | 2.359166 | 0.018316 | 0.956682702 |
| 58 | ENSMUSG00000079343 | C1s2 | protein_coding | 9.10029 | 1.484151 | 0.689287 | 2.153169 | 0.031305 | 0.956682702 |
| 59 | ENSMUSG00000024747 | Aldh1a7 | protein_coding | 7.718516 | 1.500446 | 0.744864 | 2.014389 | 0.043969 | 0.956682702 |
| 60 | ENSMUSG00000015533 | Itga2 | protein_coding | 8.354369 | 1.538192 | 0.761559 | 2.019794 | 0.043405 | 0.956682702 |
| 61 | ENSMUSG00000029917 | C130060K24Rik | protein_coding | 8.274739 | 1.57998 | 0.774911 | 2.038918 | 0.041458 | 0.956682702 |
| 62 | ENSMUSG00000022945 | Chaf1b | protein_coding | 9.507669 | 1.603364 | 0.680571 | 2.355908 | 0.018477 | 0.956682702 |
| 63 | ENSMUSG00000006270 | Vax1 | protein_coding | 9.981787 | 1.626487 | 0.72649 | 2.238829 | 0.025167 | 0.956682702 |
| 64 | ENSMUSG00000036768 | Kif15 | protein_coding | 13.6805 | 1.730554 | 0.632668 | 2.735328 | 0.006232 | 0.956682702 |
| 65 | ENSMUSG00000060030 | Olfr317 | protein_coding | 7.544364 | 1.805379 | 0.861355 | 2.095976 | 0.036084 | 0.956682702 |
| 66 | ENSMUSG00000045929 | Gm20715 | protein_coding | 6.691267 | 1.817889 | 0.878969 | 2.068205 | 0.038621 | 0.956682702 |
| 67 | ENSMUSG00000039865 | Slc44a3 | protein_coding | 5.220228 | 1.856238 | 0.895963 | 2.07178 | 0.038286 | 0.956682702 |
| 68 | ENSMUSG00000053615 | Gm9913 | protein_coding | 5.238672 | 1.975037 | 0.932256 | 2.118557 | 0.034128 | 0.956682702 |
| 69 | ENSMUSG00000050700 | Emilin3 | protein_coding | 6.323761 | 2.002667 | 1.020596 | 1.962252 | 0.049733 | 0.956682702 |
| 70 | ENSMUSG00000072972 | Adam4 | protein_coding | 8.820581 | 2.006435 | 0.771361 | 2.601161 | 0.009291 | 0.956682702 |
| 71 | ENSMUSG00000028280 | Gabrr1 | protein_coding | 5.275378 | 2.017503 | 0.966173 | 2.088139 | 0.036785 | 0.956682702 |
| 72 | ENSMUSG00000007030 | Vwa7 | protein_coding | 5.340171 | 2.079436 | 0.904234 | 2.299665 | 0.021467 | 0.956682702 |
| 73 | ENSMUSG00000079391 | Gm2974 | protein_coding | 16.02462 | 2.087025 | 0.624982 | 3.339337 | 0.00084 | 0.722917851 |
| 74 | ENSMUSG00000042379 | Esm1 | protein_coding | 6.890642 | 2.090724 | 1.036916 | 2.016292 | 0.043769 | 0.956682702 |
| 75 | ENSMUSG00000023903 | Mmp25 | protein_coding | 8.132487 | 2.093597 | 0.860179 | 2.433908 | 0.014937 | 0.956682702 |
| 76 | ENSMUSG00000026609 | Ush2a | protein_coding | 4.048815 | 2.123406 | 1.075953 | 1.973512 | 0.048437 | 0.956682702 |
| 77 | ENSMUSG00000093865 | Lrit3 | protein_coding | 4.509083 | 2.142063 | 1.020149 | 2.099754 | 0.035751 | 0.956682702 |
| 78 | ENSMUSG00000020660 | Pomc | protein_coding | 323.0661 | 2.17805 | 0.946191 | 2.301914 | 0.02134 | 0.956682702 |
| 79 | ENSMUSG00000047819 | Tigd4 | protein_coding | 4.147112 | 2.223366 | 1.088373 | 2.042835 | 0.041069 | 0.956682702 |
| 80 | ENSMUSG00000045392 | Olfr1033 | protein_coding | 4.293171 | 2.233072 | 1.041251 | 2.144605 | 0.031984 | 0.956682702 |
| 81 | ENSMUSG00000043753 | Dmrta1 | protein_coding | 4.23897 | 2.270953 | 1.080138 | 2.102465 | 0.035513 | 0.956682702 |
| 82 | ENSMUSG00000107108 | Gm9936 | protein_coding | 6.508338 | 2.358714 | 0.921265 | 2.560298 | 0.010458 | 0.956682702 |
| 83 | ENSMUSG00000020911 | Krt19 | protein_coding | 4.31822 | 2.630205 | 1.236149 | 2.127742 | 0.033359 | 0.956682702 |
| 84 | ENSMUSG00000039878 | Slc39a5 | protein_coding | 5.532374 | 2.709886 | 1.002734 | 2.702498 | 0.006882 | 0.956682702 |
| 85 | ENSMUSG00000074991 | Gabrr3 | protein_coding | 2.965448 | 2.898973 | 1.393124 | 2.080915 | 0.037442 | 0.956682702 |
| 86 | ENSMUSG00000050107 | Haspin | protein_coding | 6.006344 | 3.154193 | 0.972112 | 3.24468 | 0.001176 | 0.781516617 |
| 87 | ENSMUSG00000005763 | Cd247 | protein_coding | 2.517683 | 3.283552 | 1.552021 | 2.115662 | 0.034374 | 0.956682702 |
| 88 | ENSMUSG00000042189 | Tekt3 | protein_coding | 4.913927 | 3.565038 | 1.226818 | 2.905921 | 0.003662 | 0.956682702 |
| 89 | ENSMUSG00000037446 | Tulp1 | protein_coding | 3.271661 | 3.577026 | 1.581737 | 2.261454 | 0.023731 | 0.956682702 |
| 90 | ENSMUSG00000021790 | Dydc1 | protein_coding | 3.285849 | 3.630312 | 1.493769 | 2.430304 | 0.015086 | 0.956682702 |
| 91 | ENSMUSG00000029491 | Pde6b | protein_coding | 3.696014 | 3.780302 | 1.393645 | 2.712529 | 0.006677 | 0.956682702 |
| 92 | ENSMUSG00000028072 | Ntrk1 | protein_coding | 1.864891 | 3.875879 | 1.973408 | 1.964053 | 0.049524 | 0.956682702 |
| 93 | ENSMUSG00000078597 | Cyp4a12b | protein_coding | 2.074713 | 4.048981 | 2.04624 | 1.978742 | 0.047845 | 0.956682702 |
| 94 | ENSMUSG00000030523 | Trpm1 | protein_coding | 2.830862 | 4.425715 | 1.831959 | 2.415837 | 0.015699 | 0.956682702 |
| 95 | ENSMUSG00000079466 | Prdm12 | protein_coding | 8.614144 | 5.005167 | 2.440209 | 2.051122 | 0.040255 | 0.956682702 |
| 96 | ENSMUSG00000064252 | Olfr329-ps | protein_coding | 1.093501 | 5.901185 | 2.967358 | 1.9887 | 0.046734 | 0.956682702 |
| 97 | ENSMUSG00000091119 | Ccdc152 | protein_coding | 1.11455 | 5.925262 | 2.94639 | 2.011024 | 0.044323 | 0.956682702 |
| 98 | ENSMUSG00000001155 | Ftcd | protein_coding | 1.118905 | 5.93019 | 2.94812 | 2.011516 | 0.044271 | 0.956682702 |
| 99 | ENSMUSG00000096003 | Gm3500 | protein_coding | 1.182982 | 6.017844 | 3.057114 | 1.968472 | 0.049014 | 0.956682702 |
| 100 | ENSMUSG00000040600 | Eps8l3 | protein_coding | 1.278274 | 6.118639 | 3.045739 | 2.008918 | 0.044546 | 0.956682702 |
| 101 | ENSMUSG00000114755 | Galr3 | protein_coding | 1.36389 | 6.216311 | 2.863657 | 2.170759 | 0.029949 | 0.956682702 |
| 102 | ENSMUSG00000037944 | Ccr7 | protein_coding | 1.460084 | 6.317053 | 2.964987 | 2.13055 | 0.033126 | 0.956682702 |
| 103 | ENSMUSG00000024868 | Dkk1 | protein_coding | 1.523448 | 6.381527 | 3.150633 | 2.025475 | 0.042819 | 0.956682702 |
| 104 | ENSMUSG00000026228 | Htr2b | protein_coding | 1.618298 | 6.462382 | 2.783693 | 2.321514 | 0.020259 | 0.956682702 |
| 105 | ENSMUSG00000049560 | Defb20 | protein_coding | 1.658395 | 6.50446 | 2.905877 | 2.238381 | 0.025196 | 0.956682702 |
| 106 | ENSMUSG00000047129 | 1700113H08Rik | protein_coding | 1.792192 | 6.616415 | 2.752122 | 2.404114 | 0.016212 | 0.956682702 |
| 107 | ENSMUSG00000094651 | Gal3st2 | protein_coding | 2.136656 | 6.861577 | 2.679303 | 2.560956 | 0.010438 | 0.956682702 |
| 108 | ENSMUSG00000028314 | Toporsl | protein_coding | 2.397178 | 7.034168 | 2.759953 | 2.548655 | 0.010814 | 0.956682702 |

**Supplementary Table 5: Naïve vs CCI + TNFR2Ag Nex-Cre:TNFR2^F/F^**

| **S.No** | **Gene_ID** | **Gene_Name** | **Gene_Biotype** | **baseMean** | **log2FoldChange** | **lfcSE** | **stat** | **pvalue** | **padj** |
| --- | --- | --- | --- | --- | --- | --- | --- | --- | --- |
| 1 | ENSMUSG00000030068 | Gm20696 | protein_coding | 1.608169 | -6.66574 | 2.547534 | -2.61655 | 0.008882 |  |
| 2 | ENSMUSG00000033825 | Tpsb2 | protein_coding | 1.487647 | -6.56762 | 2.590622 | -2.53515 | 0.01124 |  |
| 3 | ENSMUSG00000044988 | Ucn3 | protein_coding | 1.330439 | -6.417 | 2.956208 | -2.17069 | 0.029955 |  |
| 4 | ENSMUSG00000024128 | Sbp | protein_coding | 1.226396 | -6.29215 | 2.801769 | -2.24578 | 0.024718 |  |
| 5 | ENSMUSG00000052187 | Hbb-y | protein_coding | 1.123849 | -6.17712 | 2.889656 | -2.13766 | 0.032544 |  |
| 6 | ENSMUSG00000093806 | Asmt | protein_coding | 1.134281 | -6.1521 | 3.058331 | -2.01159 | 0.044263 |  |
| 7 | ENSMUSG00000071322 | Tcp10a | protein_coding | 1.103926 | -6.14177 | 2.858704 | -2.14845 | 0.031678 |  |
| 8 | ENSMUSG00000004654 | Ghrhr | protein_coding | 1.062858 | -6.09096 | 2.920303 | -2.08573 | 0.037003 |  |
| 9 | ENSMUSG00000046846 | Spesp1 | protein_coding | 1.014083 | -6.01649 | 2.907906 | -2.06901 | 0.038545 |  |
| 10 | ENSMUSG00000048626 | Klf17 | protein_coding | 0.97318 | -5.92767 | 2.978298 | -1.99029 | 0.046559 |  |
| 11 | ENSMUSG00000027855 | Sycp1 | protein_coding | 0.934549 | -5.87418 | 2.969952 | -1.97787 | 0.047943 |  |
| 12 | ENSMUSG00000052396 | Mogat2 | protein_coding | 0.888893 | -5.82828 | 2.843841 | -2.04944 | 0.040419 |  |
| 13 | ENSMUSG00000079580 | Tmem217 | protein_coding | 0.870673 | -5.80374 | 2.724747 | -2.13001 | 0.033171 |  |
| 14 | ENSMUSG00000031212 | Pgr15l | protein_coding | 3.259508 | -5.26063 | 1.756257 | -2.99537 | 0.002741 |  |
| 15 | ENSMUSG00000034486 | Gbx2 | protein_coding | 4.227119 | -4.66201 | 1.611087 | -2.89371 | 0.003807 |  |
| 16 | ENSMUSG00000029193 | Cckar | protein_coding | 2.031322 | -4.58734 | 2.054416 | -2.23292 | 0.025554 |  |
| 17 | ENSMUSG00000056257 | Gm5447 | protein_coding | 1.777305 | -4.3688 | 2.222793 | -1.96546 | 0.049361 |  |
| 18 | ENSMUSG00000054764 | Mtnr1a | protein_coding | 1.890904 | -4.20656 | 2.127857 | -1.9769 | 0.048053 |  |
| 19 | ENSMUSG00000024757 | Slc22a19 | protein_coding | 1.834904 | -4.20194 | 2.003232 | -2.09758 | 0.035942 |  |
| 20 | ENSMUSG00000054362 | Lexm | protein_coding | 2.040624 | -3.52103 | 1.615125 | -2.18004 | 0.029255 |  |
| 21 | ENSMUSG00000075408 | 6030408B16Rik | protein_coding | 2.451735 | -3.16854 | 1.38718 | -2.28416 | 0.022362 |  |
| 22 | ENSMUSG00000060044 | Tmem26 | protein_coding | 2.413954 | -3.15686 | 1.427199 | -2.21193 | 0.026971 |  |
| 23 | ENSMUSG00000064010 | Dppa1 | protein_coding | 2.39062 | -3.11598 | 1.439702 | -2.16432 | 0.03044 |  |
| 24 | ENSMUSG00000017309 | Cd300lg | protein_coding | 3.276249 | -3.0889 | 1.220016 | -2.53186 | 0.011346 |  |
| 25 | ENSMUSG00000082079 | Dnmt3c | protein_coding | 2.812607 | -2.92116 | 1.42139 | -2.05515 | 0.039865 |  |
| 26 | ENSMUSG00000020808 | Pimreg | protein_coding | 7.055938 | -2.4289 | 0.967518 | -2.51045 | 0.012058 |  |
| 27 | ENSMUSG00000005892 | Trh | protein_coding | 28.51945 | -2.40968 | 1.042776 | -2.31083 | 0.020842 |  |
| 28 | ENSMUSG00000063412 | Gm10131 | protein_coding | 9.091948 | -2.08185 | 0.712154 | -2.92331 | 0.003463 |  |
| 29 | ENSMUSG00000001020 | S100a4 | protein_coding | 5.755954 | -1.95253 | 0.892211 | -2.18842 | 0.028639 |  |
| 30 | ENSMUSG00000068697 | Myoz1 | protein_coding | 6.732057 | -1.88607 | 0.889768 | -2.11973 | 0.034029 |  |
| 31 | ENSMUSG00000043664 | Tmem221 | protein_coding | 7.805783 | -1.86565 | 0.944583 | -1.97511 | 0.048256 |  |
| 32 | ENSMUSG00000032081 | Apoc3 | protein_coding | 6.389142 | -1.8245 | 0.874678 | -2.08591 | 0.036987 |  |
| 33 | ENSMUSG00000043592 | Unc5cl | protein_coding | 9.904048 | -1.60838 | 0.759155 | -2.11865 | 0.03412 |  |
| 34 | ENSMUSG00000053338 | Tarm1 | protein_coding | 8.148429 | -1.49154 | 0.677157 | -2.20266 | 0.027619 |  |
| 35 | ENSMUSG00000094083 | Gm1604a | protein_coding | 7.742843 | -1.45483 | 0.679318 | -2.14161 | 0.032225 |  |
| 36 | ENSMUSG00000057836 | Xlr3a | protein_coding | 36.87241 | -1.28062 | 0.437603 | -2.92645 | 0.003429 |  |
| 37 | ENSMUSG00000025317 | Car5a | protein_coding | 12.098 | -1.19352 | 0.547331 | -2.18061 | 0.029212 |  |
| 38 | ENSMUSG00000022676 | Snai2 | protein_coding | 16.96025 | -1.08807 | 0.493173 | -2.20627 | 0.027365 |  |
| 39 | ENSMUSG00000103442 | Pcdha1 | protein_coding | 25.21991 | -1.06917 | 0.45801 | -2.33439 | 0.019576 |  |
| 40 | ENSMUSG00000042622 | Maff | protein_coding | 20.73642 | -1.06389 | 0.448229 | -2.37355 | 0.017618 |  |
| 41 | ENSMUSG00000009097 | Tbx1 | protein_coding | 25.40226 | -1.02906 | 0.398037 | -2.58533 | 0.009729 |  |
| 42 | ENSMUSG00000042372 | Dmrt3 | protein_coding | 28.96913 | -1.00443 | 0.504087 | -1.99258 | 0.046308 |  |
| 43 | ENSMUSG00000035551 | Igfbpl1 | protein_coding | 50.73605 | 1.001332 | 0.381535 | 2.624482 | 0.008678 |  |
| 44 | ENSMUSG00000024552 | Slc14a2 | protein_coding | 26.12576 | 1.032438 | 0.427277 | 2.416318 | 0.015678 |  |
| 45 | ENSMUSG00000063681 | Crb1 | protein_coding | 21.36671 | 1.033153 | 0.492942 | 2.095891 | 0.036092 |  |
| 46 | ENSMUSG00000057280 | Musk | protein_coding | 45.04056 | 1.03887 | 0.362565 | 2.865332 | 0.004166 |  |
| 47 | ENSMUSG00000052305 | Hbb-bs | protein_coding | 88.06024 | 1.040657 | 0.347322 | 2.996228 | 0.002733 | 0.100890884 |
| 48 | ENSMUSG00000054901 | Arhgef33 | protein_coding | 40.85191 | 1.057844 | 0.315294 | 3.355102 | 0.000793 |  |
| 49 | ENSMUSG00000063594 | Gng8 | protein_coding | 18.41695 | 1.093087 | 0.456328 | 2.395396 | 0.016602 |  |
| 50 | ENSMUSG00000079553 | Kifc1 | protein_coding | 40.53712 | 1.095693 | 0.404869 | 2.706287 | 0.006804 |  |
| 51 | ENSMUSG00000027318 | Adam33 | protein_coding | 29.58296 | 1.096066 | 0.445869 | 2.458266 | 0.013961 |  |
| 52 | ENSMUSG00000046610 | Oacyl | protein_coding | 49.97109 | 1.103291 | 0.315374 | 3.498363 | 0.000468 |  |
| 53 | ENSMUSG00000048489 | 8430408G22Rik | protein_coding | 36.66169 | 1.114018 | 0.422616 | 2.636005 | 0.008389 |  |
| 54 | ENSMUSG00000030048 | Gkn3 | protein_coding | 58.22285 | 1.126792 | 0.313373 | 3.595688 | 0.000324 |  |
| 55 | ENSMUSG00000079677 | Fdx1l | protein_coding | 19.2162 | 1.137829 | 0.514483 | 2.211598 | 0.026994 |  |
| 56 | ENSMUSG00000078879 | Zfp973 | protein_coding | 31.18592 | 1.144779 | 0.504413 | 2.269525 | 0.023236 |  |
| 57 | ENSMUSG00000010476 | Ebf3 | protein_coding | 59.35076 | 1.148583 | 0.264607 | 4.340716 | 1.42E-05 |  |
| 58 | ENSMUSG00000038379 | Ttk | protein_coding | 16.41565 | 1.167719 | 0.497694 | 2.346258 | 0.018963 |  |
| 59 | ENSMUSG00000104713 | Gbp6 | protein_coding | 22.30219 | 1.185115 | 0.492035 | 2.408596 | 0.016014 |  |
| 60 | ENSMUSG00000049134 | Nrap | protein_coding | 14.4465 | 1.191067 | 0.528466 | 2.253818 | 0.024208 |  |
| 61 | ENSMUSG00000069919 | Hba-a1 | protein_coding | 46.00717 | 1.192225 | 0.450638 | 2.645638 | 0.008154 |  |
| 62 | ENSMUSG00000025069 | Gsto2 | protein_coding | 25.7027 | 1.21676 | 0.555282 | 2.191249 | 0.028434 |  |
| 63 | ENSMUSG00000017007 | Rbpjl | protein_coding | 19.31389 | 1.24517 | 0.477432 | 2.608057 | 0.009106 |  |
| 64 | ENSMUSG00000031893 | Tsnaxip1 | protein_coding | 29.14307 | 1.285217 | 0.395351 | 3.250824 | 0.001151 |  |
| 65 | ENSMUSG00000031995 | St14 | protein_coding | 15.62245 | 1.311909 | 0.545933 | 2.403058 | 0.016259 |  |
| 66 | ENSMUSG00000104301 | Wdr49 | protein_coding | 17.27708 | 1.317601 | 0.51516 | 2.557652 | 0.010538 |  |
| 67 | ENSMUSG00000024164 | C3 | protein_coding | 58.90005 | 1.321084 | 0.359256 | 3.677279 | 0.000236 |  |
| 68 | ENSMUSG00000030228 | Pik3c2g | protein_coding | 12.86152 | 1.346478 | 0.528209 | 2.549137 | 0.010799 |  |
| 69 | ENSMUSG00000037801 | Iqch | protein_coding | 9.387426 | 1.37595 | 0.668981 | 2.056785 | 0.039707 |  |
| 70 | ENSMUSG00000087075 | Lbhd2 | protein_coding | 18.3533 | 1.375995 | 0.660711 | 2.082599 | 0.037288 |  |
| 71 | ENSMUSG00000047220 | Ccdc36 | protein_coding | 14.34166 | 1.392155 | 0.538511 | 2.585195 | 0.009732 |  |
| 72 | ENSMUSG00000048924 | Ccdc125 | protein_coding | 15.18363 | 1.423284 | 0.701253 | 2.029629 | 0.042394 |  |
| 73 | ENSMUSG00000085139 | A730046J19Rik | protein_coding | 10.00028 | 1.44913 | 0.704503 | 2.056954 | 0.039691 |  |
| 74 | ENSMUSG00000026175 | Vil1 | protein_coding | 11.53944 | 1.50014 | 0.711681 | 2.107883 | 0.035041 |  |
| 75 | ENSMUSG00000044951 | Mylk4 | protein_coding | 11.5095 | 1.511049 | 0.65163 | 2.318874 | 0.020402 |  |
| 76 | ENSMUSG00000025270 | Alas2 | protein_coding | 16.12697 | 1.579519 | 0.69947 | 2.258164 | 0.023935 |  |
| 77 | ENSMUSG00000002384 | Bmp8b | protein_coding | 11.54183 | 1.596807 | 0.707733 | 2.256226 | 0.024056 |  |
| 78 | ENSMUSG00000028523 | Tctex1d1 | protein_coding | 11.17737 | 1.597674 | 0.663026 | 2.409671 | 0.015967 |  |
| 79 | ENSMUSG00000027322 | Siglec1 | protein_coding | 15.38052 | 1.600645 | 0.520376 | 3.075941 | 0.002098 |  |
| 80 | ENSMUSG00000018570 | 2810408A11Rik | protein_coding | 9.711316 | 1.604451 | 0.661285 | 2.426263 | 0.015255 |  |
| 81 | ENSMUSG00000079343 | C1s2 | protein_coding | 10.98398 | 1.654157 | 0.678697 | 2.437253 | 0.014799 |  |
| 82 | ENSMUSG00000050195 | Scd4 | protein_coding | 22.30157 | 1.65795 | 0.461731 | 3.590727 | 0.00033 |  |
| 83 | ENSMUSG00000036768 | Kif15 | protein_coding | 15.27488 | 1.759751 | 0.526347 | 3.343331 | 0.000828 |  |
| 84 | ENSMUSG00000060030 | Olfr317 | protein_coding | 8.273543 | 1.785002 | 0.854261 | 2.089527 | 0.03666 |  |
| 85 | ENSMUSG00000009210 | Prr29 | protein_coding | 9.332857 | 1.872628 | 0.712293 | 2.629016 | 0.008563 |  |
| 86 | ENSMUSG00000025044 | Msr1 | protein_coding | 5.807713 | 1.876761 | 0.920087 | 2.039765 | 0.041374 |  |
| 87 | ENSMUSG00000032860 | P2ry2 | protein_coding | 25.01791 | 1.964642 | 0.442409 | 4.440786 | 8.96E-06 |  |
| 88 | ENSMUSG00000050201 | Otop2 | protein_coding | 6.366486 | 1.974604 | 0.843384 | 2.341287 | 0.019217 |  |
| 89 | ENSMUSG00000026770 | Il2ra | protein_coding | 12.25365 | 1.997741 | 0.726987 | 2.747972 | 0.005997 |  |
| 90 | ENSMUSG00000110622 | Gm16486 | protein_coding | 5.489282 | 2.110385 | 1.025806 | 2.057295 | 0.039658 |  |
| 91 | ENSMUSG00000033182 | Kbtbd12 | protein_coding | 12.73093 | 2.130149 | 0.651051 | 3.271862 | 0.001068 |  |
| 92 | ENSMUSG00000079391 | Gm2974 | protein_coding | 19.22681 | 2.215936 | 0.491734 | 4.506373 | 6.59E-06 |  |
| 93 | ENSMUSG00000089773 | Skint1 | protein_coding | 4.001091 | 2.224177 | 1.133181 | 1.962773 | 0.049673 |  |
| 94 | ENSMUSG00000013643 | Lypd8 | protein_coding | 4.230466 | 2.29809 | 1.052793 | 2.182852 | 0.029047 |  |
| 95 | ENSMUSG00000026070 | Il18r1 | protein_coding | 3.943311 | 2.351554 | 1.150185 | 2.044501 | 0.040904 |  |
| 96 | ENSMUSG00000050107 | Haspin | protein_coding | 4.247557 | 2.422969 | 1.133448 | 2.137697 | 0.032541 |  |
| 97 | ENSMUSG00000043873 | Chil5 | protein_coding | 3.640902 | 2.424934 | 1.23706 | 1.96024 | 0.049968 |  |
| 98 | ENSMUSG00000058773 | Hist1h1b | protein_coding | 3.787456 | 2.471739 | 1.139728 | 2.16871 | 0.030105 |  |
| 99 | ENSMUSG00000058398 | Prss43 | protein_coding | 6.029461 | 2.528378 | 0.935989 | 2.701291 | 0.006907 |  |
| 100 | ENSMUSG00000020062 | Slc5a8 | protein_coding | 3.956097 | 2.536002 | 1.209268 | 2.097138 | 0.035981 |  |
| 101 | ENSMUSG00000024600 | Slc27a6 | protein_coding | 6.359816 | 2.564174 | 1.071073 | 2.394024 | 0.016665 |  |
| 102 | ENSMUSG00000079386 | Gm3173 | protein_coding | 4.478698 | 2.694261 | 1.199737 | 2.24571 | 0.024723 |  |
| 103 | ENSMUSG00000062939 | Stat4 | protein_coding | 4.959467 | 2.993045 | 1.151833 | 2.598506 | 0.009363 |  |
| 104 | ENSMUSG00000036136 | Fam110c | protein_coding | 2.602176 | 3.152323 | 1.576226 | 1.999918 | 0.045509 |  |
| 105 | ENSMUSG00000025127 | Gcgr | protein_coding | 2.659787 | 3.172154 | 1.546906 | 2.050644 | 0.040302 |  |
| 106 | ENSMUSG00000031443 | F7 | protein_coding | 3.126603 | 3.180139 | 1.533687 | 2.073525 | 0.038123 |  |
| 107 | ENSMUSG00000030775 | Trat1 | protein_coding | 2.76787 | 3.188505 | 1.552324 | 2.05402 | 0.039974 |  |
| 108 | ENSMUSG00000021359 | Tfap2a | protein_coding | 3.491117 | 3.457782 | 1.436807 | 2.406574 | 0.016103 |  |
| 109 | ENSMUSG00000091685 | Gm17359 | protein_coding | 3.497839 | 3.479462 | 1.505332 | 2.311425 | 0.020809 |  |
| 110 | ENSMUSG00000020051 | Pah | protein_coding | 5.675226 | 3.656269 | 1.103957 | 3.311968 | 0.000926 |  |
| 111 | ENSMUSG00000096546 | Smlr1 | protein_coding | 8.205328 | 3.70065 | 0.942583 | 3.926075 | 8.63E-05 |  |
| 112 | ENSMUSG00000034115 | Scn11a | protein_coding | 2.446827 | 4.055596 | 1.92922 | 2.102194 | 0.035536 |  |
| 113 | ENSMUSG00000009487 | Otog | protein_coding | 2.492784 | 4.128679 | 1.876211 | 2.20054 | 0.027769 |  |
| 114 | ENSMUSG00000079466 | Prdm12 | protein_coding | 8.278098 | 4.784532 | 1.400634 | 3.415976 | 0.000636 |  |
| 115 | ENSMUSG00000038199 | 4931409K22Rik | protein_coding | 1.103179 | 5.707501 | 2.882096 | 1.98033 | 0.047666 |  |
| 116 | ENSMUSG00000036731 | Cysrt1 | protein_coding | 1.159335 | 5.78436 | 2.93345 | 1.971862 | 0.048625 |  |
| 117 | ENSMUSG00000096003 | Gm3500 | protein_coding | 1.222254 | 5.8334 | 2.919568 | 1.998035 | 0.045713 |  |
| 118 | ENSMUSG00000072791 | Abcb5 | protein_coding | 1.226885 | 5.857521 | 2.881552 | 2.032766 | 0.042076 |  |
| 119 | ENSMUSG00000051728 | 4930563D23Rik | protein_coding | 1.256702 | 5.88828 | 2.788455 | 2.111664 | 0.034715 |  |
| 120 | ENSMUSG00000027249 | F2 | protein_coding | 1.261591 | 5.905613 | 2.897919 | 2.03788 | 0.041562 |  |
| 121 | ENSMUSG00000099762 | Gm21149 | protein_coding | 1.300885 | 5.917994 | 2.915649 | 2.029735 | 0.042383 |  |
| 122 | ENSMUSG00000028217 | Cdh17 | protein_coding | 1.345772 | 6.003028 | 2.779823 | 2.1595 | 0.030811 |  |
| 123 | ENSMUSG00000006542 | Prkag3 | protein_coding | 1.382178 | 6.022685 | 2.870854 | 2.097872 | 0.035916 |  |
| 124 | ENSMUSG00000056148 | Rdh9 | protein_coding | 1.389913 | 6.042675 | 2.834766 | 2.131631 | 0.033037 |  |
| 125 | ENSMUSG00000039335 | Spata16 | protein_coding | 1.397648 | 6.057125 | 2.979815 | 2.032719 | 0.042081 |  |
| 126 | ENSMUSG00000000204 | Slfn4 | protein_coding | 1.528882 | 6.163899 | 2.822564 | 2.183794 | 0.028977 |  |
| 127 | ENSMUSG00000004709 | Cd244 | protein_coding | 1.611923 | 6.211859 | 2.956404 | 2.101154 | 0.035627 |  |
| 128 | ENSMUSG00000059430 | Actg2 | protein_coding | 1.694133 | 6.328966 | 2.687744 | 2.354751 | 0.018535 |  |
| 129 | ENSMUSG00000024391 | Apom | protein_coding | 2.437297 | 6.820094 | 2.69067 | 2.534719 | 0.011254 |  |

**Supplementary Table 6: CCI + PBS TNFR2^F/F^ vs CCI + PBS Nex-Cre:TNFR2^F/F^**

| **S.No** | **Gene_ID** | **Gene_Name** | **Gene_Biotype** | **baseMean** | **log2FoldChange** | **lfcSE** | **stat** | **pvalue** | **padj** |
| --- | --- | --- | --- | --- | --- | --- | --- | --- | --- |
| 1 | ENSMUSG00000048349 | Pou4f1 | protein_coding | 2.575543 | -7.13636 | 2.774631 | -2.572 | 0.010111 | 0.975320767 |
| 2 | ENSMUSG00000022435 | Upk3a | protein_coding | 2.373809 | -7.02037 | 2.786346 | -2.51956 | 0.01175 | 0.975320767 |
| 3 | ENSMUSG00000034185 | 6430628N08Rik | protein_coding | 1.938147 | -6.73433 | 2.901593 | -2.32091 | 0.020292 | 0.975320767 |
| 4 | ENSMUSG00000006546 | Cryba2 | protein_coding | 1.933452 | -6.70661 | 2.594362 | -2.58507 | 0.009736 | 0.975320767 |
| 5 | ENSMUSG00000073574 | Grxcr2 | protein_coding | 1.829646 | -6.62673 | 2.868403 | -2.31025 | 0.020874 | 0.975320767 |
| 6 | ENSMUSG00000004948 | Zp3 | protein_coding | 1.472922 | -6.32725 | 2.722497 | -2.32406 | 0.020122 | 0.975320767 |
| 7 | ENSMUSG00000090206 | Tepp | protein_coding | 1.391228 | -6.23985 | 2.720252 | -2.29385 | 0.021799 | 0.975320767 |
| 8 | ENSMUSG00000045065 | 9930022D16Rik | protein_coding | 1.396879 | -6.22752 | 2.820099 | -2.20826 | 0.027226 | 0.975320767 |
| 9 | ENSMUSG00000022596 | Slurp1 | protein_coding | 1.356574 | -6.21709 | 2.971182 | -2.09246 | 0.036397 | 0.975320767 |
| 10 | ENSMUSG00000115768 | DPEP2NB | protein_coding | 1.328624 | -6.15602 | 3.019465 | -2.03878 | 0.041472 | 0.975320767 |
| 11 | ENSMUSG00000079168 | Cd209g | protein_coding | 1.296627 | -6.14563 | 2.997527 | -2.05024 | 0.040341 | 0.975320767 |
| 12 | ENSMUSG00000028008 | Asic5 | protein_coding | 1.213082 | -6.05856 | 3.047465 | -1.98807 | 0.046804 | 0.975320767 |
| 13 | ENSMUSG00000052363 | Zdhhc19 | protein_coding | 1.191916 | -6.01136 | 3.042099 | -1.97606 | 0.048149 | 0.975320767 |
| 14 | ENSMUSG00000078872 | Gm14401 | protein_coding | 1.164764 | -5.96418 | 2.813872 | -2.11956 | 0.034043 | 0.975320767 |
| 15 | ENSMUSG00000079183 | C030005K15Rik | protein_coding | 1.129775 | -5.94527 | 2.91647 | -2.03851 | 0.041499 | 0.975320767 |
| 16 | ENSMUSG00000052673 | Gm9887 | protein_coding | 1.12057 | -5.91799 | 2.928105 | -2.0211 | 0.043269 | 0.975320767 |
| 17 | ENSMUSG00000057465 | Saa2 | protein_coding | 0.983291 | -5.73127 | 2.891 | -1.98245 | 0.047429 | 0.975320767 |
| 18 | ENSMUSG00000074449 | Gm15319 | protein_coding | 3.963702 | -4.9507 | 1.806754 | -2.74011 | 0.006142 | 0.975320767 |
| 19 | ENSMUSG00000029866 | Kel | protein_coding | 2.75646 | -4.39513 | 1.848328 | -2.3779 | 0.017412 | 0.975320767 |
| 20 | ENSMUSG00000027713 | 1810062G17Rik | protein_coding | 2.309024 | -4.1385 | 1.968352 | -2.10252 | 0.035508 | 0.975320767 |
| 21 | ENSMUSG00000031362 | Xlr4c | protein_coding | 2.310775 | -4.13589 | 1.966266 | -2.10342 | 0.035429 | 0.975320767 |
| 22 | ENSMUSG00000031397 | Tktl1 | protein_coding | 2.279931 | -4.12413 | 1.839612 | -2.24185 | 0.024971 | 0.975320767 |
| 23 | ENSMUSG00000047940 | Stpg2 | protein_coding | 2.164961 | -4.0298 | 1.934583 | -2.08303 | 0.037248 | 0.975320767 |
| 24 | ENSMUSG00000090872 | Gm3239 | protein_coding | 3.690342 | -3.82903 | 1.637681 | -2.33808 | 0.019383 | 0.975320767 |
| 25 | ENSMUSG00000091648 | C2cd4d | protein_coding | 3.967252 | -3.27279 | 1.391324 | -2.35228 | 0.018658 | 0.975320767 |
| 26 | ENSMUSG00000006204 | 5430419D17Rik | protein_coding | 3.118554 | -2.90793 | 1.358444 | -2.14063 | 0.032304 | 0.975320767 |
| 27 | ENSMUSG00000044338 | Aplnr | protein_coding | 6.373305 | -2.90621 | 1.049712 | -2.76858 | 0.00563 | 0.975320767 |
| 28 | ENSMUSG00000029811 | Aoc1 | protein_coding | 3.142891 | -2.86396 | 1.403701 | -2.04029 | 0.041321 | 0.975320767 |
| 29 | ENSMUSG00000041255 | Tmco5b | protein_coding | 3.908957 | -2.75358 | 1.202805 | -2.2893 | 0.022062 | 0.975320767 |
| 30 | ENSMUSG00000009654 | Oit3 | protein_coding | 4.3523 | -2.57853 | 1.16515 | -2.21305 | 0.026894 | 0.975320767 |
| 31 | ENSMUSG00000024266 | Adad2 | protein_coding | 3.349326 | -2.49852 | 1.240922 | -2.01344 | 0.044068 | 0.975320767 |
| 32 | ENSMUSG00000025091 | Pnliprp2 | protein_coding | 3.30272 | -2.47695 | 1.233592 | -2.00792 | 0.044652 | 0.975320767 |
| 33 | ENSMUSG00000040432 | Ltb4r2 | protein_coding | 4.924435 | -2.46854 | 1.01242 | -2.43826 | 0.014758 | 0.975320767 |
| 34 | ENSMUSG00000053318 | Slamf8 | protein_coding | 4.956266 | -2.25666 | 1.031313 | -2.18815 | 0.028659 | 0.975320767 |
| 35 | ENSMUSG00000058057 | Mettl7a3 | protein_coding | 6.051745 | -1.94336 | 0.970077 | -2.00331 | 0.045144 | 0.975320767 |
| 36 | ENSMUSG00000022941 | Ripply3 | protein_coding | 5.920653 | -1.90119 | 0.958052 | -1.98444 | 0.047207 | 0.975320767 |
| 37 | ENSMUSG00000021898 | Asb14 | protein_coding | 4.730319 | -1.89243 | 0.922638 | -2.05111 | 0.040257 | 0.975320767 |
| 38 | ENSMUSG00000032717 | Mdfi | protein_coding | 7.367786 | -1.65892 | 0.792664 | -2.09285 | 0.036363 | 0.975320767 |
| 39 | ENSMUSG00000092592 | Gm20449 | protein_coding | 11.00756 | -1.60849 | 0.617292 | -2.60571 | 0.009168 | 0.975320767 |
| 40 | ENSMUSG00000009097 | Tbx1 | protein_coding | 21.04847 | -1.55353 | 0.53974 | -2.87829 | 0.003998 | 0.975320767 |
| 41 | ENSMUSG00000027377 | Mall | protein_coding | 21.8285 | -1.5365 | 0.48165 | -3.19007 | 0.001422 | 0.975320767 |
| 42 | ENSMUSG00000033182 | Kbtbd12 | protein_coding | 8.248964 | -1.52962 | 0.698446 | -2.19003 | 0.028522 | 0.975320767 |
| 43 | ENSMUSG00000020787 | P2rx1 | protein_coding | 16.43576 | -1.3248 | 0.493231 | -2.68597 | 0.007232 | 0.975320767 |
| 44 | ENSMUSG00000070891 | Gm12689 | protein_coding | 12.43195 | -1.31168 | 0.644769 | -2.03435 | 0.041917 | 0.975320767 |
| 45 | ENSMUSG00000021768 | Dusp13 | protein_coding | 14.21808 | -1.21902 | 0.552223 | -2.20748 | 0.027281 | 0.975320767 |
| 46 | ENSMUSG00000022996 | Wnt10b | protein_coding | 25.19326 | -1.19098 | 0.465226 | -2.56 | 0.010467 | 0.975320767 |
| 47 | ENSMUSG00000029727 | Cyp3a13 | protein_coding | 21.18611 | -1.17711 | 0.510703 | -2.30488 | 0.021174 | 0.975320767 |
| 48 | ENSMUSG00000027547 | Sall4 | protein_coding | 16.22412 | -1.15943 | 0.539443 | -2.14931 | 0.03161 | 0.975320767 |
| 49 | ENSMUSG00000041202 | Pla2g2d | protein_coding | 21.921 | -1.09489 | 0.496647 | -2.20456 | 0.027485 | 0.975320767 |
| 50 | ENSMUSG00000052631 | Sh2d6 | protein_coding | 14.7252 | -1.05316 | 0.523807 | -2.0106 | 0.044368 | 0.975320767 |
| 51 | ENSMUSG00000021940 | Ptpn20 | protein_coding | 34.28257 | -1.0207 | 0.415722 | -2.45525 | 0.014079 | 0.975320767 |
| 52 | ENSMUSG00000052305 | Hbb-bs | protein_coding | 95.11079 | 1.020453 | 0.325842 | 3.13174 | 0.001738 | 0.975320767 |
| 53 | ENSMUSG00000091449 | Gm10269 | protein_coding | 14.84163 | 1.038169 | 0.52297 | 1.985142 | 0.047129 | 0.975320767 |
| 54 | ENSMUSG00000069919 | Hba-a1 | protein_coding | 41.74877 | 1.075572 | 0.362834 | 2.96436 | 0.003033 | 0.975320767 |
| 55 | ENSMUSG00000007122 | Casq1 | protein_coding | 15.40454 | 1.08945 | 0.500251 | 2.177804 | 0.029421 | 0.975320767 |
| 56 | ENSMUSG00000063383 | Zfp947 | protein_coding | 26.26216 | 1.109404 | 0.486368 | 2.280997 | 0.022549 | 0.975320767 |
| 57 | ENSMUSG00000019368 | Sec14l4 | protein_coding | 30.32152 | 1.113978 | 0.346881 | 3.211413 | 0.001321 | 0.956990441 |
| 58 | ENSMUSG00000039518 | Cdsn | protein_coding | 18.24333 | 1.116298 | 0.461755 | 2.417512 | 0.015627 | 0.975320767 |
| 59 | ENSMUSG00000021751 | Acox2 | protein_coding | 17.14666 | 1.148451 | 0.555034 | 2.069154 | 0.038532 | 0.975320767 |
| 60 | ENSMUSG00000043333 | Rhbdl2 | protein_coding | 12.95649 | 1.154639 | 0.549753 | 2.100285 | 0.035704 | 0.975320767 |
| 61 | ENSMUSG00000038763 | Alpk3 | protein_coding | 25.02641 | 1.165073 | 0.419661 | 2.776225 | 0.005499 | 0.975320767 |
| 62 | ENSMUSG00000044201 | Cdc25c | protein_coding | 14.02087 | 1.184819 | 0.600849 | 1.971908 | 0.04862 | 0.975320767 |
| 63 | ENSMUSG00000048489 | 8430408G22Rik | protein_coding | 27.48995 | 1.271411 | 0.385858 | 3.295023 | 0.000984 | 0.825224356 |
| 64 | ENSMUSG00000032899 | Styk1 | protein_coding | 15.1734 | 1.28567 | 0.518555 | 2.47933 | 0.013163 | 0.975320767 |
| 65 | ENSMUSG00000068794 | Col28a1 | protein_coding | 13.8771 | 1.410915 | 0.571879 | 2.467159 | 0.013619 | 0.975320767 |
| 66 | ENSMUSG00000037600 | Kdf1 | protein_coding | 7.655039 | 1.456657 | 0.740474 | 1.967197 | 0.04916 | 0.975320767 |
| 67 | ENSMUSG00000019768 | Esr1 | protein_coding | 298.2703 | 1.490948 | 0.129201 | 11.53979 | 8.31320865684084e-31 | 2.50950829724054e-26 |
| 68 | ENSMUSG00000074817 | Papolb | protein_coding | 12.95113 | 1.496785 | 0.711184 | 2.104637 | 0.035323 | 0.975320767 |
| 69 | ENSMUSG00000055271 | 9330161L09Rik | protein_coding | 13.74891 | 1.529334 | 0.646131 | 2.36691 | 0.017937 | 0.975320767 |
| 70 | ENSMUSG00000048572 | Tmem252 | protein_coding | 17.1691 | 1.546616 | 0.546032 | 2.832463 | 0.004619 | 0.975320767 |
| 71 | ENSMUSG00000035165 | Kcne3 | protein_coding | 10.76675 | 1.575764 | 0.604065 | 2.608598 | 0.009091 | 0.975320767 |
| 72 | ENSMUSG00000079355 | Ackr4 | protein_coding | 11.77279 | 1.582034 | 0.714382 | 2.214549 | 0.026791 | 0.975320767 |
| 73 | ENSMUSG00000025064 | Col17a1 | protein_coding | 27.42161 | 1.586962 | 0.451543 | 3.514534 | 0.000441 | 0.56796057 |
| 74 | ENSMUSG00000029032 | Arhgef16 | protein_coding | 12.38901 | 1.638046 | 0.607701 | 2.695482 | 0.007029 | 0.975320767 |
| 75 | ENSMUSG00000087075 | Lbhd2 | protein_coding | 13.19764 | 1.71714 | 0.783958 | 2.190347 | 0.028499 | 0.975320767 |
| 76 | ENSMUSG00000039092 | Sptlc3 | protein_coding | 18.21938 | 1.796213 | 0.614944 | 2.920935 | 0.00349 | 0.975320767 |
| 77 | ENSMUSG00000102416 | 4933424G06Rik | protein_coding | 8.891243 | 1.865965 | 0.919429 | 2.029482 | 0.042409 | 0.975320767 |
| 78 | ENSMUSG00000057191 | AB124611 | protein_coding | 6.723174 | 1.920915 | 0.914704 | 2.100039 | 0.035725 | 0.975320767 |
| 79 | ENSMUSG00000046223 | Plaur | protein_coding | 6.96661 | 1.926516 | 0.819703 | 2.350262 | 0.01876 | 0.975320767 |
| 80 | ENSMUSG00000027833 | Shox2 | protein_coding | 9.991854 | 2.045573 | 0.832485 | 2.45719 | 0.014003 | 0.975320767 |
| 81 | ENSMUSG00000069917 | Hba-a2 | protein_coding | 8.782911 | 2.127954 | 0.827137 | 2.572676 | 0.010092 | 0.975320767 |
| 82 | ENSMUSG00000042189 | Tekt3 | protein_coding | 5.619954 | 2.173396 | 0.934709 | 2.325213 | 0.020061 | 0.975320767 |
| 83 | ENSMUSG00000041534 | Rbp3 | protein_coding | 5.765114 | 2.194542 | 1.069008 | 2.052876 | 0.040085 | 0.975320767 |
| 84 | ENSMUSG00000093865 | Lrit3 | protein_coding | 4.588715 | 2.243134 | 1.132725 | 1.980299 | 0.04767 | 0.975320767 |
| 85 | ENSMUSG00000023943 | Sult1c1 | protein_coding | 11.25703 | 2.322113 | 0.959088 | 2.421168 | 0.015471 | 0.975320767 |
| 86 | ENSMUSG00000107108 | Gm9936 | protein_coding | 6.463038 | 2.390202 | 0.879526 | 2.717602 | 0.006576 | 0.975320767 |
| 87 | ENSMUSG00000020123 | Avpr1a | protein_coding | 7.997871 | 2.42532 | 0.996691 | 2.433372 | 0.014959 | 0.975320767 |
| 88 | ENSMUSG00000023439 | Gnb3 | protein_coding | 5.770727 | 2.440525 | 1.1239 | 2.171479 | 0.029895 | 0.975320767 |
| 89 | ENSMUSG00000025270 | Alas2 | protein_coding | 8.619067 | 2.442025 | 0.895254 | 2.727746 | 0.006377 | 0.975320767 |
| 90 | ENSMUSG00000023151 | Lrrc69 | protein_coding | 4.49694 | 2.575693 | 1.307098 | 1.970543 | 0.048776 | 0.975320767 |
| 91 | ENSMUSG00000042379 | Esm1 | protein_coding | 6.620143 | 2.57796 | 1.068005 | 2.413809 | 0.015787 | 0.975320767 |
| 92 | ENSMUSG00000070368 | Prok1 | protein_coding | 4.009109 | 2.768193 | 1.299215 | 2.130666 | 0.033117 | 0.975320767 |
| 93 | ENSMUSG00000033770 | Clcnka | protein_coding | 3.219681 | 3.647355 | 1.407757 | 2.590898 | 0.009573 | 0.975320767 |
| 94 | ENSMUSG00000037953 | A4gnt | protein_coding | 2.02107 | 3.867948 | 1.863115 | 2.076066 | 0.037888 | 0.975320767 |
| 95 | ENSMUSG00000096215 | Smim22 | protein_coding | 6.22842 | 3.992469 | 1.125985 | 3.545758 | 0.000391 | 0.537171638 |
| 96 | ENSMUSG00000019987 | Arg1 | protein_coding | 2.621786 | 4.263842 | 1.893322 | 2.252043 | 0.02432 | 0.975320767 |
| 97 | ENSMUSG00000022422 | Dscc1 | protein_coding | 2.491808 | 4.408358 | 2.039652 | 2.161328 | 0.03067 | 0.975320767 |
| 98 | ENSMUSG00000028314 | Toporsl | protein_coding | 2.524323 | 4.445068 | 2.120455 | 2.09628 | 0.036057 | 0.975320767 |
| 99 | ENSMUSG00000037446 | Tulp1 | protein_coding | 3.141857 | 4.73961 | 1.944672 | 2.437228 | 0.0148 | 0.975320767 |
| 100 | ENSMUSG00000064252 | Olfr329-ps | protein_coding | 1.108004 | 5.936079 | 2.943531 | 2.016652 | 0.043732 | 0.975320767 |
| 101 | ENSMUSG00000001155 | Ftcd | protein_coding | 1.134069 | 5.965277 | 2.924348 | 2.039866 | 0.041364 | 0.975320767 |
| 102 | ENSMUSG00000066537 | Vmn2r57 | protein_coding | 1.208298 | 6.062775 | 3.083516 | 1.966189 | 0.049277 | 0.975320767 |
| 103 | ENSMUSG00000069515 | Lyz1 | protein_coding | 1.339722 | 6.211472 | 2.901801 | 2.140557 | 0.03231 | 0.975320767 |
| 104 | ENSMUSG00000025243 | Slc6a20b | protein_coding | 1.34951 | 6.221695 | 2.766061 | 2.249298 | 0.024494 | 0.975320767 |
| 105 | ENSMUSG00000069581 | Tspear | protein_coding | 1.357796 | 6.228009 | 2.982295 | 2.088328 | 0.036768 | 0.975320767 |
| 106 | ENSMUSG00000114755 | Galr3 | protein_coding | 1.381228 | 6.250383 | 2.844074 | 2.197687 | 0.027971 | 0.975320767 |
| 107 | ENSMUSG00000094151 | Gm7233 | protein_coding | 1.445741 | 6.310278 | 3.010566 | 2.096044 | 0.036078 | 0.975320767 |
| 108 | ENSMUSG00000037944 | Ccr7 | protein_coding | 1.480849 | 6.353099 | 2.946906 | 2.155854 | 0.031095 | 0.975320767 |
| 109 | ENSMUSG00000049560 | Defb20 | protein_coding | 1.681609 | 6.540316 | 2.889992 | 2.263091 | 0.02363 | 0.975320767 |
| 110 | ENSMUSG00000032591 | Mst1 | protein_coding | 1.722102 | 6.563315 | 3.092304 | 2.122468 | 0.033798 | 0.975320767 |
| 111 | ENSMUSG00000078597 | Cyp4a12b | protein_coding | 1.992947 | 6.780565 | 2.701943 | 2.509515 | 0.01209 | 0.975320767 |
| 112 | ENSMUSG00000070690 | 5830473C10Rik | protein_coding | 2.330057 | 7.001107 | 2.805657 | 2.495354 | 0.012583 | 0.975320767 |
| 113 | ENSMUSG00000036853 | Mcoln3 | protein_coding | 2.630529 | 7.1801 | 2.6247 | 2.735589 | 0.006227 | 0.975320767 |

**Supplementary Table 7: CCI + PBS vs CCI + TNFR2 Ag TNFR2^F/F^**

| **S.No** | **Gene_ID** | **Gene_Name** | **Gene_Biotype** | **baseMean** | **log2FoldChange** | **lfcSE** | **stat** | **pvalue** | **padj** |
| --- | --- | --- | --- | --- | --- | --- | --- | --- | --- |
| 1 | ENSMUSG00000027261 | Hao1 | protein_coding | 1.408130344 | -6.35740042 | 2.555802833 | -2.487437739 | 0.012866697 | 0.971759875 |
| 2 | ENSMUSG00000004948 | Zp3 | protein_coding | 1.235020793 | -6.17922366 | 2.522203912 | -2.449930249 | 0.014288389 | 0.971759875 |
| 3 | ENSMUSG00000024837 | Dmrt1 | protein_coding | 1.187646837 | -6.104621478 | 2.649791636 | -2.30381189 | 0.021233205 | 0.971759875 |
| 4 | ENSMUSG00000022596 | Slurp1 | protein_coding | 1.137696987 | -6.068527358 | 2.788395916 | -2.176350684 | 0.029529048 | 0.971759875 |
| 5 | ENSMUSG00000030402 | Ppm1n | protein_coding | 1.158345509 | -6.059462252 | 2.800745321 | -2.163517763 | 0.030501379 | 0.971759875 |
| 6 | ENSMUSG00000028786 | Tmem54 | protein_coding | 1.135524076 | -6.030215822 | 2.641049785 | -2.283264729 | 0.022414779 | 0.971759875 |
| 7 | ENSMUSG00000031204 | Asb12 | protein_coding | 1.084095647 | -5.958529256 | 2.879416513 | -2.069353019 | 0.038512972 | 0.971759875 |
| 8 | ENSMUSG00000073610 | Gm10549 | protein_coding | 0.919843426 | -5.74193102 | 2.916064316 | -1.969068717 | 0.048945201 | 0.971759875 |
| 9 | ENSMUSG00000021622 | Ckmt2 | protein_coding | 0.826432937 | -5.602567856 | 2.809550318 | -1.994115507 | 0.046139433 | 0.971759875 |
| 10 | ENSMUSG00000027048 | Abcb11 | protein_coding | 0.760992416 | -5.456918754 | 2.738315994 | -1.992800965 | 0.046283246 | 0.971759875 |
| 11 | ENSMUSG00000056054 | S100a8 | protein_coding | 2.69527525 | -4.769996389 | 1.744423503 | -2.734425661 | 0.00624892 | 0.971759875 |
| 12 | ENSMUSG00000031362 | Xlr4c | protein_coding | 1.957480378 | -4.287828369 | 1.950925675 | -2.197843015 | 0.027960295 | 0.971759875 |
| 13 | ENSMUSG00000047940 | Stpg2 | protein_coding | 1.804892708 | -4.278107107 | 1.953908369 | -2.189512658 | 0.0285596 | 0.971759875 |
| 14 | ENSMUSG00000034774 | Dsg1c | protein_coding | 2.023004781 | -3.418171965 | 1.624088273 | -2.104671293 | 0.035319931 | 0.971759875 |
| 15 | ENSMUSG00000041565 | L3mbtl4 | protein_coding | 2.140934878 | -3.30853266 | 1.594824275 | -2.074543705 | 0.038028844 | 0.971759875 |
| 16 | ENSMUSG00000025091 | Pnliprp2 | protein_coding | 2.820744723 | -2.65090184 | 1.234808558 | -2.146812009 | 0.031808251 | 0.971759875 |
| 17 | ENSMUSG00000075304 | Sp5 | protein_coding | 2.903291912 | -2.620311971 | 1.196871557 | -2.189300895 | 0.028574977 | 0.971759875 |
| 18 | ENSMUSG00000002204 | Napsa | protein_coding | 2.592274991 | -2.443313395 | 1.223069207 | -1.997690222 | 0.045750255 | 0.971759875 |
| 19 | ENSMUSG00000054003 | Tdrd9 | protein_coding | 6.101967851 | -2.361529171 | 0.823744844 | -2.866821186 | 0.004146172 | 0.971759875 |
| 20 | ENSMUSG00000021898 | Asb14 | protein_coding | 3.861618648 | -2.336821052 | 1.016411773 | -2.299088926 | 0.02149989 | 0.971759875 |
| 21 | ENSMUSG00000053441 | Adamts19 | protein_coding | 4.371267988 | -2.222042573 | 0.973884108 | -2.281629359 | 0.022511232 | 0.971759875 |
| 22 | ENSMUSG00000030789 | Itgax | protein_coding | 7.510998976 | -1.911300533 | 0.91335101 | -2.092624317 | 0.036382704 | 0.971759875 |
| 23 | ENSMUSG00000000982 | Ccl3 | protein_coding | 5.423975776 | -1.815624517 | 0.864938186 | -2.099137888 | 0.035804747 | 0.971759875 |
| 24 | ENSMUSG00000058046 | 4933430I17Rik | protein_coding | 11.22728937 | -1.772357937 | 0.613417548 | -2.889317306 | 0.003860793 | 0.971759875 |
| 25 | ENSMUSG00000031849 | Comp | protein_coding | 8.91129037 | -1.73297027 | 0.808012856 | -2.144731059 | 0.031974356 | 0.971759875 |
| 26 | ENSMUSG00000018752 | Tnfsfm13 | protein_coding | 6.370760676 | -1.710524721 | 0.815572641 | -2.097329698 | 0.0359644 | 0.971759875 |
| 27 | ENSMUSG00000110949 | Nudt8 | protein_coding | 9.345628248 | -1.533785282 | 0.679491349 | -2.257255053 | 0.023992142 | 0.971759875 |
| 28 | ENSMUSG00000025983 | Ccdc150 | protein_coding | 7.896623633 | -1.513182162 | 0.755959174 | -2.001671802 | 0.045320041 | 0.971759875 |
| 29 | ENSMUSG00000035296 | Sgcg | protein_coding | 7.520020273 | -1.503453109 | 0.673269041 | -2.233064374 | 0.0255447 | 0.971759875 |
| 30 | ENSMUSG00000025058 | 5430427O19Rik | protein_coding | 6.919349794 | -1.497286405 | 0.754412732 | -1.984704582 | 0.047177349 | 0.971759875 |
| 31 | ENSMUSG00000032380 | Dapk2 | protein_coding | 15.38792798 | -1.462072179 | 0.511733928 | -2.857094476 | 0.004275386 | 0.971759875 |
| 32 | ENSMUSG00000032643 | Fhl3 | protein_coding | 16.49150139 | -1.454886568 | 0.49669297 | -2.929146686 | 0.00339894 | 0.971759875 |
| 33 | ENSMUSG00000038151 | Prdm1 | protein_coding | 17.14040219 | -1.327358432 | 0.451869336 | -2.937482865 | 0.003308885 | 0.971759875 |
| 34 | ENSMUSG00000074635 | 3110070M22Rik | protein_coding | 22.33338916 | -1.310119961 | 0.516188287 | -2.538066039 | 0.011146695 | 0.971759875 |
| 35 | ENSMUSG00000079105 | C7 | protein_coding | 14.38986134 | -1.263270665 | 0.579335315 | -2.180551801 | 0.029216583 | 0.971759875 |
| 36 | ENSMUSG00000026628 | Atf3 | protein_coding | 19.76003986 | -1.238374069 | 0.41866453 | -2.957914944 | 0.003097276 | 0.971759875 |
| 37 | ENSMUSG00000038086 | Hspb2 | protein_coding | 9.98837905 | -1.204508464 | 0.602171422 | -2.00027504 | 0.045470573 | 0.971759875 |
| 38 | ENSMUSG00000038663 | Fsd2 | protein_coding | 13.7797735 | -1.161981853 | 0.53693618 | -2.164096773 | 0.030456923 | 0.971759875 |
| 39 | ENSMUSG00000052631 | Sh2d6 | protein_coding | 13.15353067 | -1.10646511 | 0.511644354 | -2.162566832 | 0.030574512 | 0.971759875 |
| 40 | ENSMUSG00000020787 | P2rx1 | protein_coding | 15.6773476 | -1.095561074 | 0.498884512 | -2.196021418 | 0.02809041 | 0.971759875 |
| 41 | ENSMUSG00000096169 | Olfr1564 | protein_coding | 15.10398516 | -1.069824028 | 0.479982296 | -2.228882266 | 0.025821739 | 0.971759875 |
| 42 | ENSMUSG00000079445 | B3gnt7 | protein_coding | 19.93297781 | -1.032156561 | 0.416614288 | -2.477487189 | 0.013231116 | 0.971759875 |
| 43 | ENSMUSG00000022014 | Epsti1 | protein_coding | 23.59817782 | -1.02604629 | 0.397490551 | -2.581309889 | 0.009842619 | 0.971759875 |
| 44 | ENSMUSG00000036687 | Tmem184a | protein_coding | 29.20590358 | 1.030099673 | 0.492762001 | 2.090460851 | 0.036576422 | 0.971759875 |
| 45 | ENSMUSG00000069899 | Gm12166 | protein_coding | 15.11012772 | 1.06276059 | 0.515501655 | 2.061604614 | 0.039245394 | 0.971759875 |
| 46 | ENSMUSG00000032899 | Styk1 | protein_coding | 13.72755789 | 1.113992268 | 0.499163746 | 2.231717103 | 0.025633666 | 0.971759875 |
| 47 | ENSMUSG00000021815 | Mss51 | protein_coding | 23.01884414 | 1.128597709 | 0.441992974 | 2.553429075 | 0.0106668 | 0.971759875 |
| 48 | ENSMUSG00000002769 | Gnmt | protein_coding | 22.77801917 | 1.13881622 | 0.412192711 | 2.762824744 | 0.005730353 | 0.971759875 |
| 49 | ENSMUSG00000021751 | Acox2 | protein_coding | 16.84448736 | 1.156397756 | 0.517622007 | 2.234058331 | 0.025479235 | 0.971759875 |
| 50 | ENSMUSG00000039748 | Exo1 | protein_coding | 13.33888678 | 1.162353103 | 0.573645363 | 2.026257296 | 0.042738425 | 0.971759875 |
| 51 | ENSMUSG00000043333 | Rhbdl2 | protein_coding | 12.86288359 | 1.171782254 | 0.532902068 | 2.198869783 | 0.027887183 | 0.971759875 |
| 52 | ENSMUSG00000031995 | St14 | protein_coding | 18.07234637 | 1.180104359 | 0.448400818 | 2.631806881 | 0.008493212 | 0.971759875 |
| 53 | ENSMUSG00000053914 | Kdm4d | protein_coding | 39.53487134 | 1.254274429 | 0.325464679 | 3.853795844 | 0.000116301 | 0.246422992 |
| 54 | ENSMUSG00000057068 | Fam47e | protein_coding | 21.40602769 | 1.254452613 | 0.502468291 | 2.496580652 | 0.012539715 | 0.971759875 |
| 55 | ENSMUSG00000063383 | Zfp947 | protein_coding | 27.91253771 | 1.258343459 | 0.389319575 | 3.232160776 | 0.001228579 | 0.763129523 |
| 56 | ENSMUSG00000010830 | Kdelr3 | protein_coding | 17.41468584 | 1.263763495 | 0.485768119 | 2.601577676 | 0.009279605 | 0.971759875 |
| 57 | ENSMUSG00000041301 | Cftr | protein_coding | 11.30512256 | 1.275108862 | 0.605233825 | 2.106803701 | 0.035134599 | 0.971759875 |
| 58 | ENSMUSG00000021281 | Tnfaip2 | protein_coding | 16.34679436 | 1.277112875 | 0.51586642 | 2.475665842 | 0.013298798 | 0.971759875 |
| 59 | ENSMUSG00000027318 | Adam33 | protein_coding | 32.84075814 | 1.287568716 | 0.458444225 | 2.808561319 | 0.00497634 | 0.971759875 |
| 60 | ENSMUSG00000025196 | Cpn1 | protein_coding | 14.45445689 | 1.303125514 | 0.492866153 | 2.643974445 | 0.008193884 | 0.971759875 |
| 61 | ENSMUSG00000018822 | Sfrp5 | protein_coding | 31.95443752 | 1.314665103 | 0.554649562 | 2.370262585 | 0.017775456 | 0.971759875 |
| 62 | ENSMUSG00000058183 | Mmel1 | protein_coding | 18.01010399 | 1.334491698 | 0.495902013 | 2.691039081 | 0.007122985 | 0.971759875 |
| 63 | ENSMUSG00000034387 | Ssu2 | protein_coding | 8.277400108 | 1.364617494 | 0.686536377 | 1.987684177 | 0.04684663 | 0.971759875 |
| 64 | ENSMUSG00000031022 | BC051019 | protein_coding | 8.564408999 | 1.369795339 | 0.688895227 | 1.988394296 | 0.046768099 | 0.971759875 |
| 65 | ENSMUSG00000045394 | Epcam | protein_coding | 25.68431551 | 1.377515212 | 0.588171657 | 2.34202923 | 0.019179213 | 0.971759875 |
| 66 | ENSMUSG00000039092 | Sptlc3 | protein_coding | 15.27791414 | 1.464834693 | 0.647719719 | 2.261525548 | 0.023726732 | 0.971759875 |
| 67 | ENSMUSG00000054435 | Gimap4 | protein_coding | 10.00698076 | 1.466137164 | 0.694850991 | 2.110002263 | 0.034858161 | 0.971759875 |
| 68 | ENSMUSG00000063245 | Zfp993 | protein_coding | 10.85621276 | 1.478068118 | 0.67067023 | 2.203867193 | 0.027533685 | 0.971759875 |
| 69 | ENSMUSG00000031637 | Lrp2bp | protein_coding | 10.7758873 | 1.485964712 | 0.62236707 | 2.38760176 | 0.016958707 | 0.971759875 |
| 70 | ENSMUSG00000044201 | Cdc25c | protein_coding | 16.22859033 | 1.494342037 | 0.504914516 | 2.959594129 | 0.003080446 | 0.971759875 |
| 71 | ENSMUSG00000025064 | Col17a1 | protein_coding | 26.68835754 | 1.535197584 | 0.48263204 | 3.180886179 | 0.001468253 | 0.806382704 |
| 72 | ENSMUSG00000029847 | Slc23a4 | protein_coding | 14.5012512 | 1.541537883 | 0.550156223 | 2.802000264 | 0.005078683 | 0.971759875 |
| 73 | ENSMUSG00000061104 | Sap18b | protein_coding | 12.9837466 | 1.563490781 | 0.55827154 | 2.800591948 | 0.005100897 | 0.971759875 |
| 74 | ENSMUSG00000002588 | Pon1 | protein_coding | 21.40390723 | 1.565764486 | 0.624919552 | 2.505545685 | 0.01222626 | 0.971759875 |
| 75 | ENSMUSG00000025185 | Loxl4 | protein_coding | 7.912529978 | 1.569444168 | 0.793855582 | 1.976989522 | 0.048042815 | 0.971759875 |
| 76 | ENSMUSG00000043088 | Il17re | protein_coding | 11.60113641 | 1.586297013 | 0.789634808 | 2.008899552 | 0.044547787 | 0.971759875 |
| 77 | ENSMUSG00000046223 | Plaur | protein_coding | 5.996552256 | 1.64256293 | 0.773215509 | 2.124327448 | 0.033642772 | 0.971759875 |
| 78 | ENSMUSG00000023393 | Slc17a9 | protein_coding | 9.14783818 | 1.655847159 | 0.734835131 | 2.25335873 | 0.024236539 | 0.971759875 |
| 79 | ENSMUSG00000048015 | Neurod4 | protein_coding | 7.817175098 | 1.682868289 | 0.737183945 | 2.282833613 | 0.022440171 | 0.971759875 |
| 80 | ENSMUSG00000060621 | Nkpd1 | protein_coding | 11.86731738 | 1.686248951 | 0.591916285 | 2.848796347 | 0.004388496 | 0.971759875 |
| 81 | ENSMUSG00000009596 | Taf7l | protein_coding | 6.279996635 | 1.699591364 | 0.815762992 | 2.08343769 | 0.037211346 | 0.971759875 |
| 82 | ENSMUSG00000020295 | Hbq1a | protein_coding | 9.725677987 | 1.74174302 | 0.739800861 | 2.354340352 | 0.018555613 | 0.971759875 |
| 83 | ENSMUSG00000114456 | Hist1h2bh | protein_coding | 7.709253867 | 1.76343548 | 0.792927998 | 2.223954108 | 0.026151532 | 0.971759875 |
| 84 | ENSMUSG00000023943 | Sult1c1 | protein_coding | 8.357468894 | 1.789484114 | 0.792279681 | 2.258652035 | 0.023905038 | 0.971759875 |
| 85 | ENSMUSG00000004939 | Nmrk2 | protein_coding | 6.167274321 | 1.79295751 | 0.879150564 | 2.039420304 | 0.041408098 | 0.971759875 |
| 86 | ENSMUSG00000079343 | C1s2 | protein_coding | 10.96738161 | 1.814470903 | 0.755919684 | 2.400348795 | 0.016379456 | 0.971759875 |
| 87 | ENSMUSG00000038537 | Mc3r | protein_coding | 6.780708765 | 1.821891469 | 0.894761763 | 2.036174929 | 0.041732793 | 0.971759875 |
| 88 | ENSMUSG00000030911 | Zp2 | protein_coding | 8.217737724 | 1.900906519 | 0.8246197 | 2.305191737 | 0.021155837 | 0.971759875 |
| 89 | ENSMUSG00000044092 | C130050O18Rik | protein_coding | 5.905313627 | 1.968108606 | 0.949551549 | 2.072671682 | 0.038202845 | 0.971759875 |
| 90 | ENSMUSG00000033427 | Upb1 | protein_coding | 4.809853947 | 2.054457229 | 0.924186026 | 2.222991011 | 0.026216406 | 0.971759875 |
| 91 | ENSMUSG00000072244 | Trim6 | protein_coding | 5.362484617 | 2.059732011 | 0.850464978 | 2.421889277 | 0.015440054 | 0.971759875 |
| 92 | ENSMUSG00000034452 | Slc24a1 | protein_coding | 8.615335476 | 2.08409417 | 0.857085477 | 2.431605978 | 0.01503205 | 0.971759875 |
| 93 | ENSMUSG00000029032 | Arhgef16 | protein_coding | 16.6567425 | 2.152743267 | 0.593377165 | 3.627950981 | 0.000285679 | 0.425743216 |
| 94 | ENSMUSG00000005892 | Trh | protein_coding | 30.59885868 | 2.250104526 | 0.972179199 | 2.31449565 | 0.02064055 | 0.971759875 |
| 95 | ENSMUSG00000031142 | Cacna1f | protein_coding | 5.935819467 | 2.303805418 | 0.841717243 | 2.737030086 | 0.006199662 | 0.971759875 |
| 96 | ENSMUSG00000024215 | Spdef | protein_coding | 3.692875808 | 2.344050469 | 1.053061291 | 2.225939259 | 0.02601825 | 0.971759875 |
| 97 | ENSMUSG00000041347 | Bdkrb1 | protein_coding | 4.201010982 | 2.358149002 | 1.175749387 | 2.005656161 | 0.044892946 | 0.971759875 |
| 98 | ENSMUSG00000042379 | Esm1 | protein_coding | 5.974534757 | 2.371955382 | 0.88925286 | 2.667357607 | 0.007645028 | 0.971759875 |
| 99 | ENSMUSG00000042118 | Bhmt2 | protein_coding | 12.37096508 | 2.478327482 | 0.645783645 | 3.837705555 | 0.000124189 | 0.246422992 |
| 100 | ENSMUSG00000030236 | Slco1b2 | protein_coding | 4.934381073 | 2.576996699 | 1.215150344 | 2.120722519 | 0.03394516 | 0.971759875 |
| 101 | ENSMUSG00000057439 | Kir3dl2 | protein_coding | 4.041544086 | 2.598640938 | 1.178957979 | 2.204184529 | 0.027511369 | 0.971759875 |
| 102 | ENSMUSG00000050876 | Spata31d1a | protein_coding | 5.116961426 | 2.650725364 | 0.926639933 | 2.860577522 | 0.004228702 | 0.971759875 |
| 103 | ENSMUSG00000069917 | Hba-a2 | protein_coding | 12.82763164 | 2.738073672 | 0.868645126 | 3.152119998 | 0.001620896 | 0.806382704 |
| 104 | ENSMUSG00000039462 | Col10a1 | protein_coding | 3.173418628 | 2.741361403 | 1.371857483 | 1.998284397 | 0.045685836 | 0.971759875 |
| 105 | ENSMUSG00000070683 | Lactbl1 | protein_coding | 2.994481119 | 2.742730116 | 1.361452133 | 2.014562282 | 0.043950533 | 0.971759875 |
| 106 | ENSMUSG00000018907 | Alox12e | protein_coding | 3.084302192 | 2.753625846 | 1.296668759 | 2.123615477 | 0.03370231 | 0.971759875 |
| 107 | ENSMUSG00000038567 | Cyp24a1 | protein_coding | 5.193710212 | 2.829287991 | 0.955918643 | 2.959758148 | 0.003078806 | 0.971759875 |
| 108 | ENSMUSG00000091402 | Rd3l | protein_coding | 6.27441086 | 2.881748531 | 1.2139907 | 2.37378139 | 0.017606971 | 0.971759875 |
| 109 | ENSMUSG00000030771 | Micalcl | protein_coding | 3.001944665 | 2.894921318 | 1.358477289 | 2.131004575 | 0.033088765 | 0.971759875 |
| 110 | ENSMUSG00000009941 | Nxf2 | protein_coding | 3.19533598 | 2.975203189 | 1.397128539 | 2.129512858 | 0.033211852 | 0.971759875 |
| 111 | ENSMUSG00000076438 | Oxct2b | protein_coding | 4.098717586 | 2.977059938 | 1.090966134 | 2.72882892 | 0.006355967 | 0.971759875 |
| 112 | ENSMUSG00000020660 | Pomc | protein_coding | 765.0570345 | 3.005680553 | 1.271679532 | 2.363551883 | 0.018100692 | 0.971759875 |
| 113 | ENSMUSG00000004814 | Ccl24 | protein_coding | 2.494430189 | 3.103342611 | 1.528620511 | 2.03015895 | 0.042340385 | 0.971759875 |
| 114 | ENSMUSG00000028362 | Tnfsf8 | protein_coding | 3.993811725 | 3.228238169 | 1.238016398 | 2.607589184 | 0.009118231 | 0.971759875 |
| 115 | ENSMUSG00000005836 | Gata6 | protein_coding | 2.651879028 | 3.29208428 | 1.544759509 | 2.131130613 | 0.033078383 | 0.971759875 |
| 116 | ENSMUSG00000025270 | Alas2 | protein_coding | 15.31743263 | 3.369386922 | 1.238578514 | 2.720366035 | 0.006520969 | 0.971759875 |
| 117 | ENSMUSG00000096215 | Smim22 | protein_coding | 4.831209858 | 3.537970419 | 1.338180503 | 2.643866363 | 0.008196501 | 0.971759875 |
| 118 | ENSMUSG00000050087 | Cby3 | protein_coding | 1.924208597 | 3.7475358 | 1.833204426 | 2.044254175 | 0.040928444 | 0.971759875 |
| 119 | ENSMUSG00000083193 | 4930595D18Rik | protein_coding | 2.513373413 | 4.157280841 | 1.803045514 | 2.305699334 | 0.021127438 | 0.971759875 |
| 120 | ENSMUSG00000042474 | Fcmr | protein_coding | 2.570921687 | 4.276369159 | 1.940223412 | 2.204060178 | 0.027520112 | 0.971759875 |
| 121 | ENSMUSG00000019987 | Arg1 | protein_coding | 3.940019226 | 4.840995105 | 1.668717807 | 2.901026815 | 0.00371942 | 0.971759875 |
| 122 | ENSMUSG00000031603 | Fgf20 | protein_coding | 1.063830176 | 5.802392451 | 2.796417794 | 2.074937609 | 0.037992318 | 0.971759875 |
| 123 | ENSMUSG00000079681 | Zglp1 | protein_coding | 1.065567331 | 5.828301166 | 2.789595522 | 2.089299728 | 0.036680751 | 0.971759875 |
| 124 | ENSMUSG00000044854 | 1700056E22Rik | protein_coding | 1.086823882 | 5.829310307 | 2.913680092 | 2.000669298 | 0.04542804 | 0.971759875 |
| 125 | ENSMUSG00000037393 | Nmur2 | protein_coding | 1.147534785 | 5.894076158 | 2.799559282 | 2.105358582 | 0.035260106 | 0.971759875 |
| 126 | ENSMUSG00000004651 | Tyr | protein_coding | 1.141051677 | 5.910738759 | 2.880594335 | 2.0519164 | 0.040177788 | 0.971759875 |
| 127 | ENSMUSG00000055561 | Spink5 | protein_coding | 1.217447719 | 5.994007405 | 3.00029127 | 1.997808501 | 0.045737425 | 0.971759875 |
| 128 | ENSMUSG00000073969 | Olfr556 | protein_coding | 1.2454873 | 6.023378443 | 2.992262431 | 2.012984683 | 0.044116241 | 0.971759875 |
| 129 | ENSMUSG00000018924 | Alox15 | protein_coding | 1.250397048 | 6.043327406 | 3.031432714 | 1.993554856 | 0.046200723 | 0.971759875 |
| 130 | ENSMUSG00000025977 | Boll | protein_coding | 1.32009117 | 6.125392866 | 2.837126342 | 2.159013074 | 0.030849153 | 0.971759875 |
| 131 | ENSMUSG00000046408 | 1700067K01Rik | protein_coding | 1.337963008 | 6.128464555 | 2.841556808 | 2.156727797 | 0.03102688 | 0.971759875 |
| 132 | ENSMUSG00000030149 | Klrk1 | protein_coding | 1.360367681 | 6.147703702 | 2.778634411 | 2.212491027 | 0.026932755 | 0.971759875 |
| 133 | ENSMUSG00000073967 | Olfr557 | protein_coding | 1.38626115 | 6.173755032 | 2.833117471 | 2.179138386 | 0.029321389 | 0.971759875 |
| 134 | ENSMUSG00000071716 | Apol7e | protein_coding | 1.426034097 | 6.226162555 | 2.819307406 | 2.208401447 | 0.027216301 | 0.971759875 |
| 135 | ENSMUSG00000026327 | Serpinb11 | protein_coding | 1.755575064 | 6.528992132 | 2.873816944 | 2.271888662 | 0.023093236 | 0.971759875 |
| 136 | ENSMUSG00000008601 | Rab25 | protein_coding | 2.143299783 | 6.813523012 | 2.664558289 | 2.557092874 | 0.010555103 | 0.971759875 |
| 137 | ENSMUSG00000051051 | Olfr523 | protein_coding | 2.369093828 | 6.965327477 | 2.589333588 | 2.690007773 | 0.007145035 | 0.971759875 |
| 138 | ENSMUSG00000026645 | Olah | protein_coding | 2.449036927 | 7.006875306 | 2.585283309 | 2.71029302 | 0.006722379 | 0.971759875 |

**Supplementary Table 8: CCI + PBS vs CCI + TNFR2 Ag Nex-Cre:TNFR2^F/F^**

| **S.No** | **Gene_ID** | **Gene_Name** | **Gene_Biotype** | **baseMean** | **log2FoldChange** | **lfcSE** | **stat** | **pvalue** | **padj** |
| --- | --- | --- | --- | --- | --- | --- | --- | --- | --- |
| 1 | ENSMUSG00000028314 | Toporsl | protein_coding | 2.230063 | -7.15215 | 2.539568 | -2.81629 | 0.004858 | 0.975132591 |
| 2 | ENSMUSG00000027073 | Prg2 | protein_coding | 1.96487 | -6.9712 | 2.66315 | -2.61765 | 0.008854 | 0.975132591 |
| 3 | ENSMUSG00000003484 | Cyp4f18 | protein_coding | 1.654612 | -6.71447 | 2.682421 | -2.50314 | 0.01231 | 0.975132591 |
| 4 | ENSMUSG00000024868 | Dkk1 | protein_coding | 1.417615 | -6.4999 | 2.924016 | -2.22294 | 0.02622 | 0.975132591 |
| 5 | ENSMUSG00000079580 | Tmem217 | protein_coding | 1.404965 | -6.47879 | 2.546148 | -2.54455 | 0.010942 | 0.975132591 |
| 6 | ENSMUSG00000071322 | Tcp10a | protein_coding | 1.332831 | -6.39926 | 2.773142 | -2.30759 | 0.021022 | 0.975132591 |
| 7 | ENSMUSG00000017204 | Gsdma | protein_coding | 1.317558 | -6.38326 | 2.680374 | -2.38148 | 0.017243 | 0.975132591 |
| 8 | ENSMUSG00000025243 | Slc6a20b | protein_coding | 1.238684 | -6.30411 | 2.600687 | -2.42402 | 0.01535 | 0.975132591 |
| 9 | ENSMUSG00000031886 | Ces2e | protein_coding | 1.231255 | -6.29488 | 2.853049 | -2.20637 | 0.027358 | 0.975132591 |
| 10 | ENSMUSG00000050075 | Gpr171 | protein_coding | 1.047277 | -6.05648 | 2.900174 | -2.08831 | 0.036769 | 0.975132591 |
| 11 | ENSMUSG00000091119 | Ccdc152 | protein_coding | 1.035721 | -6.04203 | 2.770118 | -2.18114 | 0.029173 | 0.975132591 |
| 12 | ENSMUSG00000033825 | Tpsb2 | protein_coding | 1.018112 | -6.02001 | 2.780277 | -2.16525 | 0.030368 | 0.975132591 |
| 13 | ENSMUSG00000022832 | Ropn1 | protein_coding | 1.010364 | -6.00943 | 2.958357 | -2.03134 | 0.042221 | 0.975132591 |
| 14 | ENSMUSG00000079516 | Reg3a | protein_coding | 0.920853 | -5.8723 | 2.981909 | -1.96931 | 0.048918 | 0.975132591 |
| 15 | ENSMUSG00000044820 | AY074887 | protein_coding | 0.839641 | -5.73232 | 2.89425 | -1.98059 | 0.047638 | 0.975132591 |
| 16 | ENSMUSG00000034833 | Tespa1 | protein_coding | 0.802495 | -5.6747 | 2.785798 | -2.03701 | 0.041649 | 0.975132591 |
| 17 | ENSMUSG00000034486 | Gbx2 | protein_coding | 3.915967 | -4.53377 | 1.588182 | -2.85469 | 0.004308 | 0.975132591 |
| 18 | ENSMUSG00000078597 | Cyp4a12b | protein_coding | 1.922635 | -4.48554 | 2.042066 | -2.19657 | 0.028051 | 0.975132591 |
| 19 | ENSMUSG00000043230 | Fam124b | protein_coding | 1.838153 | -4.41539 | 2.064659 | -2.13856 | 0.032472 | 0.975132591 |
| 20 | ENSMUSG00000068392 | Rnase13 | protein_coding | 2.024397 | -4.36146 | 1.926622 | -2.26379 | 0.023587 | 0.975132591 |
| 21 | ENSMUSG00000060044 | Tmem26 | protein_coding | 3.12019 | -3.56273 | 1.384344 | -2.57359 | 0.010065 | 0.975132591 |
| 22 | ENSMUSG00000017309 | Cd300lg | protein_coding | 4.295989 | -3.52251 | 1.122672 | -3.13761 | 0.001703 | 0.938548065 |
| 23 | ENSMUSG00000106447 | Gm42957 | protein_coding | 2.029939 | -3.52113 | 1.774358 | -1.98445 | 0.047205 | 0.975132591 |
| 24 | ENSMUSG00000075510 | Fam187a | protein_coding | 3.02428 | -3.052 | 1.343064 | -2.27242 | 0.023061 | 0.975132591 |
| 25 | ENSMUSG00000043948 | Olfr691 | protein_coding | 4.39765 | -2.934 | 1.219485 | -2.40593 | 0.016131 | 0.975132591 |
| 26 | ENSMUSG00000082079 | Dnmt3c | protein_coding | 2.803889 | -2.92867 | 1.364044 | -2.14705 | 0.031789 | 0.975132591 |
| 27 | ENSMUSG00000029597 | Sds | protein_coding | 2.699401 | -2.86432 | 1.342158 | -2.13412 | 0.032833 | 0.975132591 |
| 28 | ENSMUSG00000042189 | Tekt3 | protein_coding | 5.182979 | -2.379 | 1.085668 | -2.19128 | 0.028432 | 0.975132591 |
| 29 | ENSMUSG00000030523 | Trpm1 | protein_coding | 3.094025 | -2.36885 | 1.174456 | -2.01698 | 0.043698 | 0.975132591 |
| 30 | ENSMUSG00000039865 | Slc44a3 | protein_coding | 4.710604 | -2.34714 | 0.914635 | -2.56621 | 0.010282 | 0.975132591 |
| 31 | ENSMUSG00000041534 | Rbp3 | protein_coding | 5.430461 | -2.25599 | 0.914867 | -2.46592 | 0.013666 | 0.975132591 |
| 32 | ENSMUSG00000029491 | Pde6b | protein_coding | 4.046929 | -2.23761 | 1.009609 | -2.21631 | 0.02667 | 0.975132591 |
| 33 | ENSMUSG00000043592 | Unc5cl | protein_coding | 13.50126 | -2.19398 | 0.625108 | -3.50976 | 0.000449 | 0.669671232 |
| 34 | ENSMUSG00000111692 | AC163637.1 | protein_coding | 6.110346 | -2.08827 | 0.863697 | -2.41783 | 0.015613 | 0.975132591 |
| 35 | ENSMUSG00000039699 | Batf2 | protein_coding | 5.343858 | -2.08542 | 0.888715 | -2.34656 | 0.018948 | 0.975132591 |
| 36 | ENSMUSG00000050700 | Emilin3 | protein_coding | 6.181403 | -1.99231 | 0.96749 | -2.05926 | 0.039469 | 0.975132591 |
| 37 | ENSMUSG00000107108 | Gm9936 | protein_coding | 6.855167 | -1.7235 | 0.7809 | -2.20707 | 0.027309 | 0.975132591 |
| 38 | ENSMUSG00000021322 | Aoah | protein_coding | 10.09266 | -1.67432 | 0.851945 | -1.9653 | 0.04938 | 0.975132591 |
| 39 | ENSMUSG00000030981 | Mmp21 | protein_coding | 11.32355 | -1.58195 | 0.600536 | -2.63423 | 0.008433 | 0.975132591 |
| 40 | ENSMUSG00000048534 | Jaml | protein_coding | 6.899221 | -1.57476 | 0.773818 | -2.03505 | 0.041846 | 0.975132591 |
| 41 | ENSMUSG00000006270 | Vax1 | protein_coding | 9.977149 | -1.57366 | 0.639056 | -2.46248 | 0.013798 | 0.975132591 |
| 42 | ENSMUSG00000023903 | Mmp25 | protein_coding | 8.742369 | -1.54842 | 0.772347 | -2.00482 | 0.044982 | 0.975132591 |
| 43 | ENSMUSG00000094083 | Gm1604a | protein_coding | 7.947176 | -1.52345 | 0.645153 | -2.36138 | 0.018207 | 0.975132591 |
| 44 | ENSMUSG00000046500 | Fam19a4 | protein_coding | 11.99237 | -1.39733 | 0.644435 | -2.1683 | 0.030136 | 0.975132591 |
| 45 | ENSMUSG00000074817 | Papolb | protein_coding | 12.87468 | -1.3883 | 0.693673 | -2.00137 | 0.045352 | 0.975132591 |
| 46 | ENSMUSG00000001021 | S100a3 | protein_coding | 10.31333 | -1.31136 | 0.646721 | -2.02771 | 0.04259 | 0.975132591 |
| 47 | ENSMUSG00000035852 | Misp | protein_coding | 13.52172 | -1.29569 | 0.586453 | -2.20937 | 0.027149 | 0.975132591 |
| 48 | ENSMUSG00000037613 | Tnfrsf23 | protein_coding | 36.16457 | -1.21098 | 0.44844 | -2.70043 | 0.006925 | 0.975132591 |
| 49 | ENSMUSG00000026407 | Cacna1s | protein_coding | 21.1416 | -1.09883 | 0.530252 | -2.07228 | 0.038239 | 0.975132591 |
| 50 | ENSMUSG00000043333 | Rhbdl2 | protein_coding | 13.03219 | -1.06347 | 0.524212 | -2.02869 | 0.042489 | 0.975132591 |
| 51 | ENSMUSG00000042622 | Maff | protein_coding | 20.46273 | -1.02737 | 0.471069 | -2.18094 | 0.029188 | 0.975132591 |
| 52 | ENSMUSG00000029410 | Ppef2 | protein_coding | 27.94066 | -1.00038 | 0.492584 | -2.03088 | 0.042267 | 0.975132591 |
| 53 | ENSMUSG00000022385 | Gtse1 | protein_coding | 22.96542 | 1.07215 | 0.477809 | 2.243888 | 0.02484 | 0.975132591 |
| 54 | ENSMUSG00000031727 | Pmfbp1 | protein_coding | 15.61044 | 1.102446 | 0.528501 | 2.085987 | 0.03698 | 0.975132591 |
| 55 | ENSMUSG00000027322 | Siglec1 | protein_coding | 16.76827 | 1.11844 | 0.524909 | 2.130731 | 0.033111 | 0.975132591 |
| 56 | ENSMUSG00000079388 | 2610042L04Rik | protein_coding | 17.90544 | 1.122548 | 0.569909 | 1.969696 | 0.048873 | 0.975132591 |
| 57 | ENSMUSG00000059900 | Tmem40 | protein_coding | 12.14546 | 1.244324 | 0.631484 | 1.970477 | 0.048784 | 0.975132591 |
| 58 | ENSMUSG00000002384 | Bmp8b | protein_coding | 12.15735 | 1.274799 | 0.572524 | 2.22663 | 0.025972 | 0.975132591 |
| 59 | ENSMUSG00000056155 | Nanos3 | protein_coding | 12.22803 | 1.30744 | 0.62964 | 2.076487 | 0.037849 | 0.975132591 |
| 60 | ENSMUSG00000042638 | Gucy2c | protein_coding | 12.49275 | 1.314957 | 0.588038 | 2.236178 | 0.02534 | 0.975132591 |
| 61 | ENSMUSG00000037801 | Iqch | protein_coding | 9.470226 | 1.31618 | 0.661242 | 1.990467 | 0.046539 | 0.975132591 |
| 62 | ENSMUSG00000057606 | Colq | protein_coding | 15.01919 | 1.316607 | 0.601318 | 2.189535 | 0.028558 | 0.975132591 |
| 63 | ENSMUSG00000043286 | Pnpla1 | protein_coding | 9.646309 | 1.322962 | 0.667059 | 1.983277 | 0.047337 | 0.975132591 |
| 64 | ENSMUSG00000024972 | Lgals12 | protein_coding | 14.77974 | 1.326698 | 0.529415 | 2.505968 | 0.012212 | 0.975132591 |
| 65 | ENSMUSG00000032860 | P2ry2 | protein_coding | 27.23916 | 1.361924 | 0.459259 | 2.965483 | 0.003022 | 0.975132591 |
| 66 | ENSMUSG00000036330 | Slc18a1 | protein_coding | 11.57279 | 1.437065 | 0.665387 | 2.159742 | 0.030793 | 0.975132591 |
| 67 | ENSMUSG00000005628 | Tmod4 | protein_coding | 12.82317 | 1.473201 | 0.575969 | 2.557777 | 0.010534 | 0.975132591 |
| 68 | ENSMUSG00000033952 | Aspm | protein_coding | 24.02088 | 1.503656 | 0.511988 | 2.936897 | 0.003315 | 0.975132591 |
| 69 | ENSMUSG00000073063 | Hbq1b | protein_coding | 7.583643 | 1.582389 | 0.747205 | 2.117743 | 0.034197 | 0.975132591 |
| 70 | ENSMUSG00000000359 | Rem1 | protein_coding | 14.49813 | 1.677219 | 0.541619 | 3.096674 | 0.001957 | 0.960924694 |
| 71 | ENSMUSG00000097271 | Gm9903 | protein_coding | 13.09038 | 1.761767 | 0.614917 | 2.865048 | 0.004169 | 0.975132591 |
| 72 | ENSMUSG00000031250 | Tnmd | protein_coding | 7.704063 | 1.817932 | 0.857593 | 2.119807 | 0.034022 | 0.975132591 |
| 73 | ENSMUSG00000048065 | Cyb5r2 | protein_coding | 10.25904 | 1.843861 | 0.867997 | 2.12427 | 0.033648 | 0.975132591 |
| 74 | ENSMUSG00000116121 | Pick1 | protein_coding | 5.861215 | 1.905976 | 0.962019 | 1.981224 | 0.047566 | 0.975132591 |
| 75 | ENSMUSG00000031637 | Lrp2bp | protein_coding | 11.38462 | 1.91036 | 0.678119 | 2.817147 | 0.004845 | 0.975132591 |
| 76 | ENSMUSG00000096546 | Smlr1 | protein_coding | 9.175538 | 2.047302 | 0.734588 | 2.787007 | 0.00532 | 0.975132591 |
| 77 | ENSMUSG00000033182 | Kbtbd12 | protein_coding | 12.78307 | 2.148018 | 0.631111 | 3.403548 | 0.000665 | 0.669671232 |
| 78 | ENSMUSG00000023902 | Zscan10 | protein_coding | 7.787368 | 2.206429 | 0.865944 | 2.548005 | 0.010834 | 0.975132591 |
| 79 | ENSMUSG00000047257 | Prss45 | protein_coding | 8.116243 | 2.246759 | 0.776483 | 2.893507 | 0.00381 | 0.975132591 |
| 80 | ENSMUSG00000038600 | Atp6v0a4 | protein_coding | 9.669607 | 2.381611 | 0.838009 | 2.841988 | 0.004483 | 0.975132591 |
| 81 | ENSMUSG00000020051 | Pah | protein_coding | 6.187861 | 2.38249 | 0.954821 | 2.495221 | 0.012588 | 0.975132591 |
| 82 | ENSMUSG00000009210 | Prr29 | protein_coding | 8.796478 | 2.396265 | 0.758064 | 3.161033 | 0.001572 | 0.908527981 |
| 83 | ENSMUSG00000070708 | Gtsf1l | protein_coding | 4.375307 | 2.398352 | 1.120483 | 2.140462 | 0.032317 | 0.975132591 |
| 84 | ENSMUSG00000026068 | Il18rap | protein_coding | 3.663039 | 2.458922 | 1.167742 | 2.105706 | 0.03523 | 0.975132591 |
| 85 | ENSMUSG00000099913 | Gm28551 | protein_coding | 3.652197 | 2.482314 | 1.255539 | 1.977091 | 0.048031 | 0.975132591 |
| 86 | ENSMUSG00000027398 | Il1b | protein_coding | 4.645094 | 2.507421 | 1.030015 | 2.434355 | 0.014918 | 0.975132591 |
| 87 | ENSMUSG00000079588 | Tmem182 | protein_coding | 4.97204 | 2.648463 | 1.21468 | 2.18038 | 0.029229 | 0.975132591 |
| 88 | ENSMUSG00000044338 | Aplnr | protein_coding | 6.232861 | 2.672436 | 1.039989 | 2.569677 | 0.010179 | 0.975132591 |
| 89 | ENSMUSG00000078487 | Ankrd65 | protein_coding | 4.170177 | 2.730808 | 1.349528 | 2.023527 | 0.043019 | 0.975132591 |
| 90 | ENSMUSG00000021416 | Eci3 | protein_coding | 10.23242 | 2.840923 | 0.977868 | 2.905221 | 0.00367 | 0.975132591 |
| 91 | ENSMUSG00000005952 | Trpv1 | protein_coding | 4.619564 | 2.841549 | 1.225364 | 2.318942 | 0.020398 | 0.975132591 |
| 92 | ENSMUSG00000036858 | Ptcra | protein_coding | 3.492049 | 2.849784 | 1.295286 | 2.20012 | 0.027798 | 0.975132591 |
| 93 | ENSMUSG00000075267 | Pjvk | protein_coding | 2.586809 | 3.035146 | 1.505074 | 2.01661 | 0.043736 | 0.975132591 |
| 94 | ENSMUSG00000092097 | Gm5819 | protein_coding | 2.620071 | 3.076888 | 1.479277 | 2.079995 | 0.037526 | 0.975132591 |
| 95 | ENSMUSG00000090872 | Gm3239 | protein_coding | 2.887965 | 3.254601 | 1.473521 | 2.208724 | 0.027194 | 0.975132591 |
| 96 | ENSMUSG00000100586 | Vmn1r90 | protein_coding | 3.455382 | 3.447404 | 1.447909 | 2.380953 | 0.017268 | 0.975132591 |
| 97 | ENSMUSG00000058773 | Hist1h1b | protein_coding | 3.518358 | 3.523137 | 1.386266 | 2.541458 | 0.011039 | 0.975132591 |
| 98 | ENSMUSG00000089773 | Skint1 | protein_coding | 3.637999 | 3.552051 | 1.447697 | 2.453587 | 0.014144 | 0.975132591 |
| 99 | ENSMUSG00000013643 | Lypd8 | protein_coding | 3.844938 | 3.675914 | 1.403952 | 2.618261 | 0.008838 | 0.975132591 |
| 100 | ENSMUSG00000050578 | Mmp13 | protein_coding | 1.998494 | 3.717806 | 1.894149 | 1.962784 | 0.049671 | 0.975132591 |
| 101 | ENSMUSG00000043760 | Pkhd1 | protein_coding | 2.202299 | 3.835728 | 1.832565 | 2.093092 | 0.036341 | 0.975132591 |
| 102 | ENSMUSG00000040728 | Esrp1 | protein_coding | 2.306453 | 3.903791 | 1.93486 | 2.017609 | 0.043632 | 0.975132591 |
| 103 | ENSMUSG00000024669 | Cd5 | protein_coding | 2.911015 | 4.2477 | 1.951032 | 2.177156 | 0.029469 | 0.975132591 |
| 104 | ENSMUSG00000032315 | Cyp1a1 | protein_coding | 3.013217 | 4.315759 | 1.828671 | 2.360052 | 0.018272 | 0.975132591 |
| 105 | ENSMUSG00000091685 | Gm17359 | protein_coding | 3.378971 | 4.476208 | 1.841621 | 2.430581 | 0.015075 | 0.975132591 |
| 106 | ENSMUSG00000001444 | Tbx21 | protein_coding | 3.470778 | 4.544881 | 1.760319 | 2.581851 | 0.009827 | 0.975132591 |
| 107 | ENSMUSG00000061947 | Serpina10 | protein_coding | 1.138195 | 5.749311 | 2.832218 | 2.029968 | 0.04236 | 0.975132591 |
| 108 | ENSMUSG00000047631 | Apof | protein_coding | 1.168601 | 5.797954 | 2.85228 | 2.032743 | 0.042078 | 0.975132591 |
| 109 | ENSMUSG00000072791 | Abcb5 | protein_coding | 1.227651 | 5.85947 | 2.890936 | 2.026841 | 0.042679 | 0.975132591 |
| 110 | ENSMUSG00000079183 | C030005K15Rik | protein_coding | 1.251368 | 5.869958 | 2.935241 | 1.999821 | 0.04552 | 0.975132591 |
| 111 | ENSMUSG00000040314 | Ctsg | protein_coding | 1.358468 | 5.98046 | 3.04329 | 1.96513 | 0.049399 | 0.975132591 |
| 112 | ENSMUSG00000057465 | Saa2 | protein_coding | 1.387153 | 6.048397 | 2.982107 | 2.028229 | 0.042537 | 0.975132591 |
| 113 | ENSMUSG00000090206 | Tepp | protein_coding | 1.475532 | 6.11932 | 2.960047 | 2.067305 | 0.038705 | 0.975132591 |
| 114 | ENSMUSG00000096629 | Gm3383 | protein_coding | 1.4828 | 6.144096 | 2.950259 | 2.082561 | 0.037291 | 0.975132591 |
| 115 | ENSMUSG00000090843 | Heatr4 | protein_coding | 1.559496 | 6.178231 | 2.944236 | 2.098415 | 0.035868 | 0.975132591 |
| 116 | ENSMUSG00000044222 | Defb13 | protein_coding | 1.561627 | 6.189166 | 2.823189 | 2.192261 | 0.028361 | 0.975132591 |
| 117 | ENSMUSG00000004709 | Cd244 | protein_coding | 1.615185 | 6.215788 | 2.957293 | 2.10185 | 0.035566 | 0.975132591 |
| 118 | ENSMUSG00000022435 | Upk3a | protein_coding | 1.694986 | 6.300764 | 2.861106 | 2.202213 | 0.02765 | 0.975132591 |
| 119 | ENSMUSG00000059430 | Actg2 | protein_coding | 1.697485 | 6.332542 | 2.695074 | 2.349672 | 0.01879 | 0.975132591 |
| 120 | ENSMUSG00000022229 | Atp12a | protein_coding | 2.018505 | 6.576169 | 2.850738 | 2.306831 | 0.021064 | 0.975132591 |
| 121 | ENSMUSG00000021953 | Tdh | protein_coding | 2.310369 | 6.774965 | 2.625953 | 2.580002 | 0.00988 | 0.975132591 |

**Supplementary Table 9:** **CCI + TNFR2 Ag TNFR2^F/F^ vs CCI +TNFR2 Ag Nex-Cre:TNFR2^F/F^**

| **S.No** | **Gene_ID** | **Gene_Name** | **Gene_Biotype** | **baseMean** | **log2FoldChange** | **lfcSE** | **stat** | **pvalue** | **padj** |
| --- | --- | --- | --- | --- | --- | --- | --- | --- | --- |
| 1 | ENSMUSG00000044988 | Ucn3 | protein_coding | 2.953637 | -7.45393 | 2.457155 | -3.03356 | 0.002417 |  |
| 2 | ENSMUSG00000033825 | Tpsb2 | protein_coding | 1.720758 | -6.68382 | 2.627882 | -2.54342 | 0.010977 |  |
| 3 | ENSMUSG00000063935 | Zar1 | protein_coding | 1.66334 | -6.63 | 2.434552 | -2.72329 | 0.006463 |  |
| 4 | ENSMUSG00000028314 | Toporsl | protein_coding | 1.602188 | -6.59759 | 2.654546 | -2.48539 | 0.012941 |  |
| 5 | ENSMUSG00000000724 | Cryba1 | protein_coding | 1.520997 | -6.50601 | 2.483528 | -2.61967 | 0.008802 |  |
| 6 | ENSMUSG00000029378 | Mcub | protein_coding | 1.480213 | -6.45942 | 2.587667 | -2.49623 | 0.012552 |  |
| 7 | ENSMUSG00000039492 | Ccdc27 | protein_coding | 1.43443 | -6.43651 | 2.694339 | -2.3889 | 0.016899 |  |
| 8 | ENSMUSG00000048329 | Mfsd6l | protein_coding | 1.302219 | -6.30795 | 2.572601 | -2.45197 | 0.014208 |  |
| 9 | ENSMUSG00000049719 | Prss46 | protein_coding | 1.294617 | -6.27546 | 2.904961 | -2.16026 | 0.030753 |  |
| 10 | ENSMUSG00000030149 | Klrk1 | protein_coding | 1.268858 | -6.23299 | 2.585929 | -2.41035 | 0.015937 |  |
| 11 | ENSMUSG00000062017 | Abca14 | protein_coding | 1.20794 | -6.18439 | 2.76598 | -2.23588 | 0.02536 |  |
| 12 | ENSMUSG00000003484 | Cyp4f18 | protein_coding | 1.19609 | -6.14605 | 2.791308 | -2.20185 | 0.027676 |  |
| 13 | ENSMUSG00000055561 | Spink5 | protein_coding | 1.133367 | -6.07608 | 2.799566 | -2.17037 | 0.029979 |  |
| 14 | ENSMUSG00000024842 | Cabp4 | protein_coding | 1.113685 | -6.06802 | 2.686171 | -2.25898 | 0.023884 |  |
| 15 | ENSMUSG00000048399 | Tprg | protein_coding | 1.087051 | -6.02267 | 2.832114 | -2.12656 | 0.033456 |  |
| 16 | ENSMUSG00000031886 | Ces2e | protein_coding | 1.058038 | -5.97125 | 2.838233 | -2.10386 | 0.035391 |  |
| 17 | ENSMUSG00000116378 | Gcat | protein_coding | 1.041318 | -5.95074 | 2.841804 | -2.094 | 0.03626 |  |
| 18 | ENSMUSG00000031132 | Cd40lg | protein_coding | 1.016781 | -5.93424 | 2.848171 | -2.08353 | 0.037203 |  |
| 19 | ENSMUSG00000025243 | Slc6a20b | protein_coding | 0.998679 | -5.90938 | 2.856602 | -2.06868 | 0.038577 |  |
| 20 | ENSMUSG00000056706 | Krtap7-1 | protein_coding | 0.991471 | -5.88784 | 2.864984 | -2.05511 | 0.039869 |  |
| 21 | ENSMUSG00000035951 | Ascl3 | protein_coding | 0.953255 | -5.85591 | 2.673474 | -2.19038 | 0.028497 |  |
| 22 | ENSMUSG00000108022 | Gm7298 | protein_coding | 0.948833 | -5.84486 | 2.881856 | -2.02816 | 0.042544 |  |
| 23 | ENSMUSG00000034456 | Uroc1 | protein_coding | 0.934004 | -5.79203 | 2.900325 | -1.99703 | 0.045822 |  |
| 24 | ENSMUSG00000024678 | Ms4a4d | protein_coding | 0.891099 | -5.72902 | 2.682891 | -2.13539 | 0.032729 |  |
| 25 | ENSMUSG00000108763 | Gm36028 | protein_coding | 0.867039 | -5.71763 | 2.710453 | -2.10947 | 0.034904 |  |
| 26 | ENSMUSG00000032257 | Ankk1 | protein_coding | 0.866019 | -5.6947 | 2.831022 | -2.01154 | 0.044269 |  |
| 27 | ENSMUSG00000017204 | Gsdma | protein_coding | 0.809528 | -5.60761 | 2.824358 | -1.98545 | 0.047095 |  |
| 28 | ENSMUSG00000028362 | Tnfsf8 | protein_coding | 3.50309 | -5.27019 | 1.733003 | -3.04107 | 0.002357 |  |
| 29 | ENSMUSG00000031212 | Pgr15l | protein_coding | 2.505696 | -4.80068 | 1.797955 | -2.67008 | 0.007583 |  |
| 30 | ENSMUSG00000026645 | Olah | protein_coding | 2.358256 | -4.70214 | 1.884875 | -2.49467 | 0.012608 |  |
| 31 | ENSMUSG00000083193 | 4930595D18Rik | protein_coding | 2.302208 | -4.67018 | 1.854341 | -2.51851 | 0.011785 |  |
| 32 | ENSMUSG00000029193 | Cckar | protein_coding | 2.033276 | -4.47468 | 1.933961 | -2.31374 | 0.020682 |  |
| 33 | ENSMUSG00000076438 | Oxct2b | protein_coding | 3.66096 | -4.32044 | 1.337521 | -3.23019 | 0.001237 |  |
| 34 | ENSMUSG00000050087 | Cby3 | protein_coding | 1.753509 | -4.27299 | 1.898424 | -2.25081 | 0.024398 |  |
| 35 | ENSMUSG00000052471 | Gm9881 | protein_coding | 1.663782 | -3.98846 | 1.922156 | -2.07499 | 0.037987 |  |
| 36 | ENSMUSG00000038567 | Cyp24a1 | protein_coding | 4.633492 | -3.96007 | 1.117793 | -3.54276 | 0.000396 |  |
| 37 | ENSMUSG00000062007 | Hsh2d | protein_coding | 1.564285 | -3.88914 | 1.872109 | -2.07741 | 0.037763 |  |
| 38 | ENSMUSG00000060044 | Tmem26 | protein_coding | 3.620572 | -3.7068 | 1.312104 | -2.82508 | 0.004727 |  |
| 39 | ENSMUSG00000029811 | Aoc1 | protein_coding | 2.194099 | -3.58293 | 1.572077 | -2.27911 | 0.022661 |  |
| 40 | ENSMUSG00000049409 | Prokr1 | protein_coding | 2.0843 | -3.49985 | 1.716938 | -2.03843 | 0.041507 |  |
| 41 | ENSMUSG00000067006 | Serpinb5 | protein_coding | 2.355342 | -3.44209 | 1.526564 | -2.2548 | 0.024146 |  |
| 42 | ENSMUSG00000054362 | Lexm | protein_coding | 1.947936 | -3.37031 | 1.66364 | -2.02586 | 0.042779 |  |
| 43 | ENSMUSG00000022584 | Ly6c2 | protein_coding | 2.547909 | -3.16334 | 1.374546 | -2.30137 | 0.021371 |  |
| 44 | ENSMUSG00000020660 | Pomc | protein_coding | 727.0298 | -3.04582 | 1.160707 | -2.62411 | 0.008688 | 0.497407141 |
| 45 | ENSMUSG00000037161 | Mgarp | protein_coding | 2.253895 | -2.99735 | 1.46252 | -2.04945 | 0.040419 |  |
| 46 | ENSMUSG00000070504 | Fcrl6 | protein_coding | 2.987975 | -2.52963 | 1.155709 | -2.18881 | 0.02861 |  |
| 47 | ENSMUSG00000005892 | Trh | protein_coding | 28.96557 | -2.39576 | 0.857566 | -2.79367 | 0.005211 | 0.410188476 |
| 48 | ENSMUSG00000054763 | Defb42 | protein_coding | 4.921535 | -2.30013 | 0.993611 | -2.31492 | 0.020617 |  |
| 49 | ENSMUSG00000039865 | Slc44a3 | protein_coding | 4.592218 | -2.25306 | 0.87983 | -2.56079 | 0.010443 |  |
| 50 | ENSMUSG00000048469 | Olfr564 | protein_coding | 4.20025 | -2.06292 | 0.986216 | -2.09175 | 0.036461 |  |
| 51 | ENSMUSG00000045502 | Hcar2 | protein_coding | 4.84935 | -1.90254 | 0.918651 | -2.07102 | 0.038357 |  |
| 52 | ENSMUSG00000048015 | Neurod4 | protein_coding | 7.539119 | -1.81615 | 0.800239 | -2.26952 | 0.023237 |  |
| 53 | ENSMUSG00000001131 | Timp1 | protein_coding | 8.11068 | -1.74181 | 0.846198 | -2.05839 | 0.039552 |  |
| 54 | ENSMUSG00000048534 | Jaml | protein_coding | 7.434577 | -1.7167 | 0.703734 | -2.43942 | 0.014711 |  |
| 55 | ENSMUSG00000049307 | Fut4 | protein_coding | 7.544356 | -1.61848 | 0.745164 | -2.17198 | 0.029857 |  |
| 56 | ENSMUSG00000054435 | Gimap4 | protein_coding | 9.653138 | -1.54361 | 0.777701 | -1.98484 | 0.047162 |  |
| 57 | ENSMUSG00000022676 | Snai2 | protein_coding | 20.59326 | -1.50531 | 0.418514 | -3.59681 | 0.000322 |  |
| 58 | ENSMUSG00000057836 | Xlr3a | protein_coding | 40.75198 | -1.50356 | 0.631039 | -2.38267 | 0.017187 | 0.609617755 |
| 59 | ENSMUSG00000024670 | Cd6 | protein_coding | 9.023017 | -1.39967 | 0.651813 | -2.14735 | 0.031766 |  |
| 60 | ENSMUSG00000017817 | Jph2 | protein_coding | 20.2571 | -1.38826 | 0.50021 | -2.77536 | 0.005514 |  |
| 61 | ENSMUSG00000041431 | Ccnb1 | protein_coding | 9.53285 | -1.36416 | 0.580304 | -2.35077 | 0.018734 |  |
| 62 | ENSMUSG00000024907 | Gal | protein_coding | 16.45477 | -1.3095 | 0.58803 | -2.22693 | 0.025952 |  |
| 63 | ENSMUSG00000114456 | Hist1h2bh | protein_coding | 8.236188 | -1.29431 | 0.638919 | -2.02578 | 0.042787 |  |
| 64 | ENSMUSG00000042045 | Sln | protein_coding | 11.5258 | -1.26928 | 0.629303 | -2.01696 | 0.0437 |  |
| 65 | ENSMUSG00000079652 | Fam71f2 | protein_coding | 16.84496 | -1.25208 | 0.488472 | -2.56326 | 0.010369 |  |
| 66 | ENSMUSG00000025359 | Pmel | protein_coding | 19.28425 | -1.16231 | 0.451762 | -2.57285 | 0.010087 |  |
| 67 | ENSMUSG00000075023 | Accsl | protein_coding | 11.816 | -1.14074 | 0.51821 | -2.2013 | 0.027715 |  |
| 68 | ENSMUSG00000041831 | Sytl3 | protein_coding | 12.01627 | -1.13336 | 0.517566 | -2.18978 | 0.02854 |  |
| 69 | ENSMUSG00000038805 | Six3 | protein_coding | 29.39774 | -1.11923 | 0.493657 | -2.26722 | 0.023377 | 0.660559165 |
| 70 | ENSMUSG00000046168 | Kcnrg | protein_coding | 23.78126 | -1.10283 | 0.415925 | -2.65152 | 0.008013 |  |
| 71 | ENSMUSG00000043333 | Rhbdl2 | protein_coding | 12.94878 | -1.07935 | 0.502393 | -2.14841 | 0.031681 |  |
| 72 | ENSMUSG00000035852 | Misp | protein_coding | 12.03018 | -1.07581 | 0.542505 | -1.98304 | 0.047363 |  |
| 73 | ENSMUSG00000010830 | Kdelr3 | protein_coding | 18.07678 | -1.07016 | 0.479822 | -2.23033 | 0.025726 |  |
| 74 | ENSMUSG00000037613 | Tnfrsf23 | protein_coding | 32.832 | -1.03109 | 0.425153 | -2.42522 | 0.015299 | 0.589739168 |
| 75 | ENSMUSG00000019768 | Esr1 | protein_coding | 299.9069 | 1.020459 | 0.152931 | 6.672683 | 2.51168728468478e-11 | 1.91980817604881e-7 |
| 76 | ENSMUSG00000020437 | Myo1g | protein_coding | 20.22998 | 1.077485 | 0.465722 | 2.313581 | 0.020691 |  |
| 77 | ENSMUSG00000046886 | Zfp474 | protein_coding | 24.57544 | 1.078351 | 0.416454 | 2.589364 | 0.009615 |  |
| 78 | ENSMUSG00000073409 | H2-Q6 | protein_coding | 29.58934 | 1.095031 | 0.414787 | 2.639985 | 0.008291 | 0.495440297 |
| 79 | ENSMUSG00000000359 | Rem1 | protein_coding | 14.39138 | 1.135193 | 0.503685 | 2.253774 | 0.02421 |  |
| 80 | ENSMUSG00000058626 | Capn11 | protein_coding | 12.83133 | 1.1405 | 0.572094 | 1.993553 | 0.046201 |  |
| 81 | ENSMUSG00000007035 | Msh5 | protein_coding | 16.01261 | 1.194286 | 0.572947 | 2.08446 | 0.037118 |  |
| 82 | ENSMUSG00000079677 | Fdx1l | protein_coding | 17.23701 | 1.195033 | 0.569962 | 2.096688 | 0.036021 |  |
| 83 | ENSMUSG00000002500 | Rpl3l | protein_coding | 10.55084 | 1.218759 | 0.553828 | 2.20061 | 0.027764 |  |
| 84 | ENSMUSG00000052485 | Tmem171 | protein_coding | 18.60593 | 1.220741 | 0.450205 | 2.711521 | 0.006698 |  |
| 85 | ENSMUSG00000090124 | Ugt1a7c | protein_coding | 13.37191 | 1.231478 | 0.54191 | 2.272475 | 0.023058 |  |
| 86 | ENSMUSG00000032643 | Fhl3 | protein_coding | 16.25649 | 1.242942 | 0.556886 | 2.231953 | 0.025618 |  |
| 87 | ENSMUSG00000032860 | P2ry2 | protein_coding | 25.10892 | 1.264981 | 0.391937 | 3.227509 | 0.001249 | 0.252150581 |
| 88 | ENSMUSG00000026628 | Atf3 | protein_coding | 21.61812 | 1.26575 | 0.468912 | 2.699336 | 0.006948 |  |
| 89 | ENSMUSG00000048572 | Tmem252 | protein_coding | 15.30996 | 1.305966 | 0.546511 | 2.389643 | 0.016865 |  |
| 90 | ENSMUSG00000056155 | Nanos3 | protein_coding | 11.00527 | 1.330836 | 0.656609 | 2.026831 | 0.04268 |  |
| 91 | ENSMUSG00000006313 | Upk1a | protein_coding | 12.38697 | 1.366948 | 0.601006 | 2.274433 | 0.02294 |  |
| 92 | ENSMUSG00000002384 | Bmp8b | protein_coding | 10.74203 | 1.376823 | 0.562011 | 2.449817 | 0.014293 |  |
| 93 | ENSMUSG00000032769 | Trpa1 | protein_coding | 17.41682 | 1.401726 | 0.621213 | 2.256433 | 0.024044 |  |
| 94 | ENSMUSG00000090707 | Gm8237 | protein_coding | 6.130172 | 1.460197 | 0.74027 | 1.972521 | 0.04855 |  |
| 95 | ENSMUSG00000038151 | Prdm1 | protein_coding | 20.03356 | 1.476014 | 0.402259 | 3.669316 | 0.000243 |  |
| 96 | ENSMUSG00000021213 | Akr1c13 | protein_coding | 6.327335 | 1.512321 | 0.767213 | 1.971188 | 0.048702 |  |
| 97 | ENSMUSG00000043085 | Tmem82 | protein_coding | 11.08545 | 1.531324 | 0.63116 | 2.426204 | 0.015258 |  |
| 98 | ENSMUSG00000074489 | Bglap3 | protein_coding | 32.72484 | 1.59278 | 0.401839 | 3.963724 | 7.38E-05 | 0.059369489 |
| 99 | ENSMUSG00000074635 | 3110070M22Rik | protein_coding | 28.37747 | 1.613209 | 0.402013 | 4.012823 | 6E-05 | 0.055586811 |
| 100 | ENSMUSG00000057457 | Phex | protein_coding | 10.38911 | 1.617744 | 0.675879 | 2.393541 | 0.016687 |  |
| 101 | ENSMUSG00000038403 | Hfe2 | protein_coding | 7.756353 | 1.65381 | 0.793311 | 2.084694 | 0.037097 |  |
| 102 | ENSMUSG00000103800 | Pcdha8 | protein_coding | 9.575452 | 1.658746 | 0.625938 | 2.650018 | 0.008049 |  |
| 103 | ENSMUSG00000024903 | Lao1 | protein_coding | 8.252326 | 1.791777 | 0.830503 | 2.157461 | 0.03097 |  |
| 104 | ENSMUSG00000038600 | Atp6v0a4 | protein_coding | 9.044578 | 1.889706 | 0.702879 | 2.688524 | 0.007177 |  |
| 105 | ENSMUSG00000037600 | Kdf1 | protein_coding | 9.435603 | 1.891012 | 0.709718 | 2.664455 | 0.007711 |  |
| 106 | ENSMUSG00000005640 | Insrr | protein_coding | 4.225291 | 1.953935 | 0.943511 | 2.070919 | 0.038366 |  |
| 107 | ENSMUSG00000030789 | Itgax | protein_coding | 8.773135 | 1.983473 | 0.709947 | 2.793834 | 0.005209 |  |
| 108 | ENSMUSG00000009248 | Ascl2 | protein_coding | 5.506804 | 1.989723 | 0.831853 | 2.391915 | 0.016761 |  |
| 109 | ENSMUSG00000078630 | Tomt | protein_coding | 5.030787 | 2.219169 | 0.995463 | 2.229284 | 0.025795 |  |
| 110 | ENSMUSG00000062939 | Stat4 | protein_coding | 4.654842 | 2.282949 | 1.042775 | 2.189302 | 0.028575 |  |
| 111 | ENSMUSG00000054003 | Tdrd9 | protein_coding | 6.512788 | 2.283139 | 0.856654 | 2.665182 | 0.007695 |  |
| 112 | ENSMUSG00000079588 | Tmem182 | protein_coding | 4.45288 | 2.39363 | 1.101265 | 2.173527 | 0.029741 |  |
| 113 | ENSMUSG00000023829 | Slc22a1 | protein_coding | 2.913508 | 2.473919 | 1.254385 | 1.972217 | 0.048585 |  |
| 114 | ENSMUSG00000006389 | Mpl | protein_coding | 3.761739 | 2.515652 | 1.174479 | 2.141931 | 0.032199 |  |
| 115 | ENSMUSG00000027833 | Shox2 | protein_coding | 7.254562 | 2.65684 | 1.018908 | 2.607535 | 0.00912 |  |
| 116 | ENSMUSG00000069792 | Wfdc17 | protein_coding | 3.96672 | 2.918251 | 1.294253 | 2.254776 | 0.024147 |  |
| 117 | ENSMUSG00000040935 | Padi6 | protein_coding | 4.201135 | 3.14546 | 1.249215 | 2.517949 | 0.011804 |  |
| 118 | ENSMUSG00000036136 | Fam110c | protein_coding | 2.289219 | 3.194842 | 1.495843 | 2.135813 | 0.032695 |  |
| 119 | ENSMUSG00000001504 | Irx2 | protein_coding | 2.441805 | 3.309168 | 1.625392 | 2.035921 | 0.041758 |  |
| 120 | ENSMUSG00000021590 | Spata9 | protein_coding | 2.445686 | 3.335246 | 1.411997 | 2.362078 | 0.018173 |  |
| 121 | ENSMUSG00000074472 | Zfp872 | protein_coding | 2.428052 | 3.395735 | 1.528265 | 2.221954 | 0.026286 |  |
| 122 | ENSMUSG00000047485 | Klhl34 | protein_coding | 2.650139 | 3.419466 | 1.390104 | 2.459863 | 0.013899 |  |
| 123 | ENSMUSG00000027718 | Il21 | protein_coding | 2.696215 | 3.536556 | 1.488592 | 2.375772 | 0.017512 |  |
| 124 | ENSMUSG00000034774 | Dsg1c | protein_coding | 2.479732 | 3.54284 | 1.546375 | 2.291061 | 0.02196 |  |
| 125 | ENSMUSG00000020051 | Pah | protein_coding | 4.876171 | 3.823022 | 1.089099 | 3.510262 | 0.000448 |  |
| 126 | ENSMUSG00000041565 | L3mbtl4 | protein_coding | 3.451917 | 3.874556 | 1.438153 | 2.69412 | 0.007057 |  |
| 127 | ENSMUSG00000078763 | Slfn1 | protein_coding | 1.865352 | 3.944077 | 2.005409 | 1.966719 | 0.049216 |  |
| 128 | ENSMUSG00000022595 | Lypd2 | protein_coding | 1.845849 | 3.983497 | 1.973441 | 2.018554 | 0.043534 |  |
| 129 | ENSMUSG00000109713 | Pvrig | protein_coding | 1.914444 | 4.010419 | 1.875385 | 2.138451 | 0.03248 |  |
| 130 | ENSMUSG00000035932 | Olfr750 | protein_coding | 1.962689 | 4.035717 | 1.981628 | 2.036566 | 0.041694 |  |
| 131 | ENSMUSG00000024391 | Apom | protein_coding | 2.190131 | 4.225967 | 1.978671 | 2.13576 | 0.032699 |  |
| 132 | ENSMUSG00000025127 | Gcgr | protein_coding | 2.204895 | 4.267191 | 1.852128 | 2.303939 | 0.021226 |  |
| 133 | ENSMUSG00000055138 | Gm4861 | protein_coding | 2.511893 | 4.443873 | 1.729292 | 2.569764 | 0.010177 |  |
| 134 | ENSMUSG00000091685 | Gm17359 | protein_coding | 2.893042 | 4.643538 | 1.82106 | 2.549909 | 0.010775 |  |
| 135 | ENSMUSG00000056054 | S100a8 | protein_coding | 3.236349 | 4.804924 | 1.799816 | 2.669675 | 0.007592 |  |
| 136 | ENSMUSG00000026070 | Il18r1 | protein_coding | 3.019452 | 4.805671 | 1.708447 | 2.812888 | 0.00491 |  |
| 137 | ENSMUSG00000059383 | Gfral | protein_coding | 0.907777 | 5.4824 | 2.792588 | 1.963197 | 0.049623 |  |
| 138 | ENSMUSG00000039639 | Kcne1 | protein_coding | 0.938126 | 5.544792 | 2.675343 | 2.072554 | 0.038214 |  |
| 139 | ENSMUSG00000091756 | Gm3095 | protein_coding | 0.960699 | 5.57136 | 2.745506 | 2.029265 | 0.042431 |  |
| 140 | ENSMUSG00000027048 | Abcb11 | protein_coding | 1.013876 | 5.660382 | 2.738193 | 2.067196 | 0.038716 |  |
| 141 | ENSMUSG00000072791 | Abcb5 | protein_coding | 1.04516 | 5.713224 | 2.692317 | 2.122047 | 0.033834 |  |
| 142 | ENSMUSG00000045989 | 4930451I11Rik | protein_coding | 1.060531 | 5.739953 | 2.875477 | 1.996174 | 0.045915 |  |
| 143 | ENSMUSG00000039335 | Spata16 | protein_coding | 1.192549 | 5.914171 | 2.782434 | 2.125538 | 0.033542 |  |
| 144 | ENSMUSG00000032357 | Tinag | protein_coding | 1.251937 | 5.981841 | 2.933047 | 2.039463 | 0.041404 |  |
| 145 | ENSMUSG00000029417 | Cxcl9 | protein_coding | 1.276259 | 5.989718 | 2.506664 | 2.389518 | 0.016871 |  |
| 146 | ENSMUSG00000044222 | Defb13 | protein_coding | 1.329559 | 6.044323 | 2.6172 | 2.309461 | 0.020918 |  |
| 147 | ENSMUSG00000027261 | Hao1 | protein_coding | 1.350151 | 6.096153 | 2.616907 | 2.329526 | 0.019831 |  |
| 148 | ENSMUSG00000028786 | Tmem54 | protein_coding | 1.366877 | 6.114873 | 2.528398 | 2.418477 | 0.015586 |  |
| 149 | ENSMUSG00000072511 | Hhla1 | protein_coding | 1.497367 | 6.236203 | 2.63467 | 2.366977 | 0.017934 |  |
| 150 | ENSMUSG00000074115 | Saa1 | protein_coding | 2.397564 | 6.902256 | 2.379417 | 2.900818 | 0.003722 |  |
